# Naked antisense oligonucleotides access the cytosol during early endocytic trafficking

**DOI:** 10.64898/2026.05.11.724403

**Authors:** Ewa Sitarska, Anand Saminathan, Gustavo Scanavachi, Elliott Somerville, Patrick Stock, Pengxiang Shen, Daniel Kahne, Margo F. Courtney, Dylan A. Reid, Mathias Bogetoft Danielsen, Fie Kristine Noergaard Davidsen, Knud J. Jensen, C. Frank Bennett, Tom Kirchhausen

**Affiliations:** Program in Cellular and Molecular Medicine, Boston Children’s Hospital, 200 Longwood Ave, Boston, MA 02115, USA; Department of Cell Biology, Harvard Medical School, Boston Children’s Hospital, 200 Longwood Ave, Boston, MA 02115, USA; Department of Pediatrics, Harvard Medical School, Boston Children’s Hospital, 200 Longwood Ave, Boston, MA 02115, USA; Department of Chemistry and Chemical Biology, Harvard University, 12 Oxford Street, Cambridge, MA 02138; Ionis Pharmaceuticals, Carlsbad, CA 92010, USA; Department of Chemistry & Nanoscience Center, University of Copenhagen, Copenhagen, Denmark; Novo Nordisk Center for Optimized Oligo Escape, Copenhagen, Denmark

**Author notes:** Corresponding author: Tom Kirchhausen. Ph.D. Harvard Medical School 200 Longwood Ave Boston, MA 02115 /. Ewa Sitarska and Anand Saminathan contributed equally to this work.

**Keywords:** Antisense oligonucleotides, endosomal escape, early endosomes, endolysosomal sequestration, membrane damage

## Abstract

Phosphorothioate-modified antisense oligonucleotides (ASOs) enter cells, but their site of productive escape remains uncertain. We combined live-cell imaging and ratiometric pH measurements to follow a *Malat1*-targeting ASO internalized without a delivery vehicle. In U2OS cells, less than 4% of internalized ASO reached the nucleus under conditions that reduced *Malat1* RNA by >90%. Most ASO accumulated in late endosomes, endolysosomes and lysosomes as foci and broadly distributed luminal signal. These compartments underwent neutralization without detectable membrane rupture or galectin-3 recruitment. ASO foci showed constrained motion, consistent with association with endolysosomal constituents. L-leucyl-L-leucine methyl ester-induced membrane damage or Ca²⁺-dependent lysosome exocytosis released dextran while retaining luminal ASO. Loss of the stress-granule scaffolding proteins G3BP1/2 modestly enhanced ASO activity without detectable endolysosomal damage. Tantalosin, a small molecule that perturbs endosomal recycling, redistributed the ESCRT-III protein IST1 to endosomes. It enhanced early escape of SYTOX Green and ASOs, detected by nuclear accumulation, and increased *Malat1* knockdown, with cell-type-dependent responses. Tantalosin did not detectably increase nuclear ASO accumulation after sequestration in late compartments. These findings place productive ASO escape early in endocytic trafficking, most likely in early or recycling endosomes, before sequestration in largely nonproductive late compartments.

**GRAPHICAL ABSTRACT:** 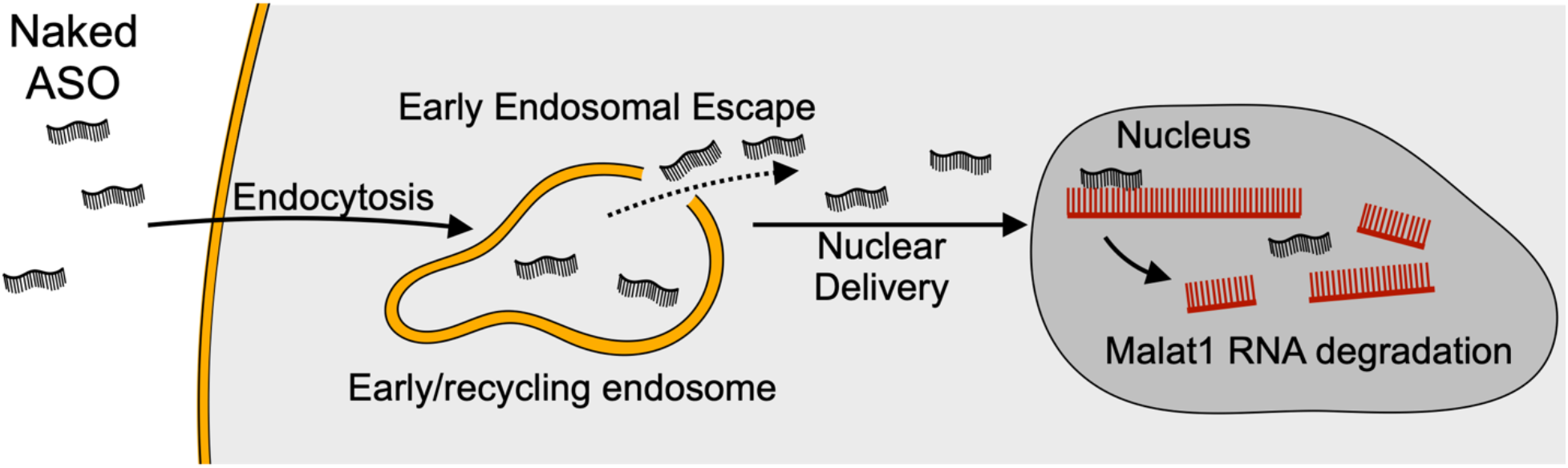

## INTRODUCTION

Antisense oligonucleotides (ASOs) are short, chemically modified nucleic acids that bind complementary RNA and alter its fate (1). They can reduce RNA abundance, often through recruitment of RNase H1, or modulate pre-mRNA splicing by blocking access of splicing factors (2, 3). These properties have made ASOs useful both as experimental tools and as therapeutics, particularly for targets not readily addressed by small molecules or antibodies.

For an ASO to act, it must enter a cell and reach its RNA target in the cytosol or nucleus. Current evidence indicates that endocytosis is the principal route of entry (4, 5). Internalized ASOs traffic through the endosomal system and must cross through a membrane at some stage along the endocytic pathway to gain access to the cytosol, from which they can diffuse or be transported to sites of action in the cytosol or nucleus (4, 6). Membrane crossing and escape from the endosome is an essential and potentially rate limiting step to achieve productive activity, but its mechanism remains poorly understood (7).

Several strategies have been developed to improve delivery. One packages ASOs in carriers, including lipid nanoparticles and related formulations, that promote uptake and are thought to facilitate release after internalization, although the underlying mechanism remains uncertain (8–10). Another links ASOs covalently to peptides, aptamers, lipids, or other targeting groups to enhance binding to selected cell types and increase uptake (11–14). Such conjugates can improve delivery to specific cell-and tissue-types, but endosomal escape is likewise unclear. Unconjugated, or “naked,” phosphorothioate modified (PS) and 2’-modified ASOs are also widely used, including in the clinic (15). The PS modification enhances binding to plasma and cell-surface proteins and increases resistance to nuclease degradation (16). Ribose 2′ modifications increase affinity for the target RNA and can further enhance nuclease resistance; some, including 2′-MOE, also reduce proinflammatory effects and improve tolerability (1, 3). ASOs enter cells without transfection reagents by a process termed gymnosis (17), generally thought to depend on one or more endocytic pathways. Here too, the critical unresolved step is escape from intracellular vesicles (17, 18).

Chemically stabilized PS-ASOs can modify target RNA for weeks to months after administration. For example, a 2′-MOE PS ASO directed against SMN2 produced sustained splicing correction for six months after treatment ended in mice, in parallel with slow clearance from CNS tissue (19). The intracellular basis for such prolonged activity remains uncertain. ASOs that escape during the initial exposure might persist in the cytosol and nucleus, whereas ASOs retained in endolysosomal compartments might provide a delayed source through gradual release or through exocytosis, re-uptake, and escape during a subsequent passage through the endocytic pathway. Altering endolysosomal trafficking can enhance ASO activity: the sphingolipid analog SH-BC-893 increased ASO accumulation in extra-lysosomal compartments and enhanced ASO activity in cultured cells and in mice, leading the authors to propose that escape occurs before lysosomal delivery (20). The compartment from which ASOs cross the limiting membrane nevertheless remains unresolved.

Free uptake of phosphorothioate ASOs can produce measurable biological activity, but only a small fraction of internalized molecules reaches the cytosol and nucleus (21). Most enter through endocytic pathways and progress through the canonical endosomal system (4). The principal barrier is therefore not uptake, but release from membrane-bound compartments. A common assumption has been that ASOs that fail to escape are delivered to lysosomes for degradation - a view that is incompletely characterized (22, 23). Chemically stabilized ASOs persist for long periods, and lysosomal accumulation more likely reflects sequestration rather than degradation. For ASOs, lysosomes appear to function primarily as terminal storage compartments that render the molecules biologically unavailable (24).

Although many studies identified factors that enhance ASO activity, including perturbations of endosomal maturation, retrograde trafficking, lipid composition, and endolysosomal fusion they do not establish how escape occurs (25–28). In most cases, the data support an indirect interpretation, that altered trafficking changes the residence time or routing of internalized ASOs and thereby changes the probability that a small escaping fraction is retained in pre-lysosomal compartments. While such results do not demonstrate a specific release mechanism, they are consistent with redistribution of cargo away from terminal sequestration, with presumed retention in earlier endosomal intermediates.

A further difficulty is that both ensemble measurements and time-lapse imaging reported so far are poorly suited to detect sporadic or infrequent catastrophic release events. Bulk colocalization, endpoint activity assays, population-averaged perturbations, and conventional live-cell imaging cannot readily distinguish true membrane translocation from delayed progression through the endolysosomal pathway, nor can they exclude rare rupture events that involve only a very small subset of compartments. This limitation is especially important for phosphorothioate-modified ASOs, whose chemical stabilization was introduced to protect them from degradation; these molecules can therefore persist and accumulate in lysosomes rather than be rapidly destroyed. Their long intracellular lifetime strengthens the conclusion that productive escape is a rare event, because most internalized ASOs appear to survive sequestration without gaining access to the cytosol. Mechanistic inference therefore requires direct observation of individual organelles and of cargo movement relative to membrane integrity.

We therefore used single-organelle imaging in living cells to follow fluorescently conjugated ASOs, soluble luminal markers, and nuclear ASO accumulation. We found that internalized ASOs remained associated with endosomes and lysosomes and that only about 4% of the internalized material reached the nucleus; any residual cytosolic pool was below detection. Moreover, we found no evidence for catastrophic release of ASOs from late endosomes or lysosomes, even when their limiting membranes were sufficiently compromised to release similarly sized dextran. Overt membrane damage in late compartments is therefore insufficient to account for productive ASO delivery.

These observations shifted our attention to the early-to-recycling endosomal pathway. Cargo sorting and recycling in these compartments entail repeated membrane tubulation, constriction, and scission (29–31). IST1, an ESCRT-III protein, participates in this membrane remodeling (30, 32), and the small molecule tantalosin has been reported to disrupt IST1-dependent recycling (33). Because ESCRT factors also repair damaged endolysosomal membranes (34–36), we asked whether IST1 helps close transient membrane discontinuities that might arise during tubule remodeling. We reasoned that pharmacological interference with this activity could prolong such openings and permit luminal cargo to reach the cytosol.

Consistent with this model, tantalosin increased the early luminal escape of internalized SYTOX Green, a membrane-impermeant fluorogenic dye whose nuclear fluorescence increases many folds upon DNA binding (37). Tantalosin also increased nuclear accumulation of fluorescent ASOs and enhanced ASO-mediated depletion of *Malat1* RNA. These effects occurred early after uptake, before substantial accumulation of either cargo in late endosomes or lysosomes and were not reproduced when the cargoes were first delivered to terminal lysosomes.

We also examined whether G3BP1/2 influences ASO delivery because previous work in macrophages suggested that these proteins are recruited to sites of acute endolysosomal damage and contribute to membrane repair (38). Depletion of G3BP1/2 enhanced ASO activity, consistent with an independent study published while our work was in progress (39). We nevertheless detected neither G3BP1/2 recruitment to overtly damaged endosomes nor an effect of pharmacological interference with G3BP1/2 function on ASO activity. Together, our observations place productive ASO release at an early stage of endocytic trafficking rather than in late endosomes or lysosomes. By defining both the timing of escape and the compartments from which overt membrane damage fails to release ASOs, they narrow the search for the underlying mechanism.

## MATERIALS AND METHODS

### Reagents

The following reagents were used: Apilimod (HY-14644, MedChemExpress), Alexa Fluor 647-dextran 10 KDa (D22914, Thermo Scientific), Alexa Fluor 568-Dextran 10 KDa (D22912, Thermo Scientific), pHrodo Green Dextran (P35368, Thermo Fisher Scientific), Alexa Fluor 647 human transferrin (T23366, Invitrogen), Calcein-AM (22002, AAT Bioquest), hIST1 monoclonal antibody (66989-1-IG, Proteintech), L-Leucyl-L-Leucine methyl ester (16008, Cayman Chemical Company), Sodium arsenite (S7400, Sigma-Aldrich), LysoTracker^TM^ Red DND-99 (L7528, Thermo Fisher Scientific), ISRIB (HY-12495A, MedChem Express), FAZ-3532 (G3Ia, HY-162288, MedChem Express), Milli-Q water (Z00QSV0US, MilliporeSigma), Minimum Essential Medium (MEM; 10-010-CV, Corning), Dulbecco’s Modified Eagle Medium (DMEM; 10-013-CV, Corning), fetal bovine serum (S11150H, Atlanta Biologicals), FluoroBrite DMEM (A1896701, Gibco), DMEM/F-12 (Gibco 11320-033), HEPES buffer pH 7.2 (SH30237.01, Cytiva), Guanidine isothiocyanate (V2791, Promega), Nigericin (11437, Cayman Chemicals), Monensin (16488, Cayman Chemicals), SYTOX^TM^ Green Nucleic Acid Stain (S7020, Invitrogen), Ionomycin (I242222, Invitrogen), Geneticin (10131-035, Thermo Fisher Scientific), Hydrocortisone (H4001, Sigma), Insulin (I1882, Sigma), BioReagent Penicillin-Streptomycin, Sterile, 100X (P4333-100ML, Sigma-Aldrich), Hoechst 33342 (H1399, ThermoFisher Scientific), Lipofectamine 2000 Transfection Reagent (Invitrogen, 11668030), propylene oxide (14121, Electron Microscopy Sciences (EMS)), Janelia Fluor 549 HaloTag ligand (JF549-HTL), Hoechst Janelia Fluor 549 (JF549-HTL and Hoechst JF549, gift from Luke Lavis lab, Janelia Research Campus, HHMI). Janelia Fluor 646 azide (TOCRIS).

### Plasmids and antisense oligonucleotides

The plasmid pEGFP-C1-G3BP1-WT was obtained from Dr. Anthony Leung (Addgene plasmid 135997), and pEGFP-hGal3 was obtained from Dr. Tamotsu Yoshimori (Addgene plasmid 73080). The MALAT1 antisense oligonucleotide (ASO) was a 5-10-5 gapmer with sequence **CCAGG**<u>C</u>TGGTTATGA**CTCAG**. Bold characters indicate RNA residues bearing 2′-O-methoxyethyl (MOE) modifications, and the underlined base indicates 5-methylcytosine. All nucleotides were linked by phosphorothioate linkages. Fluorescently labeled ASOs had the same sequence and were conjugated at the 5′ end to Alexa Fluor 488 or Alexa Fluor 647 through a amino C6 linker. Fluorescent dye conjugation did not alter ASO activity (24). ASOs were purchased from Integrated DNA Technologies (IDT).

### Cell culture

U2OS wild-type and U2OS G3BP1/2-knockout cells (double knockout of G3BP1 and G3BP2) were kindly provided by the laboratory of Dr. P. Anderson (40). Cells were maintained in DMEM containing 10% fetal bovine serum (FBS) and 1% penicillin-streptomycin at 37°C in 5% CO₂. SVGA cells were obtained from ATCC (CRL-8621) and cultured in MEM supplemented with 10% FBS and 1% penicillin-streptomycin. SUM159 human breast carcinoma cells were provided by Dr. J. Brugge (41) and cultured in DMEM/F-12 supplemented with 5% FBS, 1% penicillin-streptomycin, 1 μg/ml hydrocortisone, 5 μg/ml insulin and 10 mM HEPES, pH 7.2. All cell lines were routinely tested for mycoplasma contamination by PCR using a published protocol (42) and confirmed to be negative.

### Establishment of stably expressing cell lines

U2OS wild-type and G3BP1/2 knockout cells were transfected with pEGFP-C1-G3BP1-WT or pEGFP-hGal3 using Lipofectamine 2000 according to the manufacturer’s instructions. Cells were seeded in 12-well plates (Celltreat, 229111) 1 day before transfection to reach 70-90% confluency at the time of transfection. The medium was replaced 6-hours after transfection, and stable transfectants were selected for 2 weeks with 600 μg/ml Geneticin. eGFP-positive cells were then sorted on a Sony SH-800Z cell sorter at the BCH FICR-PCMM Flow and Imaging Cytometry Facility to obtain matched expression levels in U2OS wild-type and G3BP1/2-knockout cells. Cell stocks were frozen and stored in liquid nitrogen.

We generated heterozygous IST1–Halo SVG-A cells by CRISPR–Cas9 genome editing as described previously (43, 44). HaloTag (45) was inserted at the endogenous IST1 C terminus, preceded by a (GGS)₃ linker and followed by the stop codon. Parental SVG-A cells were seeded in 12-well plates and transfected the next day at approximately 80% confluence with 0.25 μg each of the pUC19-based IST1–Halo repair-template plasmid and pX459 encoding the single-guide RNA and *Streptococcus pyogenes* Cas9. Transfection used 1.5 μl of Lipofectamine 2000 (Invitrogen) per well, following the manufacturer’s protocol. The guide-cloning oligonucleotides were 5′-CACCGAAACATAGGTCTCTTAAACC-3′ (forward) and 5′-AAACGGTTTAAGAGACCTATGTTTC-3′ (reverse).

We replaced the medium 6 h after transfection and selected cells with 10 μg/ml puromycin for 3 days. Single cells were then seeded in 96-well plates, and the resulting colonies were manually picked and expanded. We screened clones for Halotag expression by PCR of genomic DNA using set of primers amplifying the inserted tag (forward 5’-CGGAGGTTTGAAGAGCTGAA-3’, reverse 5’-CTCAGGATCAACAGAGAACTGG-3’). IST1–Halo expression was confirmed by immunoblotting for hIST1 and HaloTag and by fluorescence imaging after HaloTag labeling. We identified one heterozygous clone carrying a single edited IST1-Halo allele.

### Quantitative PCR

Cells were lysed in 50 μl per well of buffer containing 4 M guanidine isothiocyanate, 50 mM Tris-HCl (pH 7.5) and 25 mM EDTA. RNA was precipitated with 50 μl per well of 70% ethanol and further purified with AcroPrep Advance filter plates (Pall). Real-time RT-PCR used AgPath-ID One-Step RT-PCR reagents (Applied Biosystems) carried with a QuantStudio 7 Flex Sequence Detection System (Applied Biosystems). Each 10 μl reaction contained 1 μl RNA. For MALAT1 (assay ID RTS2738), the sequences were forward, 5′-GAATTGCGTCATTTAAAGCCTAGTT-3′; reverse, 5′-TCATCCTACCACTCCCAATTAATCT-3′; probe, 5’-FAM-ACGCATTTACTAAACGCAGACGAAAATGGA-BHQ1-3’. For cyclophilin (assay ID 14541), the sequences were forward, 5′-TGCTGGACCCAACACAAATG-3′; reverse, 5′-TGCCATCCAACCACTCAGTC-3′; probe, 5’-FAM-TTCCCAGTTTTTCATCTGCACTGCCAX-BHQ1-3’. Reactions were run in triplicate. MALAT1 values were normalized first to cyclophilin mRNA and then to control samples and are reported as percent of a control. Cells were incubated with the indicated amounts of ASO for 24 h.

### Endolysosomal colocalization and trafficking assays

U2OS wild-type and G3BP1/2-knockout cells were seeded at 40% confluence in 8-well glass-bottom chambers and cultured for 24-hours at 37°C in 5% CO₂. For colocalization experiments, cells were incubated with 8 μM Alexa Fluor 568-dextran and 1 μM Alexa Fluor 647-ASO for 2-hours in complete DMEM containing 10% FBS and 1% penicillin-streptomycin, washed three times with complete medium and imaged by spinning-disk confocal microscopy.

For trafficking assays, cells were loaded with 8 μM Alexa Fluor 568–dextran for 4 h, washed three times, and chased for 16 h to label terminal lysosomal compartments (46). Dextran-labeled cells were then incubated with 1 μM Alexa Fluor 647–ASO or 8 μM Alexa Fluor 647–dextran for 15 min in complete medium, washed three times, and imaged for 2 h by time-lapse lattice light-sheet microscopy (Zeiss) in complete medium containing 10% FBS and 1% penicillin–streptomycin, as described under Fluorescence imaging.

Transferrin uptake and recycling assays were performed in SVG-A, U2OS, and SUM159 cells in the absence or presence of 5 μM tantalosin. Cells were first incubated for 1 h in starvation medium, corresponding to the culture medium used for each cell type lacking FBS. Where indicated, 5 μM tantalosin was then added for an additional 30 min in the same starvation medium. Cells were next incubated for 10 min at 37 °C with 5 μg/ml Alexa Fluor 647–labeled holo-transferrin in the same medium, with or without tantalosin. Cells were then washed three times with complete medium containing 10% FBS and 1% penicillin–streptomycin, with or without 5 μM tantalosin as appropriate, chased for 10 min in the same complete medium, and imaged live for 60 min.

### Endolysosomal pH by imaging

*(i) Dextran-based assay.* To assess the effect of ASOs on endolysosomal membrane damage, pH in late endosomes and endolysosomes was estimated with 8 μM pHrodo Green-dextran and 8 μM Alexa Fluor 647-dextran, the latter serving as a pH-insensitive normalization marker. Cells were co-incubated with the dextran mixture in the presence or absence of 1 μM non-fluorescent ASO for 2-hours at 37°C, washed three times, and transferred to phenol red-free FluoroBrite medium supplemented with 5% FBS and 1% penicillin-streptomycin for imaging at 37°C in 5% CO₂. For calibration at pH 7.4, cells were incubated for 15 min in universal buffer containing 10 mM HEPES, 10 mM MES, 10 mM sodium acetate, 10 mM EDTA, 140 mM KCl, 5 mM NaCl and 1 mM MgCl₂, adjusted to pH 7.4, together with 10 μM nigericin and 10 μM monensin (47).
*(ii) Lysotracker-based assay*. To assess the effect of G3BP1/2 knockout on lysosomal membrane damage, U2OS wild-type and G3BP1/2-knockout cells were preloaded with 4 μM Alexa Fluor 647-dextran and then incubated with 100 nM LysoTracker^TM^ Red for 10 min at 37°C in 5% CO₂ in complete DMEM supplemented with 10% FBS and 1% penicillin-streptomycin. LysoTracker^TM^ Red fluorescence served as a readout for lysosomal acidification. After three washes, cells were chased for 10 min in complete medium and imaged by lattice light-sheet microscopy (Zeiss). As positive controls for pH neutralization, cells were treated before the LysoTracker^TM^ Red pulse with 100 nM bafilomycin A1 for 30 min or with pH 7.4 universal buffer containing 10 μM nigericin plus 10 μM monensin.

### Endosomal dye leakage assay

Cells were incubated for 2-hours at 37°C with a 1:1 mixture of 8 μM Alexa Fluor 568 carboxylic acid and Alexa Fluor 647-dextran (10 kDa), in the presence or absence of 1 μM non-fluorescent ASO. Cells were then washed three times and imaged at 37°C in complete medium (DMEM supplemented with 10% FBS and 1% penicillin-streptomycin). As a positive control for lysosomal damage, labeled cells were treated with 0.5 mM LLOMe for 30 min before imaging. We quantified dye leakage from individual endosomes from the Alexa Fluor 568 signal relative to retained Alexa Fluor 647-dextran. Note: as LLOMe promotes dextran escape, Alexa Fluor 568: Alexa Fluor 647-dextran ratio were only computed on compartments positive for Alexa Fluor 647-dextran.

### LLOMe-mediated release assay

Endolysosomal membrane permeability to macromolecules was assessed by inducing membrane damage with LLOMe. U2OS wild-type cells stably expressing galectin-3-eGFP were pulsed with 8 μM Alexa Fluor 568-dextran and 1 μM Alexa Fluor 647-ASO for 3-hours and then chased for 16-hours to label lysosomal compartments. Cells were imaged in complete DMEM supplemented with 10% FBS and 1% penicillin-streptomycin by spinning-disk confocal microscopy or Zeiss lattice light-sheet microscopy. LLOMe was added to a final concentration of 0.5 mM during imaging, and time-lapse images were acquired for 30 min to monitor release of dextran and ASO.

To examine late endosomal compartments, we first labeled terminal lysosomes by pulsing cells with Alexa Fluor 488-ASO for 3-hours followed by a 16-hours chase. Cells were then incubated with Alexa Fluor 647-ASO and Alexa Fluor 568-dextran for 2.5-hours to label the endolysosomal population. LLOMe was added to a final concentration of 0.5 mM during imaging for 30 min, and we assessed compartment permeability from the release of dextran and ASO. Triple-labeled cells were imaged by spinning-disk confocal microscopy in complete DMEM supplemented with 10% FBS and 1% penicillin-streptomycin.

### Lysosome exocytosis assay

U2OS cells were co-incubated for 2 h at 37 °C with 4 μM Alexa Fluor 568–dextran and 1 μM Alexa Fluor 647–labeled ASO in DMEM supplemented with 10% FBS and 1% penicillin–streptomycin. Cells were then washed three times and chased overnight to allow both probes to accumulate in terminal lysosomes. After the chase, cells were treated for 10 min at 37 °C with Versene, an EDTA-containing, nonenzymatic dissociation reagent that chelates Ca²⁺ and Mg²⁺ and weakens cation-dependent cell adhesion without proteolysis of cell-surface proteins. Cells were then replated and allowed to adhere for 12–36 h. This step reduced background ASO fluorescence from nonspecific binding to the coverslip, which otherwise obscured signal at the cell–coverslip interface.

Before imaging, cells were washed and the medium was replaced with FluoroBrite DMEM supplemented with 2 mM CaCl₂ and 1% penicillin–streptomycin, without FBS. Ten fields of view were selected, and baseline 3D images were acquired by live-cell spinning-disk confocal microscopy. Lysosome exocytosis was then triggered by addition of 2 μM ionomycin, and 3D time-lapse images were acquired at 1-min intervals for 16 min by spinning-disk confocal microscopy, as described under Imaging methods.

### Galectin-3 imaging

U2OS wild-type and G3BP1/2-knockout cells stably expressing eGFP-galectin-3 were seeded at 60% confluency in 8-well glass-bottom chambers and imaged the following day. Cells were incubated for 2-hours at 37°C in 5% CO_2_ with 8 μM Alexa Fluor 568-dextran, with or without 1 μM Alexa Fluor 647-ASO, in complete DMEM supplemented with 10% FBS and 1% penicillin-streptomycin. After three washes with complete medium, cells were imaged by spinning-disk microscopy or Zeiss lattice light-sheet microscopy to monitor galectin-3 recruitment. For a positive control of induced membrane damage, cells were incubated for 2-hours with 8 μM Alexa Fluor 568-dextran and then treated with 0.5 mM LLOMe for 30 min in complete medium before imaging.

### G3BP1-eGFP recruitment assay

G3BP1/2-knockout cells stably expressing G3BP1-eGFP were seeded at 60% confluence in 8-well glass-bottom chambers and imaged the next day. Cells were incubated with 8 μM Alexa Fluor 568-dextran with or without 1 μM Alexa Fluor 647-ASO for 2-hours in complete DMEM containing 10% FBS and 1% penicillin-streptomycin at 37°C in 5% CO₂. After three washes, cells were imaged by spinning-disk confocal microscopy to assess G3BP1 recruitment to dextran-positive compartments. As positive controls, cells loaded with 8 μM Alexa Fluor 568-dextran for 2-hours were treated with either 4 mM LLOMe for 30 min or 0.25 mM sodium arsenite.

### Apilimod-induced endolysosomal expansion

Cells seeded the previous day at 60% confluency were incubated for 24-hours at 37°C and 5% CO₂ in complete DMEM containing 10% FBS, 1% penicillin-streptomycin, 1 µM Janelia Fluor 646-ASO, and 8 µM Alexa Fluor 568-dextran. Janelia Fluor 646-ASO was used for these time-lapse experiments because it was more resistant to photobleaching than the corresponding Alexa Fluor conjugate. Cells were then treated with 5 μM apilimod for 2 h, washed three times, and imaged with a Zeiss Lattice Light Sheet 7 microscope or MOSAIC system. Volumetric images were acquired at 0.25-μm z spacing at 2.3-s intervals.

### SYTOX Green release assay

Endolysosomal release of small-molecule cargo was assessed using SYTOX Green, a cell-impermeant DNA-binding dye, as previously described (37). SVG-A, U2OS, or SUM159 cells were incubated with 5 μM SYTOX Green for the indicated times at 37 °C and 5% CO₂ in the corresponding complete culture medium supplemented with 10% FBS and 1% penicillin– streptomycin. Cells were then washed three times with complete medium and incubated with 0.5 μM Hoechst 33342 for 30 min to label nuclei.

To assess SYTOX Green release from terminal endolysosomal compartments, cells were loaded with 5 μM SYTOX Green for 2 h and chased for 22 h in complete medium. Cells were then treated for 1 h with either 5 μM tantalosin or 0.5 mM LLOMe; 0.5 μM Hoechst 33342 was added during the final 30 min. Cells were imaged by spinning-disk confocal microscopy in complete medium supplemented with 10% FBS and 1% penicillin–streptomycin, with the corresponding treatment maintained throughout imaging.

### ASO uptake

U2OS wild-type, G3BP1/2-knockout and G3BP1/2-knockout cells expressing G3BP1-eGFP-were seeded at 60% confluence in 8-well glass-bottom chambers (Cellvis, C8-1.5H-N) and imaged the next day. Cells were incubated with 1 μM Alexa Fluor 647-ASO and 8 μM Alexa Fluor 568-dextran for 2-hours at 37°C in 5% CO₂. Before imaging, cells were labeled with 100 nM calcein-AM for 5 mins at 37°C in 5% CO₂ to label cytosol for single cell analysis and imaged by live-cell spinning-disk microscopy in FluoroBrite DMEM supplemented with 5% FBS, 1% penicillin-streptomycin and 25 mM HEPES (pH 7.2).

### Nuclear ASO imaging

We quantified the nuclear concentration of fluorescent ASO in U2OS wild-type cells by live-cell 3D LLSM after incubation for 24 h at 37°C with 1 μM Alexa Fluor 647–ASO. We derived ASO concentrations from a fluorescence-intensity calibration curve generated in imaging medium (FluoroBrite DMEM supplemented with 5% FBS and 1% penicillin–streptomycin, pH 7.2) containing increasing concentrations of Alexa Fluor 647 carboxylic acid.

U2OS or SVGA wild-type cells were seeded at 60% confluence on 25-mm coverslips in 6-well plates and imaged the following day. Cells were incubated for 24 h at 37°C and 5% CO₂ in complete medium (DMEM supplemented with 10% FBS and 1% penicillin–streptomycin for U2OS cells) containing 1 μM Alexa Fluor 647–ASO and 8 μM Alexa Fluor 488 as a fluid-phase marker to label endolysosomal compartments. Before imaging, cells were washed three times with complete medium, and nuclei were stained for 30 min at 37°C with Hoechst JF549 in the presence of 25 μM verapamil to inhibit dye efflux and improve intracellular retention of the nuclear stain.

To test the effect of tantalosin on ASO egress, we performed nuclear ASO imaging in SVG-A cells after two shorter pulse–chase protocols. Cells were incubated in MEM supplemented with 10% FBS and 1% penicillin-streptomycin at 37°C with or without 5 μM tantalosin and either 1 μM Alexa Fluor 647–ASO for 2 h or 4 μM Alexa Fluor 647–ASO for 30 min. For experiments with tantalosin treatment on ASO located in the lysosomes, 1 μM Alexa Fluor 647–ASO was pulsed for 2 h, chased for 22 hours and then 5 μM tantalosin was treated for 2 hours before imaging. Cells were then washed three times with MEM supplemented with 10% FBS and 1% penicillin–streptomycin and chased in the same medium for 2 h or 30 min, respectively, to minimize nonspecific fluorescence from ASO bound to the coverslip. Subsequent nuclear staining and imaging were performed as described above.

We selected nuclear regions of interest for LLSM at least 2.5 μm above the coverslip to minimize out-of-focus fluorescence from ASO bound nonspecifically to the glass surface. We also selected regions for which no endolysosomal compartments containing Alexa Fluor 488 and Alexa Fluor 647–ASO lay along the same z-axis, thereby minimizing out-of-focus fluorescence from these bright structures. For each cell, we acquired images sequentially under two conditions. We first collected a rapid image z-stack with a 5-ms exposure per plane per channel to minimize apparent mis-colocalization between Alexa Fluor 488 and Alexa Fluor 647–ASO puncta caused by vesicle movement. We then immediately collected a long-exposure series with a 100-ms exposure per plane per channel to estimate the nuclear ASO concentration.

### Spinning-disk confocal microscopy

Live-cell imaging was carried out on a Zeiss Axio Observer Z1 inverted microscope equipped with environmental control (37°C, 5% CO₂) and a CSU-X1 spinning-disk unit (Yokogawa). The system was operated through the Marianas platform (Slidebook version 2025, 3I Intelligent Imaging Innovations) and included a spherical aberration correction module and a 3i LaserStack with diode lasers at 488, 561 (150 mW) and 640 nm (100 mW). Emission was collected through 525/40, 609/54 and 692/40 filters (Semrock). Z-stacks of 60 optical planes at 0.270 μm spacing were acquired with 20 ms exposure per plane by using a dual-camera sCMOS system (Prime 95B, Teledyne Photometrics) and a custom-made bandpass dichroic (Semrock; transmission 350-560 nm and 641-800 nm, reflection 561-640 nm). Final sampling in the xy plane was 0.145 × 0.145 μm/pixel.

### Lattice light-sheet microscopy

#### LLSM (Zeiss)

Lattice light-sheet imaging was performed with a Zeiss Lattice Lightsheet 7 system using a dithered lattice light sheet (30 μm × 0.7 μm). Cells were maintained in a humidified chamber at 37°C in 5% CO₂. Illumination used 488-nm (10 mW, 2 mW at the pupil), 561-nm (10 mW, 2 mW at the pupil) or 640-nm (5 mW, 1 mW at the pupil) lasers, typically operated at 1-1.5% power. A 13.3× NA 0.4 objective (Zeiss) was used for illumination, and a 44.83× NA 1.0 objective (Zeiss) was used for detection. Z-stacks of 101 planes were acquired in sample-scan mode with 0.5 μm sample spacing and 0.25 μm optical sectioning. Each plane was recorded with 20 ms exposure by using two ORCA-Fusion sCMOS cameras (Hamamatsu), giving xy sampling of 0.145 × 0.145 μm/pixel. Time-lapse imaging was conducted at 2-min intervals for sodium arsenite-induced G3BP1-eGFP recruitment or 20-min intervals for endolysosomal trafficking, over periods of 30 min to 3 h. Raw data were deskewed in ZEISS ZEN 3.11 before analysis.

#### MOSAIC (Multimodal Optical Scope with Adaptive imaging Correction)

An in-house-built lattice light-sheet microscope modified with adaptive optics (MOSAIC) (48) was used for dextran-based endolysosomal pH imaging and nuclear ASO volumetric live-cell imaging by acquiring time points every 2.6 s for 5 min. Sequential images were acquired in sample-scan mode at 0.25 μm spacing along the z axis, and each time point consisted of a z-stack of 81 planes. Samples were illuminated with a dithered multi-Bessel lattice light sheet generated with annular-mask inner and outer numerical apertures of 0.50 and 0.55, respectively. Lasers from MPB Communications Inc. emitting at 488 or 642 nm were used for illumination. A 0.65 NA objective (Special Optics) and a 1.0 NA objective (Zeiss) were used for illumination and detection, respectively. Images were recorded with sCMOS cameras (Hamamatsu, ORCA Flash 4.0 v3) at 0.104 × 0.104 μm/pixel in xy for visualization. Typical exposure times were 10 ms for 488 nm (pH sensitive pHrodo Green dextran) and 10 ms for 642 nm (pH-insensitive Alexa Fluor 647-dextran). Nuclear ASO imaging was performed by acquiring sequential z-stacks of 101 planes, at 5 ms exposure per plane per channel (488 nm, 561 nm, 642 nm) (fast imaging) or 100 ms per plane per channel (long-exposure) in sample-scan mode at 0.25 μm spacing along the z axis.

### Image analysis

#### Colocalization

Colocalization analysis for Figure 2 was performed on 3D binary masks of endolysosomal compartments. Masks were generated by identifying pixels above background with the 3D CME Analysis software package, with the z axis temporarily treated as time (49, 50). After reconversion of the time axis to the z axis, binary images were rescaled to values from 0 to 255 by using the segmentation function in the Fiji 3D Suite(51). Overlap between binary masks from channels of interest was calculated in Fiji by image multiplication. For each segmented object, colocalization was defined as the ratio of integrated binary values within the logical overlap to the integrated binary values of the full 3D object. Objects with at least 50% volumetric overlap were scored as colocalized.

#### Endolysosomal pH

The ratio of fluorescence from pH-sensitive pHrodo Green-dextran to pH-insensitive dextran provided a relative estimate of endolysosomal pH (52). Three-dimensional binary masks of dextran-containing compartments were generated as described above for colocalization analysis. Integrated fluorescence intensity for each object in the 3D masks was quantified with the Fiji “Quantify 3D” function after subtraction of global image background. Only objects larger than 50 voxels were included. The normalized ratio was calculated for each individual object as the ratio of integrated pH-sensitive to pH-insensitive dextran fluorescence, normalized to the median ratio obtained for the corresponding condition after pH 7.4 calibration with clamping solution.

For quantification in Figure S7, 3D binary masks of dextran-containing compartments were generated as described above. Integrated fluorescence intensity for each object was measured with the Fiji “Quantify 3D” function after subtraction of global image background. Median LysoTracker^TM^ Red or dextran intensity within dextran-positive compartments was then calculated for each field of view.

#### ASO uptake

We quantified ASO uptake at single-cell resolution by assigning endolysosomal ASO signal to individual cells. Immediately before live-cell imaging, cells were briefly incubated with calcein-AM to label the cytosol. We used the calcein-AM signal only to obtain a coarse cell outline and thereby separate the endolysosomal content of neighboring cells within the imaging field. Single-cell ROIs were manually drawn on maximum-intensity projections and extended through the full z-stack to generate 3D single-cell ROIs. Within each 3D ROI, we segmented endolysosomal compartments from the dextran channel. We then calculated total ASO uptake per cell by integrating ASO fluorescence over all dextran-positive endolysosomal objects contained within the corresponding single-cell ROI.

#### Nuclear ASO concentration

We generated fluorescence calibration curves relating Alexa Fluor 647 intensity to ASO concentration in the medium imaged by LLSM (MOSAIC) at four exposure times: 20, 100, 200, and 1000 ms. For each concentration and exposure time, we averaged the mean fluorescence intensity from five ROIs. We then fit each calibration dataset by simple linear regression to obtain the relationship between fluorescence intensity and ASO concentration.

We generated binary nuclear masks from Hoechst Janelia Fluor 549 stained cells with the Labkit segmentation tool in Fiji (53, 54). We estimated nuclear volume by counting the voxels in each binary mask and multiplying by the voxel volume, 0.1 × 0.1 × 0.25 µm³. To minimize fluorescence contributed by out-of-focus ASO adsorbed to the coverslip, we measured fluorescence in cell-free regions and estimated the corresponding background contribution across nuclear z-planes. This analysis showed that background contributed less than 10% of the measured signal at distances greater than 2.5 µm above the coverslip, corresponding to 10 z-planes.

We therefore selected five ROIs within each segmented nucleus at least 2.5 µm above the coverslip. We further inspected these ROIs in Mirante4D (55) (https://kirchhausenlab.github.io/llsm_viewer/) to verify that they were spatially separated from bright endolysosomal structures. We determined nuclear ASO concentration by applying the appropriate calibration curve to the mean Alexa Fluor 647 intensity measured in the selected nuclear ROIs. We then calculated the total number of nuclear ASO molecules in each cell from the measured nuclear concentration and the Labkit-derived nuclear volume.

To estimate the fraction of cellular ASO present in the nucleus, we quantified ASO in endolysosomal compartments. We generated endolysosomal masks from Alexa Fluor 488-positive compartments using the 3D CME package (49, 50). We converted the mean ASO fluorescence intensity within these masks to concentration using the same fluorescence calibration curve. We calculated the nuclear fraction as the number of ASO molecules in the nucleus divided by the combined number of ASO molecules in the nucleus and endolysosomal compartments.

#### Nuclear SYTOX Green content

We developed a custom machine-learning pipeline to segment nuclei and quantify nuclear SYTOX Green fluorescence. Multichannel image stacks were separated into Hoechst and SYTOX Green channels. Nuclei were segmented independently in each optical plane of the Hoechst channel to generate 2D binary masks. We refined the masks by removing small spurious objects and incomplete nuclei intersecting the image boundaries, filling internal holes, and interpolating missing masks across planes with discontinuous segmentations. Segmented objects were linked across adjacent planes according to their spatial overlap and merged into coherent 3D nuclear objects, each assigned a unique numerical identifier.

We applied the resulting 3D nuclear masks to the SYTOX Green channel to isolate fluorescence within each nuclear volume. We calculated the mean and integrated SYTOX Green fluorescence intensity for each nucleus. Background fluorescence was measured in a cell-free region of interest and subtracted from the nuclear measurements. For integrated-intensity measurements, the mean background intensity was multiplied by the number of pixels within the corresponding nuclear mask before subtraction. Measurements from multiple image stacks were pooled for comparisons among experimental conditions.

Nuclear segmentation used a deep-learning model based on the Segment Anything Model architecture with a ViT-H backbone (56), implemented through micro-sam (57). Because the pretrained micro-sam model did not segment nuclei with sufficient accuracy, we fine-tuned it using a task-specific annotated dataset. We first used the base micro-sam model to generate segmentation predictions for 40 representative nuclear image stacks. We then manually curated and corrected these predictions using the brush-annotation tools in napari. The resulting training dataset comprised verified ground-truth segmentation masks from 40 image stacks, each containing 26 optical planes. We fine-tuned the ViT-H backbone using these curated data for 50 epochs. Model performance was evaluated using an independent holdout set of 10 nuclear image stacks and confirmed satisfactory segmentation accuracy.

#### ASO in Apilimod-induced expanded endolysosomes

We used live-cell 3D time-lapse datasets to track individual Janelia646-ASO spots within endolysosomal compartments expanded by incubation of the cells with apilimod. For each expanded Alexa Fluor 568-Dextran-positive endolysosome, we defined a 3D spatial constraint centered on the compartment, typically 25 × 25 pixels in xy, corresponding to approximately 2.5 × 2.5 µm, and 9 planes in z corresponding to 0.9 µm. We detected ASO spots in 3D and determined their centroid positions at each time point using 3D CME Analysis software. We assembled ASO trajectories by linking detections across frames based on spatial confinement and signal continuity.

We calculated the mean-squared displacement (MSD) from the time series of 3D centroid positions for each ASO particle associated with an endolysosomal compartment. For each trajectory, we computed the time-averaged squared displacement as a function of lag time. We classified particle motion from the lag-time dependence of the MSD. Freely diffusing particles showed an initial rise in MSD followed by a plateau, consistent with confinement within a bounded endolysosomal volume. Particles associated with endolysosomal structural features, such as the limiting membrane, lacked a clear plateau and showed broader MSD variation after the initial rise. This behavior is consistent with motion constrained along a curved surface and influenced by endolysosomal dynamics.

#### LLOMe-mediated release

We generated 3D binary masks of ASO-containing compartments as described for the colocalization analysis. After subtracting the background signal corresponding to regions outside the cell, we quantified the integrated fluorescence intensity of each masked object with the Fiji Quantify 3D function. We included only objects larger than 50 voxels.

We used the temporal progression of endosomal cargoes to assess the effect of LLOMe on dextran and ASO release. Sequential labeling with two spectrally distinct fluorescent ASOs enabled classification of compartments along the endosomal pathway. Terminal lysosomes were defined as compartments labeled exclusively with Alexa Fluor 488-ASO following an overnight pulse-chase. Endolysosomes were defined as compartments containing both Alexa Fluor 488-ASO from the long chase period, 18 h, and Alexa Fluor 647-ASO from a continuous 2.5-hours pulse. Late endosomes were defined as compartments positive for Alexa Fluor 647-ASO but negative for Alexa Fluor 488-ASO. Release was established by measuring the relative amounts of dextran (Alexa Fluor 568-Dextran) and ASO fluorescence within the 3D masks containing both ASO signals (e.g. lysosomes) in images acquired before and after LLOMe treatment. We also quantified in the same images the Alexa Fluor 647-ASO intensity in late endosomal and lysosomal compartments before and after LLOMe addition.

#### Analysis of Fluorescence Loss upon Ionomycin-Mediated Exocytosis

We acquired three-dimensional spinning-disk confocal time series of cells containing Alexa Fluor 568–labeled 10-kDa dextran and Alexa Fluor 647–labeled ASO. Multiple fields of view, each containing several cells, were imaged sequentially, with each cell sampled at 1-min intervals for up to 16 min. Fluorescent spots were detected and segmented in three dimensions using 3D-CME(58). Objects were linked between consecutive frames by one-to-one centroid assignments within five voxels, without gaps or branching. Fluorescence intensities were corrected by subtracting the median intensity of a surrounding three-dimensional background shell; dextran signals also received an empirical mean-decay correction.

Candidate dextran-loss events required a step model with a Bayesian information criterion (BIC) at least six units lower than that of a linear trend, decreases of at least 30% between adjacent frames and between pre-and post-drop medians, and at least 80% of post-drop observations at or below 75% of the pre-drop median. Trajectories spanning at least six consecutive frames and their paired dextran and ASO fluorescence measurements were displayed in an interactive HTML viewer. The first acquisition frame was excluded from plots and decay fits. We validated each candidate event by visual inspection. The viewer and analysis scripts are available in our GitHub repository.

### Volume Electron Microscopy

#### High-Pressure Freezing, Freeze-Substitution and Embedding

High-pressure freezing, freeze-substitution and embedding were carried out as described previously (52, 59). Cells were plated on 6 x 0.1-mm sapphire disks in complete DMEM containing 10% FBS and 1% penicillin-streptomycin. For freezing, each disk was sandwiched between a gold-plated spacer with a 150-μm recess (Technotrade, 1649-100) and a flat-bottom gold spacer (Technotrade, 1653-100), and frozen with a Wohlwend HPF Compact 03 high-pressure freezer (Technotrade).

Under liquid nitrogen, frozen samples were transferred to cryotubes containing substitution medium composed of 2% OsO₄, 0.1% uranyl acetate and 3% water in acetone. Freeze-substitution used an EM AFS2 system (Leica Microsystems) programmed to warm from −140°C to −90°C for 2 h, hold at −90°C for 24 h, warm to 0°C for 12-hours and then to 22°C for 1 h. Samples were rinsed sequentially three times with anhydrous acetone, propylene oxide and 50% Embed 812 resin in propylene oxide, transferred to embedding molds (EMS, 70900) containing 100% resin and polymerized at 65°C for 48 h. Sapphire disks were then removed from the resin blocks by alternating immersion in liquid nitrogen and boiling water.

#### Crossbeam FIB-SEM Imaging

As described previously (59), we detached polymerized resin blocks from the molds and mounted them on aluminum pin stubs (Ted Pella) with conductive silver epoxy (EPO-TEK H20S, Electron Microscopy Sciences), leaving the block face exposed. We coated the exposed surface with 20 nm carbon in a Quorum Q150R ES sputter coater (Quorum Technologies) and transferred the sample to a Zeiss Crossbeam 540 microscope for FIB-SEM imaging. After eucentric alignment, we tilted the stage to 54° and set the working distance to 5 mm for beam calibration. We identified the cell of interest by SEM, milled a coarse trench adjacent to the cell with a 30 kV, 30 nA gallium ion beam, and polished the exposed block face at 30 kV, 7 nA. We acquired serial images by interlacing FIB milling with a 30 kV, 3 nA gallium beam and SEM imaging at 1.5 kV, 400 pA, using 10 nm milling steps to generate isotropic 10 nm voxels (10 x 10 x 10 nm). Images were collected from the InLens secondary-electron and SEBI backscattered-electron detectors, with a pixel dwell time of 20 μs and line averaging of 1. Resulting InLens images were denoised using the FastFIB pipeline (Nair et al, JCB 2025) (60).

### Statistical analysis

Statistical significance between control and experimental groups was assessed with a nonparametric one-way analysis of variance. Asterisks denote P values as follows: P < 0.05 (*), P < 0.01 (**), P < 0.001 (***) and P < 0.0001 (****). Comparisons without indicated P values were not significant.

## RESULTS

### Cell models for imaging ASO trafficking and measuring its functional delivery

We sought an adherent cell line suitable for high-quality live-cell imaging, responsive to ASOs with measurable functional delivery, and amenable to testing whether endosomal escape and productive ASO delivery depend on G3BP1/2, core RNA-binding proteins that nucleate stress granules and bind ASOs (61). We therefore compared SUM159, SVGA, and U2OS cells for their response to a phosphorothioate (PS)-modified ASO (hereafter, ASO) that directs RNaseH1-dependent degradation of *Malat1* RNA. qPCR measurements showed a concentration-dependent decrease in *Malat1* RNA in U2OS and SVGA cells, but little or no response in SUM159 cells under the same conditions (Supplementary Fig. S1). This weak ASO response may reflect inefficient escape during early endocytosis, consistent with their lack of an early response to tantalosin stimulation, as described below. We therefore chose U2OS and SVGA cells for most subsequent experiments as they are well suited for live-cell fluorescence microscopy and because a U2OS cell line is available with simultaneous knockout of G3BP1 and G3BP2 (62). This system allowed direct tests of whether G3BP1/2 links endosomal escape to productive ASO delivery. We used live-cell rather than fixed-cell imaging to avoid fixation-induced permeabilization of endolysosomal compartments and consequent redistribution of luminal contents.

### ASOs enter U2OS cells by endocytosis and accumulate in late endolysosomal compartments

Fluorescently labeled ASOs entered U2OS cells readily from the medium and accumulated in punctate structures that colocalized with internalized dextran, consistent with sequestration in the endolysosomal system (Fig. 1A, B). Co-incubation for 2-hours with pH-insensitive Alexa Fluor 568-labeled 10-kDa dextran gave extensive colocalization (Fig. 1C), indicating uptake through a common endocytic pathway and passage through early and late endosomes before accumulation in late endosomes and lysosomes. Similar accumulation profiles supported similar uptake and trafficking kinetics.

**Figure 1.**
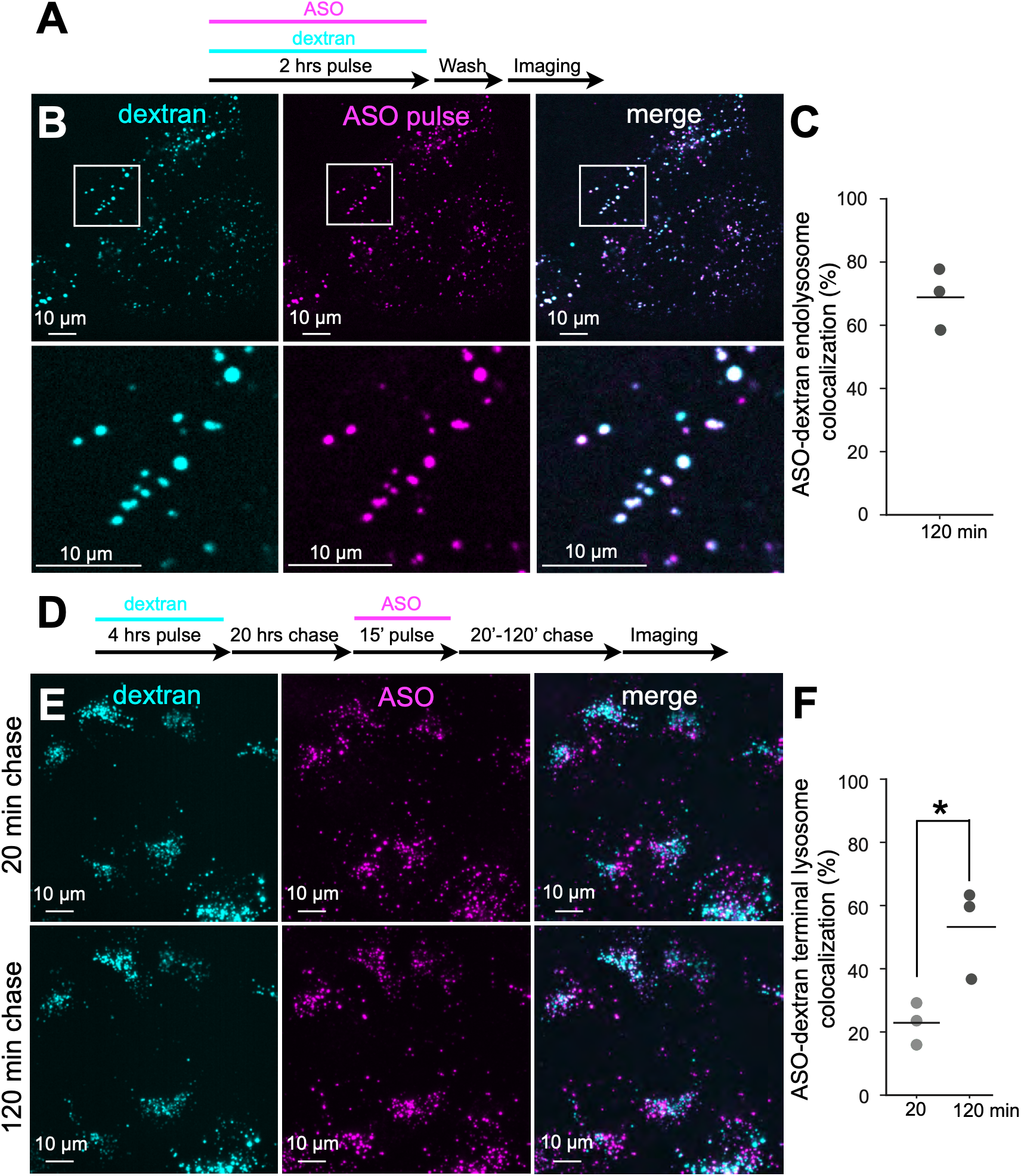
Naked ASOs enter and accumulate in dextran-labeled endolysosomal compartments(A) Experimental protocol. Cells were co-incubated for 2-hours with 8 μM Alexa Fluor 568-dextran and 1 μM Alexa Fluor 647-ASO, washed, and live-cell imaged. **(B)** Representative maximum-projection spinning-disc images of Alexa Fluor 568-dextran (cyan) and Alexa Fluor 647-ASO (magenta) in wild-type U2OS cells after a 2-hours pulse. Scale bar, 10 μm. **(C)** Quantification of the fraction of dextran-labeled compartments that colocalize with ASOs (# endosomes=16377) in **(B)**. The median is shown as a black line; each dot represents an independent experiment, for each experiment the data are from at least 10 fields of view. **(D)** Experimental protocol. 8 μM Alexa Fluor 568-dextran was pulsed for 4-hours and chased for 20-hours to label terminal compartments, followed by a 15 min pulse with 1 μM Alexa Fluor 647-ASO. Cells were imaged after 20 min and 120 min chase. **(E)** Representative maximum-projection lattice light-sheet images of Alexa Fluor 568-dextran (cyan) and Alexa Fluor 647-ASO (magenta) in wild-type U2OS cells after a 20 min chase (upper) or a 120 min chase (lower). Scale bar, 10 μm. **(F)** Quantification of the fraction of dextran-labeled terminal compartments that is colocalizing with Alexa Fluor 647-ASO in **(E)** after 20 min (# endosomes = 7,388) and 120 min chase (# endosomes = 8,323). The median is shown as a black line; each dot represents an independent experiment. Statistical significance p<0.05 (*).

**Figure 2.**
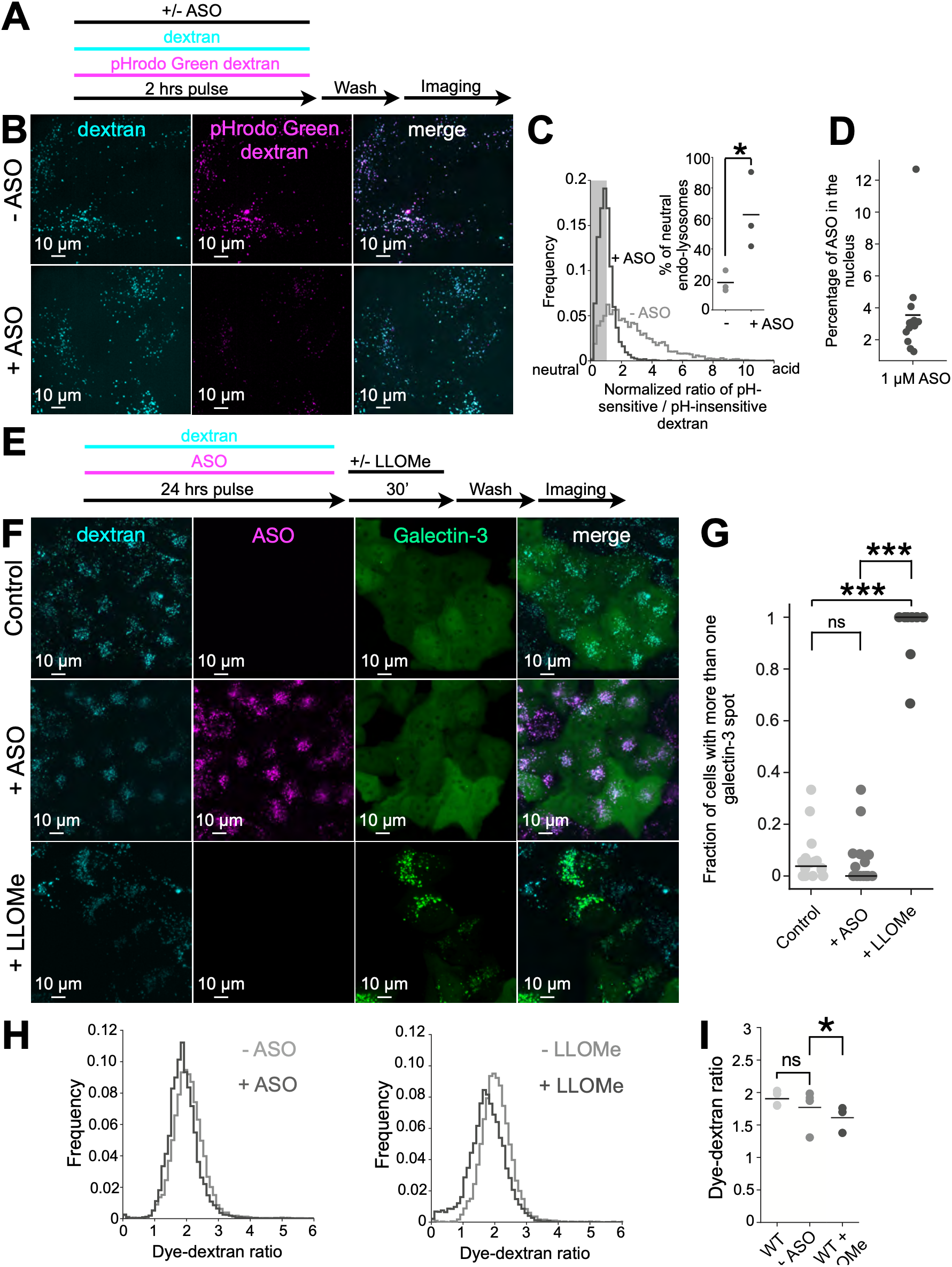
ASOs neutralize a subset of endolysosomal compartments(A) Experimental protocol. Cells were incubated for 2 hours with 8 μM Alexa Fluor 647-dextran (pH-insensitive) and 8 μM pHrodo Green-dextran (pH sensitive) in presence or absence of 1μM non-fluorescent ASOs, washed and imaged. **(B)** Representative spinning-disc microscopy max-projected images of Alexa Fluor 647-dextran pulse (cyan) and pHrodo Green-dextran pulse (magenta) in absence (upper) and presence (lower) of non-fluorescent ASOs. Data are representative from one of three independent experiments. Scale bar: 10 μm. **(C)** Quantification of endolysosomal pH in U2OS wild-type from **(B)**, calculated using the ratio of pH-sensitive pHrodo Green-dextran to pH-insensitive dextran fluorescence normalized to mean + 1 standard deviation of pH 7.4 calibrated fluorescence. Frequency plots show the distribution of pH-sensitive/insensitive ratio from one of three biological replicates, 10 fields of view per replicate, (total # endosomes = 15,476 (-ASO) and 16,093 (+ASO)) for wild-type U2OS cells. The shaded region indicates mean +1 standard deviation of the pH 7.4 clamped ratio of pH sensitive to insensitive fluorescence. The inset shows percentage of neutral endosomes within mean +1 standard deviation of pH 7.4 clamped value of pH-sensitive/insensitive ratio. Endosomes are considered neutral if they are within the shaded region of the distribution. Statistical significance: p>0.05 (ns); p<0.05 (*). **(D)** Quantification of the percentage of Alexa Fluor 647-ASO localized to the nucleus relative to total cellular ASO in wild-type U2OS cells following 24-hours incubation with 1 μM Alexa Fluor 647-ASO. The median is shown as a black line; each dot represents a cell. Data are from two independent experiments. **(E)** Experimental protocol. 8 μM Alexa Fluor 568-Dextran and 1 μM Alexa Fluor 647-ASO were co-pulsed for 24 hours and further incubated in presence or absence of 0.5 mM LLOMe prior to imaging. **(F)** Representative spinning-disc confocal microscopy images (max intensity projections) of Alexa Fluor 568-Dextran (cyan) in U2OS wild-type stably expressing eGFP-galectin-3 (green) as untreated (control), treated with 1 μM Alexa Fluor 647-ASO (magenta) for 24 hours or 0.5 mM LLOMe for 30 minutes. Data are representative from one of two independent experiments. Scale bar 10 µm. **(G)** Quantification of **(F)** as fraction of cells with one or more galectin-3 spots in the cytoplasm. Median fractional value is represented as black line, where each dot represents fraction of cells in field of view (FOV). Data are from two independent experiments. Statistical significance: p>0.05 (ns); p<0.001 (***). **(H)** Frequency distributions of the ratio of Alexa Fluor 568 intensity to Alexa Fluor 647-dextran intensity within endolysosomal compartments (# endosomes = 8,687) following treatment with 1 μM non-fluorescent ASO (# endosomes = 10,098) or 0.5 mM LLOMe (# endosomes = 10,221). Data are representative from one of three independent experiments. **(I)** Quantification of panel **(H)**, shown as the median Alexa Fluor 568 (dye): Alexa Fluor 647-dextran (dextran) intensity ratio in endolysosomal compartments in the presence or absence of 1 μM non-fluorescent ASO or 0.5 mM LLOMe. The median is shown as a black line; each dot represents an independent experiment. Statistical significance: p>0.05 (ns); p<0.05 (*).

To benchmark delivery to terminal lysosomes, we pulse-labeled cells with Alexa Fluor 568-labeled 10-kDa dextran (4-hours pulse, 20-hours chase; Fig. 1D) and added pH-insensitive Alexa Fluor 647-labeled ASO during the final 2-hours. ASO fluorescence shifted progressively to dextran-labeled terminal lysosomes during the chase. Overlap was limited at 20 minutes, consistent with passage through earlier endocytic compartments, but increased substantially by 120 minutes and approached that of dextran in those compartments (Fig. 1E, F; supplementary video S1 and S2).

### ASOs neutralize endolysosomes without detectable cytosolic release

Endosomes and lysosomes maintain an acidic lumen through V-ATPase-driven proton pumping and low permeability of the limiting membrane (63). In U2OS cells, ratiometric imaging with pHrodo Green–labeled 10-kDa dextran, whose fluorescence decreases as pH rises over approximately pH 4–8 (apparent pKa ≈6.8), and a pH-insensitive Alexa Fluor 647–labeled 10-kDa dextran reference showed that endosomes and lysosomes remained predominantly acidic under basal conditions (Fig. 2A–C; Supplementary Fig. S2A). Incubation with 1 uM of our PS modified ASOs shifted the luminal pH distribution toward higher values. We also observed a shift toward higher luminal pH at 300 nM ASO (Supplementary Fig. S2C). Many ASO-accessible endosomes and lysosomes became partially neutralized, and a substantial fraction became fully neutralized (Fig. 2B, C). We quantified these distributions by live 3D volumetric imaging, sampling thousands of endosomal compartments per condition (Fig. 2A-C and Supplementary Fig. S2A). Although imaging of a single cell volume does not report the lifetime of an individual neutralization event, the abundance of neutral compartments implies persistence at least over the timescale of volumetric acquisition. Short-interval volumetric time series (a cell imaged every 2.6 seconds) supported that inference: neutralized compartments persisted across successive stacks (Supplementary Fig. S2B).

Despite this marked neutralization, we detected only trace diffuse ASO fluorescence in the nucleus above background. We incubated cells with 1 μM fluorescent ASO for 24 hours, a condition that in independent experiments reduced *Malat1* RNA in U2OS cells by more than 90%, as measured by qPCR (Supplementary Fig. S1). We then used lattice light-sheet microscopy (LLSM), calibrated for single-molecule detection, to estimate the total number of ASO molecules in endosomes and lysosomes, and LLSM calibrated for non-point-source fluorescence to measure total nuclear ASO signal (Fig. 2D, Supplementary Fig. S3A-G). These measurements indicated that, after 24-hours of continuous incubation, no more than ∼4% of the ASO retained in endosomes and lysosomes had been released and accumulated in the nucleus. Thus, under these conditions, ASOs accumulated in endolysosomal compartments with only minimal release.

Our *in vitro* experiments used 1 μM ASO, near the lower end of the micromolar concentrations used in published gymnotic uptake studies (6, 64). This exposure may exceed that encountered by some cells *in vivo*, and substantially lower concentrations might not produce detectable neutralization. Our results nevertheless show that, when neutralization occurs, any associated membrane perforations are insufficient to release a substantial fraction of the accumulated ASO.

### ASOs remain sequestered despite LLOMe-induced limiting-membrane damage

While ASOs neutralized endolysosomal compartments, we had no evidence in support of overt membrane rupture, as judged by the absence of cytosolic eGFP-galectin-3 recruitment, a reporter of substantial endolysosomal damage (36, 65) (Fig. 2E-G). ASO treatment did not increase the number of galectin-3 puncta above baseline (Fig. 2F, + ASO and Fig. 2G), despite causing strong neutralization (Fig. 2C). In contrast, the positive control, a 30-min exposure to 0.5 mM L-leucyl-L-leucine methyl ester (LLOMe), produced abundant galectin-3 puncta (Fig. 2F, + LLOMe and Fig. 2G), confirming that the assay sensitively detected LLOMe-induced injury. We infer that ASO-associated neutralization reflects either membrane lesions below the threshold required for galectin-3 recruitment or impaired acidification without large discontinuities in the limiting membrane. The latter interpretation is more likely. In U2OS cells incubated with a 1:1 mixture of Alexa Fluor 568 and Alexa Fluor 647 10 kDa dextran, ASO did not increase escape of the small fluorescent dye (Fig. 2H, I). By contrast, LLOMe, used as a positive control, produced an endolysosomal population with reduced dye content, consistent with membrane damage (Fig. 2H, I).

We next asked whether LLOMe-induced permeabilization releases ASOs from endolysosomes, as it releases dextran, used here as a soluble luminal tracer. We used 10-kDa dextran as a size-matched control (Stokes radius, ∼2.3 nm) relative to a ∼20-nucleotide single-stranded ASO (∼1-2 nm) (66). After a 3-hour pulse and 18-hour chase with Alexa Fluor 488-ASO to label terminal lysosomes, we co-internalized Alexa Fluor 647-ASO with Alexa Fluor 568-dextran for 2.5-hours to label late endosomes and endolysosomes and quantified fluorescence in these compartments before and after a 30-minutes exposure to 0.5 mM LLOMe (Fig. 3A). LLOMe rapidly depleted dextran from punctate compartments and generated diffuse weak cytosolic fluorescence, consistent with leakage from these damaged compartments (Fig. 3B-D; Supplementary Fig. S4-S5, Supplementary video S3). The unconjugated Alexa Fluor 647 carboxylic acid used to label the ASO also accumulated in the same endosomal compartments and escaped after LLOMe treatment. By contrast, fluorescence from the labeled ASO remained organelle-associated without a detectable decrease, and no diffuse signal appeared in the cytosol or nucleus above our detection threshold (Fig. 3B–D; Supplementary Fig. S5A, B). If ASO degradation released free dye, that dye would therefore be expected to escape under these conditions. The persistence of fluorescence argues against free dye as the source of the retained signal and is consistent with retention of intact ASOs in late endosomes, endolysosomes, and lysosomes. Thus, lesions sufficient to release soluble 10-kDa dextran from late endosomes, endolysosomes, and terminal lysosomes did not measurably release co-internalized ASO under the same conditions. Given the sensitivity limits of our fluorescence assay, these data do not exclude rare release events or escape from compartments not well sampled in this analysis, such as early or recycling endosomes (see next section). LLOMe treatment did not significantly enhance the activity of unlabeled ASO, as measured by qPCR (Supplementary Fig. S9C). This result is consistent with minimal additional release of functional ASO and argues against fluorophore conjugation as the cause of retention.

**Figure 3.**
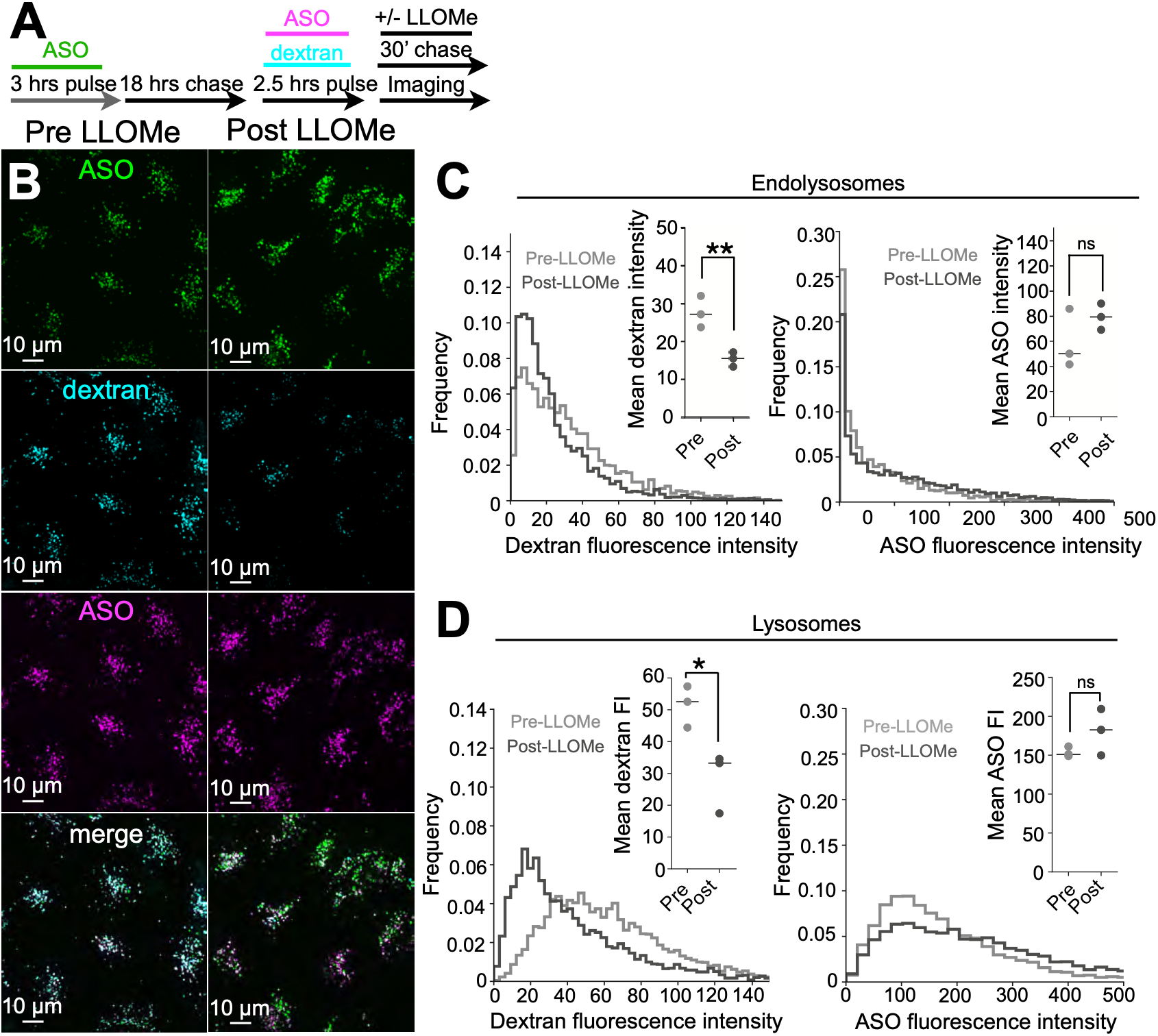
ASOs remain luminally sequestered after extensive endolysosomal damage.(A) Experimental protocol. Wild-type U2OS cells were preloaded with 1 μM Alexa Fluor 488-ASO for 3-hours and chased for 18-hours to terminal lysosomes (green). Cells were then pulsed for 2.5-hours with 8 μM Alexa Fluor 568-dextran (cyan) and 1 μM Alexa Fluor 647-ASO (magenta) to late endosomes, endolysosomes and lysosomes, with or without 0.5 mM LLOMe for 30 min before imaging. **(B)** Representative maximum-projection spinning-disc images of wild-type U2OS cells preloaded with Alexa Fluor 488-ASO and chased to terminal lysosomes (green), then pulsed for 2.5-hours with Alexa Fluor 568-dextran (cyan) and Alexa Fluor 647-ASO (magenta) to label late endosomes, endolysosomes and lysosomes, without treatment (left) or after treatment with 0.5 mM LLOMe for 30 min (right). Scale bar, 10 μm. **(C)** Quantification of **(B)**, shown as frequency plots of Alexa Fluor 568-dextran fluorescence (left) and Alexa Fluor 647-ASO fluorescence (right) in endolysosomal compartments negative for Alexa Fluor 488-ASO after an 18-hours chase and positive for Alexa Fluor 647-ASO after 2.5-hours pulse in wild-type cells, before and after treatment with 0.5 mM LLOMe for 30 min (# endosomes pre-LLOMe = 10,197; post-LLOMe = 15,539). Insets show median fluorescence intensities from the distributions, plotted as dot plots. Each dot represents an independent biological replicate. Data are from three biological replicates. Statistical significance: p>0.05 (ns); p<0.05 (**). **(D)** Quantification of **(B)**, shown as frequency plots of Alexa Fluor 568-dextran fluorescence (left) and Alexa Fluor 647-ASO fluorescence (right) in lysosomal compartments positive for both Alexa Fluor 647-ASO after 2.5-hours pulse and for Alexa Fluor 488-ASO after an 18-hours chase in wild-type cells before and after treatment with 0.5 mM LLOMe for 30 min (# endosomes pre-LLOMe = 16,092; post-LLOMe = 14,404). Insets show median fluorescence intensities from the distributions, plotted as dot plots. Each dot represents an independent biological replicate. Data are from three biological replicates. Statistical significance: p>0.05 (ns); p<0.05 (*).

A pre-print study published while our work was in progress examined the response to LLOMe during the first few minutes of treatment, whereas we followed this response for up to 30 min (67). That study detected luminal neutralization and galectin-8 recruitment (another reporter of membrane damage) but no loss of dextran fluorescence, and attributed ASO retention to condensation. Without evidence that the membrane had become permeable to a soluble tracer of comparable hydrodynamic size, however, failure to observe ASO release cannot distinguish retention by condensation from insufficient membrane permeability. In our experiments, LLOMe released dextran while ASOs remained associated with the damaged compartments marked by galectin-3 (Supplementary Fig. S5A, B).

To determine whether ASO retention was specific to LLOMe treatment, we used ionomycin to trigger Ca²⁺-dependent lysosome exocytosis, which opens the lysosomal lumen directly to the extracellular medium (68, 69). Cells were loaded with Alexa Fluor 568–labeled 10-kDa dextran and Alexa Fluor 647–labeled ASO, then chased overnight to allow both probes to accumulate in terminal lysosomes. Before inducing exocytosis, we briefly incubated the cells with Versene, an EDTA-containing dissociation solution, and transferred them to fresh coverslips that had not been exposed to ASO. After an additional day in culture, we imaged individual lysosomes at the ventral cell surface by three-dimensional live-cell microscopy. Transfer to fresh coverslips reduced background fluorescence from ASOs adsorbed to the glass during loading (Supplementary Fig. S6A).

Before inducing exocytosis, we observed discrete ASO foci superimposed on a more uniform luminal ASO signal. Dextran overlapped with the uniform component but showed no corresponding foci. We then triggered exocytosis and followed both cargoes, identifying 56 lysosome-fusion events in 67 cells from two independent experiments. In each event, dextran fluorescence disappeared rapidly after fusion, consistent with release into the extracellular medium. ASO fluorescence frequently showed significantly less decrease; when it declined, residual signal persisted at the fusion site throughout the acquisition. In a representative event, single-particle tracking showed loss of the dextran track at 7th min after the start of imaging, whereas the corresponding ASO track persisted until the end of the time series (Supplementary Fig. S6B, C). Thus, direct exposure of the lysosomal lumen to the extracellular medium released dextran while leaving a substantial fraction of ASO associated with the residual structure at the fusion site. ASO retention therefore extended beyond LLOMe-induced membrane damage.

By chance, we captured a single spontaneous dextran-release event in a 3D live-cell time series of apilimod-treated cells. Apilimod inhibits PIKfyve and enlarges late endosomes and lysosomes, allowing their luminal contents to be resolved (70). One endolysosome lost its dextran signal within a single frame and subsequently contracted, retaining both an ASO foci and the surrounding distributed signal without detectable loss of total ASO fluorescence (Supplementary Video S4). This event provided additional evidence for ASO retention despite release of soluble luminal cargo.

We further used these enlarged compartments to examine systematically the distribution of internalized ASOs relative to their luminal contents and limiting membranes. Alexa Fluor 568– labeled 10-kDa dextran filled the enlarged lumen uniformly, as expected for an unbound soluble tracer (Fig. 4A,B, dextran; Supplementary Fig. S7A,B; Supplementary Video S5). By contrast, Alexa Fluor 647–ASO and Janelia Fluor 646–ASO showed strikingly heterogeneous distributions (Fig. 4B, ASO; Supplementary Fig. S7A,B; Supplementary Fig. S8A-C; Supplementary Videos S5). Both ASO probes frequently concentrated in discrete intraluminal foci, consistent with earlier reports of punctate intraluminal ASO signal (71), and along the limiting membrane. We tracked these foci in three dimensions by live-cell LLSM imaging at 2.3-s intervals (Fig. 4C; Supplementary Fig. S7C). Their trajectories showed constrained movement rather than free diffusion. We used the same analysis recently applied to virions in apilimod-expanded endolysosomes (72) which similarly permitted direct visualization of lumen-associated particles.

**Figure 4.**
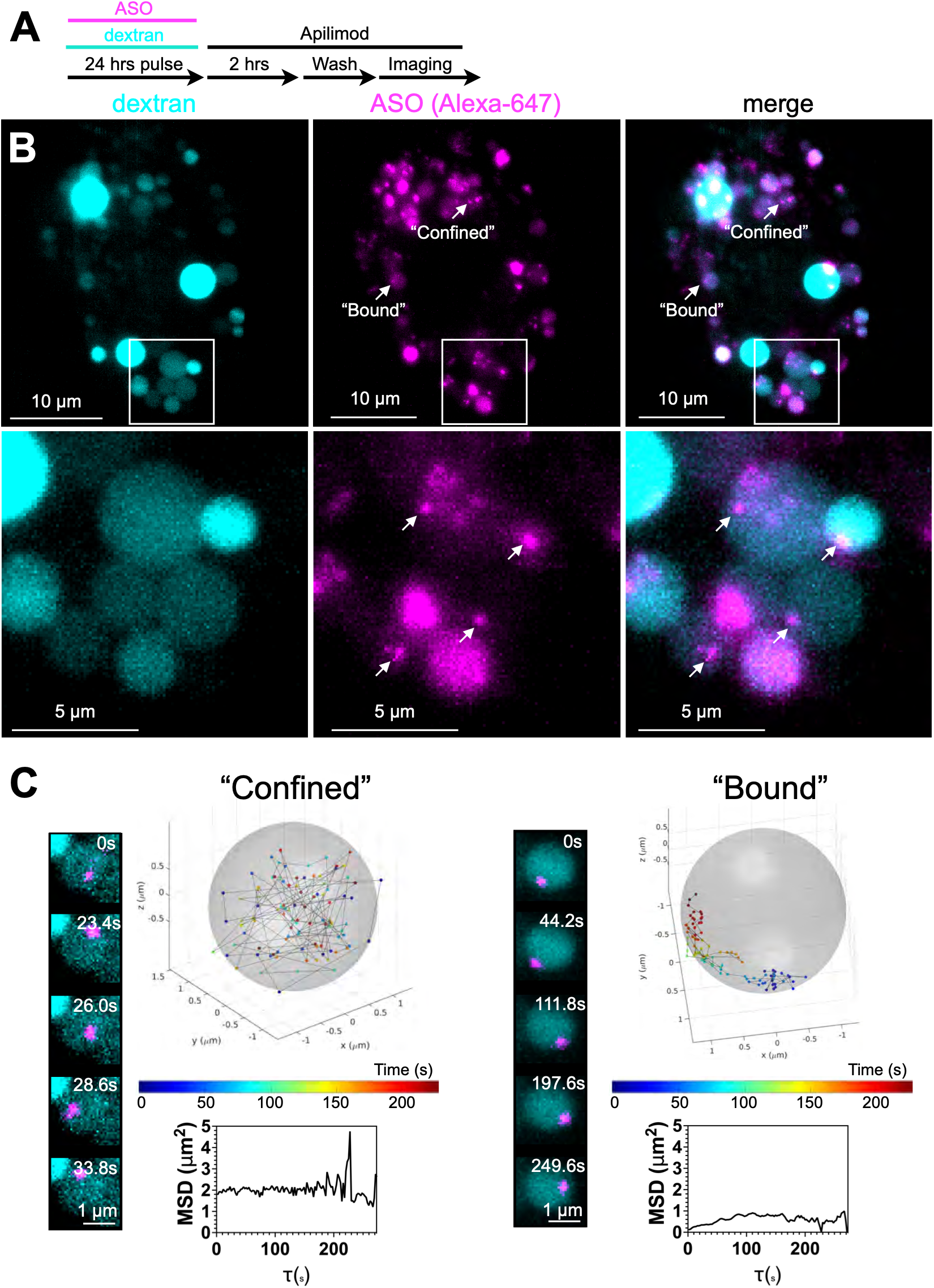
ASOs are heterogeneously distributed within the endolysosomal lumen.(A) Experimental protocol. Wild-type U2OS cells were preloaded for 24-hours with 8 μM Alexa Fluor 568-dextran and 1 μM Alexa Fluor 647-ASO, chased for 2-hours in the presence of 5 μM apilimod, washed, and imaged. **(B)** Representative lattice light-sheet (Zeiss) images of wild-type U2OS cells labeled as in **(A)**. A single middle plane is shown. Alexa Fluor 568-dextran is shown in cyan and Alexa Fluor 647-ASO in magenta. Arrows indicate ASO puncta undergoing confined and bound motion. Scale bar, 10 μm. Lower panel: Arrows indicate discrete punctate of intraluminal ASO in endolysosomes, whereas the rest of the intraluminal ASO and dextran fills the same compartment uniformly. Scale bar, 5 μm. **(C)** Representative three-dimensional trajectories of individual ASO particles exhibiting confined or bound motion within enlarged endolysosomal compartments. Endolysosomes are rendered as semitransparent spheres, and ASO positions are color-coded by time from blue to red. Insets show time-resolved maximum-intensity projections of representative ASO-containing endolysosomes, with ASO in magenta and dextran in cyan. Scale bar, 1 μm. Mean squared displacement plots are shown for each trajectory. Data are representative from one of three independent experiments.

We used Janelia Fluor 646–ASO for MSD analysis because its greater photostability than Alexa Fluor 647 enabled longer trajectories and more reliable estimates. Alexa Fluor 647–ASO preserved ASO activity in U2OS cells at 1 μM (Supplementary Fig. S9A), whereas Janelia Fluor 646 conjugation reduced activity, as measured by quantitative PCR (Supplementary Fig. S9B). Nevertheless, the two conjugates showed indistinguishable heterogeneous distributions and constrained movements within apilimod-expanded endolysosomes (Fig. 4C; Supplementary Fig. S7C).

Taken together, these observations indicate that ASOs are retained extensively within endocytic compartments, particularly in late endosomes, endolysosomes, and terminal lysosomes. The retention may reflect association with luminal components, the limiting membrane, or both; its molecular basis remains unknown. Such association would be expected to restrict ASO release even when membrane permeability increases.

### Tantalosin promotes early endosomal escape of luminal cargo and ASOs

Our preceding results pointed to an infrequent escape event during early endocytic trafficking, before ASOs accumulate in late endosomes and lysosomes. Early and recycling endosomes undergo repeated membrane tubulation, constriction, and fission. The ESCRT-III protein IST1 contributes to recycling-tubule scission and cargo transport from early endosomes to the recycling compartment (30, 32). Independent studies have established that ESCRT assemblies also promote repair of damaged endolysosomal membranes (34–36). These observations led us to ask whether IST1-containing assemblies help close transient membrane discontinuities that might arise during recycling-tubule remodeling.

Tantalosin provided a candidate tool for acutely perturbing this process. The compound was reported to inhibit recycling of internalized transferrin and disrupt IST1–CHMP1B copolymer formation in vitro ((73), subsequently retracted). In their retraction notice, the authors reaffirmed the effect on copolymer formation but withdrew the conclusion that tantalosin reduced IST1– CHMP1B binding affinity (74). We therefore used tantalosin to test whether acute perturbation of IST1-associated remodeling would increase luminal escape, as expected if it delayed closure or repair of transient openings. We first monitored escape of internalized SYTOX Green, whose fluorescence increases strongly upon binding DNA, allowing nuclear fluorescence to serve as a sensitive readout of cytosolic access. We then examined nuclear accumulation and functional activity of internalized ASOs.

We synthesized tantalosin as described previously and verified its purity by NMR (Supplementary Fig. S8D) (75). We first examined its effect on transferrin recycling (Supplementary Fig. S10). In untreated U2OS, SVG-A, and SUM159 cells, fluorescent transferrin entered peripheral endosomes. During the chase, intracellular fluorescence declined rapidly in U2OS and SVG-A cells, consistent with recycling to the cell surface (76). Tantalosin delayed clearance, with a pronounced effect in U2OS and SVG-A cells and a smaller effect in SUM159 cells. Transferrin accumulated in perinuclear compartments and persisted during the chase. Their distribution and prolonged cargo retention were consistent with late endosomes or slowly recycling compartments.

Previous work established a role for IST1 in early endosomal sorting: siRNA-mediated depletion impaired transferrin receptor delivery to the endocytic recycling compartment while increasing rapid recycling through peripheral endosomes (32). In our gene-edited cells expressing IST1– Halo, tantalosin increased IST1 accumulation on transferrin-positive early and/or recycling endosomes (Supplementary Fig. S11). Together with delayed transferrin recycling, this redistribution supports an effect on IST1-associated endosomal remodeling, although it does not identify the compound’s direct molecular target. Although siRNA-mediated depletion of VPS23 or VPS28, components of ESCRT-I, has been reported to enhance the activity of transfected ASOs (77), depletion of TSG101 or VPS28 also enhanced the activity of naked anti-miR-21 (78). Although the latter cargo was anti-miRNA rather than an ASO, it provides the more relevant precedent because it entered without transfection. Its sensitivity to depletion of an ESCRT-I component is also consistent with our tantalosin experiments, for which the presumed target, IST1, is an ESCRT-III protein. We nevertheless did not pursue IST1 depletion or genetic ablation, because sustained IST1 loss has pleiotropic effects, including inhibition of abscission (79).

Early and recycling endosomes undergo extensive tubulation and fission during cargo sorting (29, 80). Given the established role of ESCRT machinery in membrane repair, we postulated that IST1-containing ESCRT-III assemblies might help close transient membrane discontinuities arising at sites of fission. Interference with this activity could prolong such openings and increase luminal cargo escape. We therefore tested whether tantalosin enhanced escape during early endocytosis. Dextran lacked sufficient sensitivity: the amount internalized per early endosome was below our detection limit, and any released dextran would become too dilute in the cytosol to detect. We instead used SYTOX Green, which accumulates in the nucleus after gaining cytosolic access and becomes strongly fluorescent upon binding DNA.

Co-incubation with tantalosin increased nuclear SYTOX Green fluorescence in U2OS and SVG-A cells within 2 h (Fig. 5A,C). The increase was detectable after 30 min (Fig. 5B), and time-lapse imaging beginning 10 min after dye addition showed elevated nuclear fluorescence throughout the subsequent 160 min (Fig. 5D). To distinguish escape during early trafficking from release after lysosomal accumulation, we chased SYTOX Green into terminal lysosomes before adding tantalosin. Nuclear fluorescence increased only modestly under these conditions, whereas LLOMe produced a strong response (Supplementary Fig. S12A–C). Tantalosin thus acted preferentially before cargo reached terminal lysosomes.

**Figure 5.**
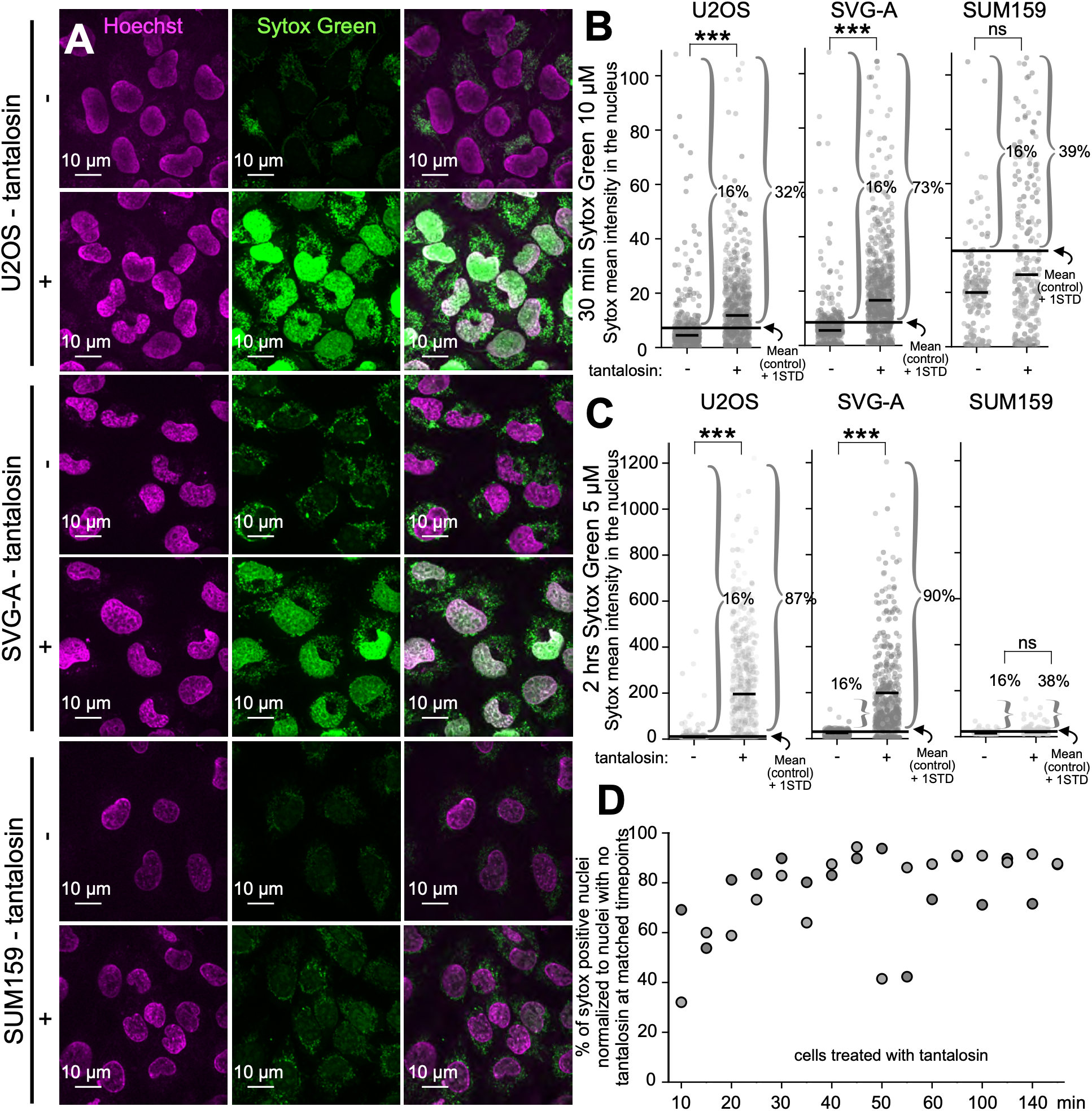
Tantalosin promotes early endosomal escape of SYTOX Green (A) Representative maximum-intensity projections from spinning-disk confocal image stacks of wild-type U2OS (top), SVG-A (middle), and SUM159 (bottom) cells incubated with 5 μM SYTOX Green in the absence or presence of 5 μM tantalosin. SYTOX Green is shown in green and Hoechst-labeled nuclei in magenta. (B) Mean SYTOX Green fluorescence within three-dimensionally segmented nuclei. Cells were incubated with 10 μM SYTOX Green for 30 min, in the absence or presence of 5 μM tantalosin. Each point represents one nucleus; black horizontal lines indicate medians. The numbers were: U2OS, 534 control and 536 tantalosin-treated, from four independent experiments; SVG-A, 578 control and 562 tantalosin-treated, from four independent experiments; and SUM159, 113 control and 246 tantalosin-treated, from two independent experiments. The percentage of SYTOX-positive cells are shown in brackets of each graph. Cells were scored as positive when their mean nuclear fluorescence exceeded the mean of the corresponding control by more than one SD. Statistical significance: p>0.05 (ns); p<0.001 (***). **(C)** Mean SYTOX Green fluorescence within three-dimensionally segmented nuclei. Cells were incubated with 5 μM SYTOX Green for 2 h, in the absence or presence of 5 μM tantalosin. Each point represents one nucleus; black horizontal lines indicate medians. The numbers of nuclei analyzed were: U2OS, 866 control and 792 tantalosin-treated, from four independent experiments; SVG-A, 837 control and 850 tantalosin-treated, from four independent experiments; and SUM159, 225 control and 289 tantalosin-treated, from one experiment. The percentage of SYTOX-positive cells are shown in brackets of each graph. Cells were scored as positive when their mean nuclear fluorescence exceeded the mean of the corresponding control by more than one SD. Statistical significance: p>0.05 (ns); p<0.001 (***). **(D)** Time course of nuclear SYTOX Green accumulation. Cells were pulsed with 10 μM SYTOX Green for 5 min and then chased and imaged from 10 to 160 min after the start of the pulse, in the absence or presence of 5 μM tantalosin. Points show the percentage of SYTOX-positive cells from two independent experiments. Cells were scored as positive when their mean nuclear fluorescence exceeded the mean of the untreated 10-min control by more than one SD.

ASO delivery showed the same dependence on trafficking stage. We measured nuclear Alexa Fluor 647–ASO by LLSM calibrated for non-point-source fluorescence, using nuclear accumulation as a readout of prior escape from the endosomal lumen into the cytosol. A 30-min co-incubation with 4 μM ASO and 5 μM tantalosin, followed by a 30-min chase, increased nuclear ASO fluorescence approximately twofold relative to ASO alone (Fig. 6A-C). A 2-h co-incubation with 1 μM ASO and 5 μM tantalosin, followed by a 2-h chase, yielded an approximately threefold increase (Fig. 6A–C; Supplementary Fig. S13A-B). In contrast, tantalosin produced no detectable increase when added after a 2-h pulse with 1 μM ASO and an overnight chase into terminal lysosomes (Supplementary Fig.S13C,D). These measurements place the tantalosin-sensitive step in ASO escape early in endocytic trafficking.

**Figure 6.**
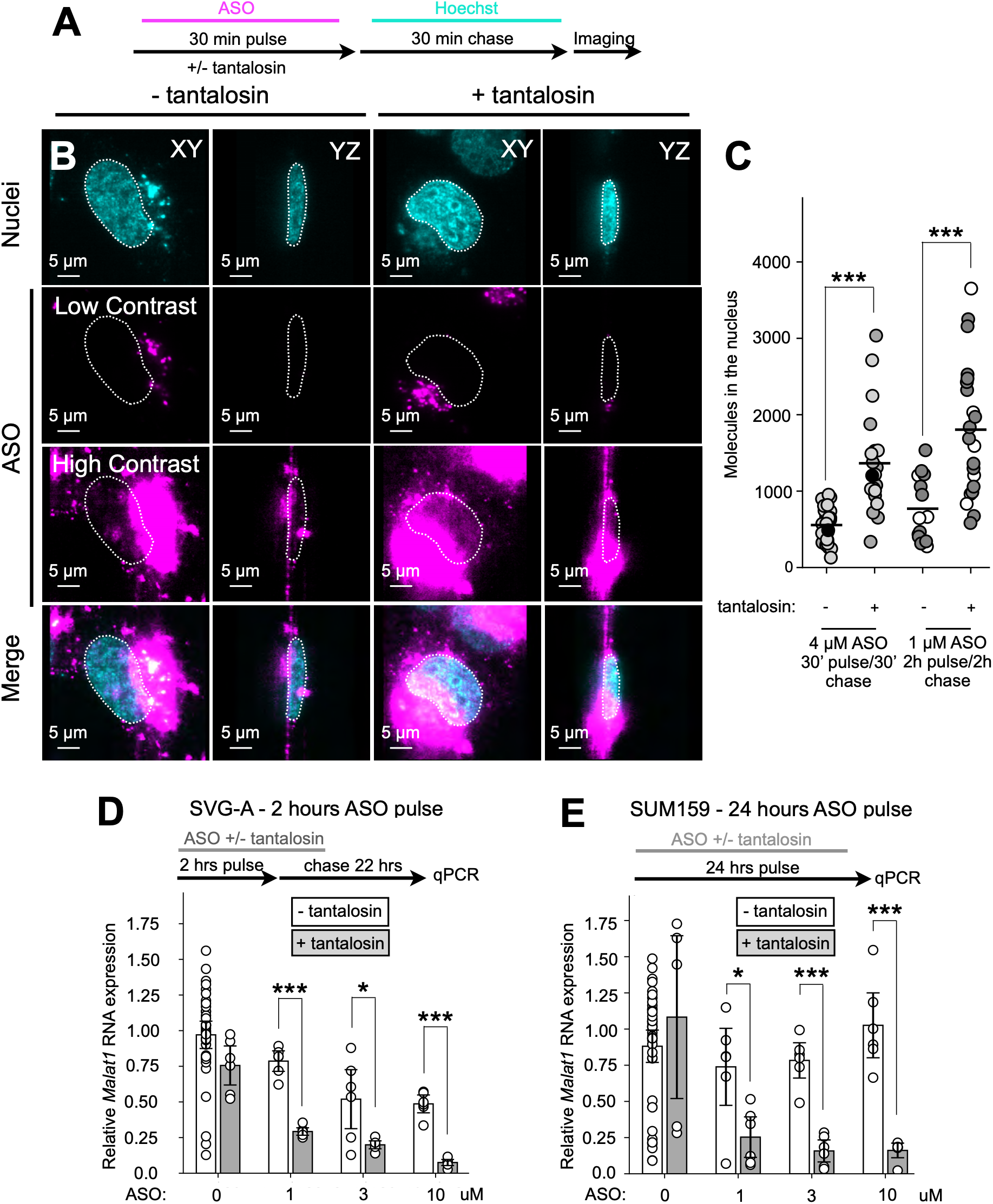
Tantalosin promotes ASO escape during the first hour of endocytic trafficking and enhances functional delivery (A) Experimental protocol. Cells were incubated with 4 μM ASO for 30 min in the absence or presence of 5 μM tantalosin, chased for 30 min with Hoechst labeling, and then imaged live. **(B)** Representative single optical sections acquired by MOSAIC lattice light-sheet microscopy from wild-type SVG-A cells labeled with JF549–Hoechst (cyan) and incubated for 30 min with Alexa Fluor 647–ASO (magenta), in the absence or presence of 5 μM tantalosin. Alexa Fluor 647–ASO is displayed at increased contrast to reveal the nuclear signal; fluorescence from endosomal ASO is therefore saturated. White dotted lines outline the nuclei. Corresponding XY and YZ views are shown. Scale bar, 5 μm. **(C)** Number of ASO molecules per nucleus after a 30-min pulse with 4 μM ASO followed by a 30-min chase, or after a 2-h pulse with 1 μM ASO followed by a 2-h chase, in the absence or presence of 5 μM tantalosin. Each point represents one nucleus; the horizontal line indicates mean. Numbers of nuclei analyzed were: 30-min pulse, 24 control and 18 tantalosin-treated, 2-h pulse, 14 control and 22 tantalosin-treated. Data shown are representative of two independent experiments. **(D)** Effect of tantalosin on the functional activity of ASOs in SVG-A cells. Cells were incubated for 2 h with increasing concentrations of *Malat1* ASO, in the absence or presence of 5 μM tantalosin, and then chased for 22 h. *Malat1* RNA abundance was measured by quantitative PCR, normalized to cyclophilin, and expressed relative to the corresponding untreated control. Each point represents one biological replicate; black lines indicate means and error bars indicate SEM. Data are from at least six biological replicates collected in two independent experiments. **(E)** Effect of tantalosin on the functional activity of ASOs in SUM159 cells. Cells were incubated for 24 h with increasing concentrations of *Malat1* ASO, in the absence or presence of 5 μM tantalosin. *Malat1* RNA abundance was measured and plotted as in D. Each point represents one biological replicate; black lines indicate means and error bars indicate SEM. Data are from at least six biological replicates collected in two independent experiments.

The increase in nuclear ASO accumulation was accompanied by enhanced activity. In SVG-A cells, a 2-h co-incubation with ASO and tantalosin followed by a 22-h chase increased *Malat1* RNA knockdown at every ASO concentration tested (Fig. 6D). SUM159 cells, which showed little or no response to ASO alone at 24 h (Supplementary Fig. S1), showed *Malat1* knockdown at all tested concentrations after 24-h co-incubation with tantalosin (Fig. 6E), but not after 2-h co-incubation (Supplementary Fig. S13E). Thus, tantalosin enhanced both nuclear delivery and target suppression, with the magnitude and kinetics of the response depending on cell type.

Together, the SYTOX Green, nuclear ASO and *Malat1*-suppression measurements show that productive ASO delivery can be enhanced by perturbing an early endocytic step. While our results are consistent with prolonged membrane openings, they do not establish delayed repair as the mechanism. Testing the complementary prediction that enhanced repair suppresses escape awaits identification of an intervention that selectively promotes membrane repair in early endosomes.

### G3BP1/2 loss modestly enhances functional delivery without overt endolysosomal rupture or stress-granule dependence

Stress granules are membraneless ribonucleoprotein condensates that assemble in the cytosol in response to diverse stressors (81, 82). In macrophages, Bussi et al. reported that phagosome damage by internalized pathogens triggers local stress-granule assembly and recruits the core stress-granule protein G3BP1 at membrane injury sites, where it contributes to membrane plugging and repair (38). This model predicts that loss of G3BP1/2 could enhance functional delivery of internalized ASOs by permitting greater escape across damaged endolysosomal membranes.

We tested whether G3BP1/2 limits ASO delivery by promoting endolysosomal membrane repair, using U2OS cells for which a G3BP1/2 double-knockout line is available (62). A study published while our work was in progress reported that CRISPR-mediated G3BP1/2 double knockout and ISRIB treatment enhanced ASO-mediated *Kras* knockdown in KPC pancreatic cancer cells. The authors attributed these effects to impaired stress-granule-mediated repair of ASO-containing, galectin-9-positive endosomes near the nucleus (39).

In U2OS cells, G3BP1/2 loss caused a modest shift toward higher endolysosomal pH but did not produce overtly neutral endosomes or lysosomes compared to the parental U2OS wild-type-cells, as measured with an internalized dextran-based ratiometric sensor (Fig. 7A–D; Supplementary Fig. S14F). LysoTracker Red staining gave a consistent result, and ectopic eGFP–G3BP1 expression reversed the modest pH shift (Supplementary Fig. S14A–E). We detected neither increased dextran release nor galectin-3 puncta in knockout cells, whereas LLOMe induced substantial galectin-3 recruitment (Fig. 7E–G). Thus, G3BP1/2 loss did not cause detectable basal endolysosomal membrane damage in this system.

**Figure 7.**
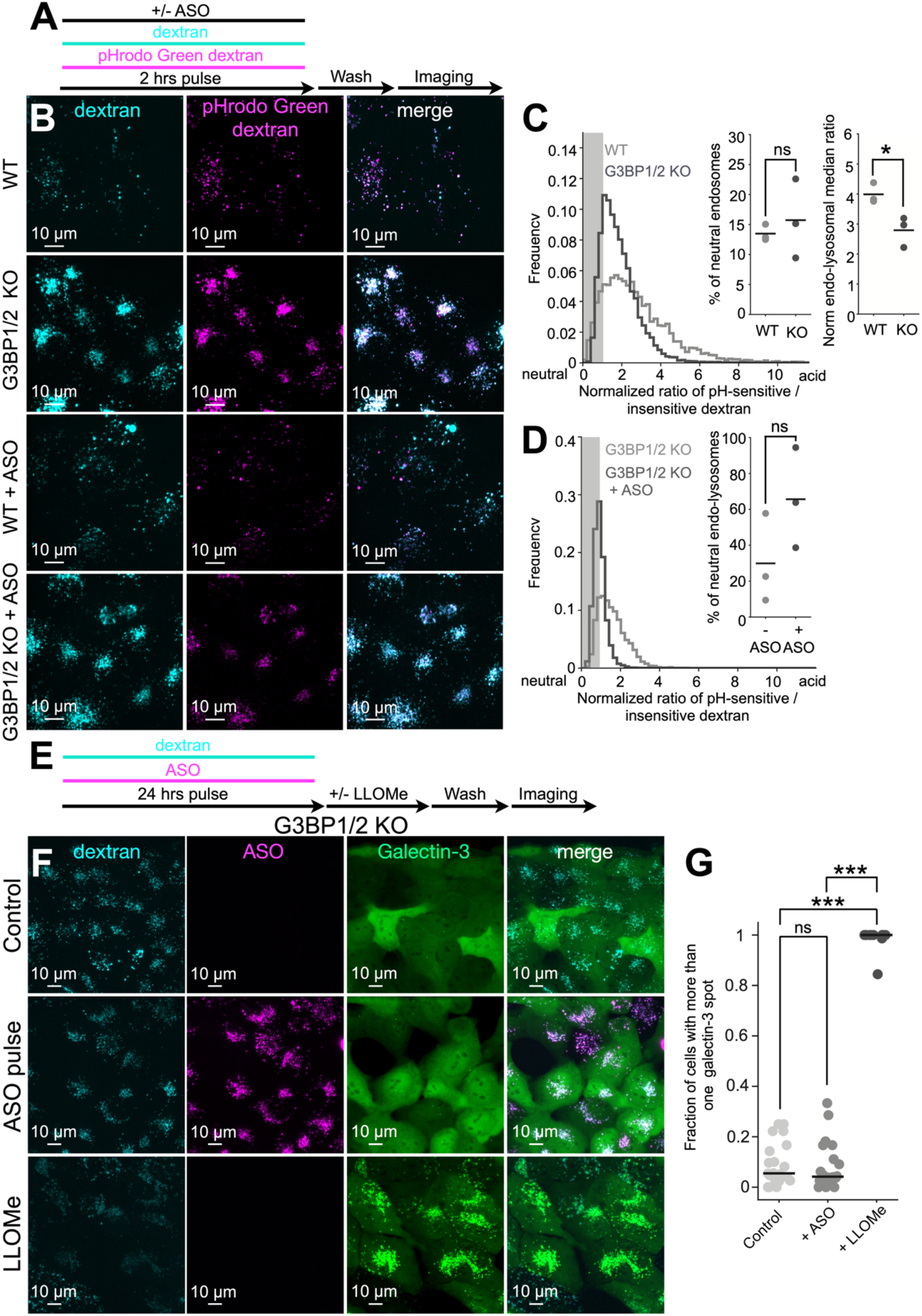
G3BP1/2 loss does not cause endolysosomal membrane damage.(A) Experimental protocol. 8 μM Alexa Fluor 647-dextran (pH-insensitive) and 8 μM pHrodo Green-dextran (pH-sensitive) were co-pulsed for 2-hours in U2OS cells expressing or not G3BP1/2 in the presence or absence of 1 μM non-fluorescent ASO, washed, and imaged. **(B)** Representative maximum-projection spinning-disc images of wild-type U2OS cells and G3BP1/2 knockout cells labeled for 2-hours with Alexa Fluor 647-dextran (cyan) and pHrodo Green-dextran (magenta), with or without ASO. Scale bar, 10 μm. **(C)** Quantification of endolysosomal pH in wild-type and G3BP1/2 knockout cells from **(B)**, calculated from the ratio of pHrodo Green-dextran to Alexa Fluor 647-dextran fluorescence and normalized to the mean + 1 standard deviation of the pH 7.4 calibration value. Frequency plots show distributions of the pH-sensitive: pH-insensitive fluorescence ratio from three biological replicates, with 10 fields of view per replicate, for wild-type cells (# endosomes =16,660) and G3BP1/2 knockout cells (# endosomes = 25,693). The shaded region denotes the mean + 1 standard deviation of the pH 7.4-clamped ratio. The left inset shows the fraction of neutral endosomes, defined as compartments with ratios within the shaded region. The right inset shows the median normalized ratio for each condition; each dot represents an independent biological replicate. Statistical significance: p>0.05 (ns); p<0.05 (*). **(D)** Quantification of endolysosomal pH in G3BP1/2 knockout cells in the presence or absence of 1 μM ASO from **(B)**, calculated from the ratio of pH-sensitive to pH-insensitive dextran fluorescence and normalized to the mean + 1 standard deviation of the pH 7.4 calibration value. Frequency plots show distributions of the pH-sensitive: pH-insensitive fluorescence ratio from three biological replicates, with 10 fields of view per replicate, for untreated G3BP1/2 knockout cells (# endosomes = 24,989) and ASO-treated G3BP1/2 knockout cells (# endosomes = 27,451). The shaded region denotes the mean + 1 standard deviation of the pH 7.4-clamped ratio. The inset shows the fraction of neutral endosomes in G3BP1/2 knockout cells in the presence or absence of ASO. Each dot represents an independent biological replicate. Statistical significance: p>0.05 (ns). **(E)** Experimental protocol. 8 μM Alexa Fluor 568-dextran and 1 μM Alexa Fluor 647-ASO were co-pulsed for 24-hours and then incubate in the presence or absence of 0.5 mM LLOMe before imaging. **(F)** Representative maximum-projection spinning-disc images of G3BP1/2 knockout U2OS cells stably expressing eGFP-galectin-3 (green), untreated or treated with 1 μM Alexa Fluor 647-ASO (magenta) for 24-hours or with 0.5 mM LLOMe for 30 min. Alexa Fluor 568-dextran is shown in cyan. Scale bar, 10 μm. Data are representative of two independent experiments. **(G)** Quantification of **(F)**, shown as the fraction of cells with one or more cytoplasmic galectin-3 puncta. The median is shown as a black line; each dot represents one field of view. Data are from two independent experiments. Statistical significance: p>0.05 (ns); p<0.001 (***).

We also examined whether eGFP–G3BP1 localized to damaged endolysosomal membranes. Brief treatment with 0.5 mM LLOMe, sufficient to induce dextran leakage(82, 83), did not recruit eGFP–G3BP1 to dextran-labeled endolysosomes (Supplementary Fig. S15). At 4 mM LLOMe, eGFP–G3BP1 formed cytosolic puncta that remained separate from the labeled endolysosomes. Cells also rounded slightly, consistent with a generalized stress response. These puncta therefore appeared more consistent with stress granules than with assemblies at membrane lesions. Ectopic eGFP–G3BP1 restored arsenite-induced stress-granule formation in the knockout cells (Supplementary Fig. S16A,B), but ASO treatment induced no eGFP–G3BP1 puncta in the cytosol or on ASO-containing endolysosomes (Supplementary Fig. S15C). Under these conditions, eGFP–G3BP1 did not mark damaged endolysosomes or putative ASO escape sites.

Despite the absence of increased membrane damage or dextran release, G3BP1/2 loss modestly enhanced ASO activity. After 24 h of continuous incubation, *Malat1* RNA measurements by qPCR indicated approximately two-to threefold greater ASO activity in knockout cells than in controls. Ectopic eGFP–G3BP1 expression restored activity to control levels (Fig. 8A). Unlike the response reported in KPC cells, ISRIB did not enhance ASO activity in U2OS cells. G3Ia likewise had no effect, although both compounds inhibited stress-granule formation (84, 85)(Supplementary Fig. S16A–D; Supplementary Video S6). The enhanced activity in knockout cells therefore did not require loss of stress-granule assembly. It also occurred despite slightly lower bulk uptake of ASO and dextran (Supplementary Fig. S17A–F) and more rapid delivery of both cargos to late endosomes, endolysosomes and terminal lysosomes (Fig. 8C-F; Supplementary Videos S7 and S8), where ASOs remained largely sequestered.

**Figure 8.**
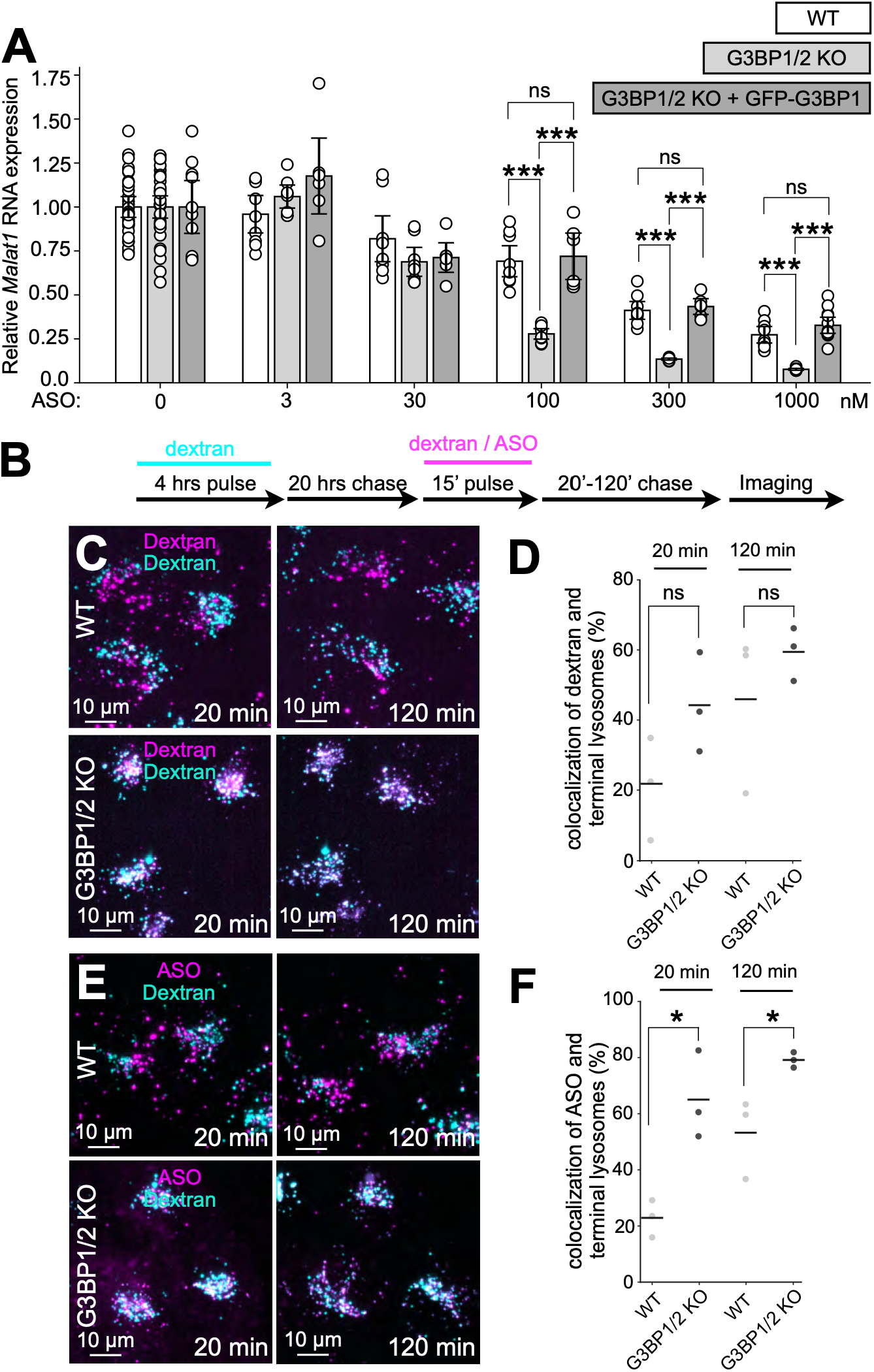
G3BP1/2 knockout increases ASO activity and accelerates ASO delivery to lysosomes in U2OS cells.(A) Effect of anti-Malat1 ASO treatment for 24-hours on *MALAT1* RNA levels in wild-type U2OS cells (white), G3BP1/te2 knockout U2OS cells (light gray) and G3BP1/2 knockout U2OS cells stably expressing eGFP-G3BP1 (dark gray). RNA abundance was measured by quantitative PCR, normalized to cyclophilin, and plotted as fold change relative to the corresponding untreated control. Black lines indicate the mean and error bars indicate SEM. Data are from at least nine biological replicates from three independent experiments. Statistical significance: p>0.05 (ns); p<0.001 (***). **(B)** Experimental protocol. 8 μM Alexa Fluor 568-dextran (cyan) was pulsed for 4-hours and chased for 20-hours to label terminal compartments, followed by a 15 min pulse with 8 μM Alexa Fluor 647-dextran (magenta) or 1 μM Alexa Fluor 647-ASO (magenta). Cells were imaged after a 20 min or 120 min chase. **(C)** Representative maximum-intensity projections acquired by lattice light-sheet microscopy (Zeiss) showing Alexa Fluor 647–dextran (magenta) after a 15-min pulse and a 20-or 120-min chase in wild-type and G3BP1/2-knockout U2OS cells. Cells were preloaded with Alexa Fluor 568–dextran and chased to label terminal lysosomes (cyan). **(D)** The percentage of Alexa Fluor 647–dextran-positive endosomes colocalizing with terminal lysosomes was slightly higher in knockout cells at both chase times, but neither difference was statistically significant. Numbers of endosomes analyzed: wild type, 7,751 at 20 min and 9,064 at 120 min; knockout, 14,597 at 20 min and 15,136 at 120 min. Data are from three biological replicates. Scale bar, 10 μm. Statistical significance: p > 0.05 (ns). **(E)** Representative maximum-intensity projections acquired by lattice light-sheet microscopy (Zeiss) showing Alexa Fluor 647–ASO (magenta) after a 15-min pulse and a 20-or 120-min chase in wild-type and G3BP1/2-knockout U2OS cells. Cells were preloaded with Alexa Fluor 568– dextran and chased to label terminal lysosomes (cyan). **(F)** The percentage of Alexa Fluor 647–ASO-positive endosomes colocalizing with terminal lysosomes was significantly higher in knockout cells than in wild-type cells. Numbers of endosomes analyzed: wild type, 7,388 at 20 min and 8,323 at 120 min; knockout, 14,815 at 20 min and 16,014 at 120 min. Data are from three biological replicates; a subset is also shown in Fig. 1F. Scale bar, 10 μm. Statistical significance: p < 0.05 (*).

G3BP1/2 loss did not alter endolysosomal compartments (Supplementary Fig. S18) but caused an unexpected reorganization of the endoplasmic reticulum we noticed by volume EM imaging using focused ion beam scanning electron microscopy (FIBSEM). Ribosome-bearing regions appeared expanded relative to the corresponding compact regions in parental cells and were connected by extended, narrow membrane profiles lacking detectable ribosomes. Parental cells showed abundant ribosomes across both the expanded regions and their connecting membranes. These changes affected extensive portions of the ER, although we did not quantify their prevalence. Ectopic G3BP1 expression rescued the altered architecture.

Thus, the two cell systems agree that G3BP1/2 loss can enhance ASO activity, but our U2OS measurements do not support increased endolysosomal leakage or impaired stress-granule-mediated repair as the explanation. The accompanying changes in cargo trafficking and ER organization leave open an indirect effect of G3BP1/2 loss on productive ASO delivery.

## DISCUSSION

Our results indicate that productive ASO escape occurs early during endocytic trafficking, most likely during passage through early endosomes toward the recycling pathway. Tantalosin enhanced escape during this early phase, increasing both nuclear ASO accumulation and functional delivery. Most internalized ASOs nevertheless continue their traffic and accumulated in late endosomes, endolysosomes, and terminal lysosomes, where they remained largely sequestered even when the limiting membrane became permeable to soluble luminal cargo. The timing of escape and the subsequent retention of ASOs therefore distinguish productive delivery from bulk cellular uptake.

Previous studies have measured nuclear ASO after gymnotic uptake by fluorescence imaging or analysis of isolated nuclei, with results that depend on the experimental system and measurement method (6, 18, 64, 86). Buntz et al. (2019) (85) estimated that approximately one-quarter of cellular ASO reached the nucleus, compared with approximately 4% in our measurements. (86) Two distinctions matter when interpreting these estimates. First, correlative NanoSIMS and electron microscopy detected ASO along the inner nuclear membrane as well as within the nucleoplasm (6). Measurements of isolated nuclei can therefore include membrane-associated ASO even when cytoplasmic contamination is carefully controlled, as in Pendergraff et al. (2020) (64). Such measurements need not represent the amount present in the nucleoplasm. Second, we identified an optical contribution from the abundant ASO bound at or near the coverslip. Fluorescence from this material extends into nuclear optical sections through the microscope point-spread function and can inflate the apparent nuclear signal. This contribution affects wide-field and conventional confocal imaging and remains detectable, although reduced, in LLSM. We corrected for it under our acquisition conditions, approximately 2.5 μm above the glass. Uncorrected coverslip fluorescence could contribute to differences among imaging-based estimates, alongside differences in ASO chemistry, exposure and cell type. Our corrected estimate of approximately 4% is consistent with the predominantly endolysosomal distribution resolved by NanoSIMS and provides a well-supported measure of nuclear accumulation under the conditions examined here.

The behavior of the much larger ASO pool in late compartments indicates that sequestration involves more than an intact membrane barrier. ASO accumulation shifted luminal pH toward neutrality, but even subsequent LLOMe treatment sufficient to recruit galectin-3 and release 10-kDa dextran produced neither detectable diffuse cytosolic ASO fluorescence nor additional nuclear accumulation. The estimated hydrodynamic radius of a 20-nucleotide single-stranded oligonucleotide is approximately 1.6 nm, compared with approximately 2.3 nm for 10-kDa dextran (66, 87). Pore size alone therefore cannot explain why dextran escaped while ASO remained. Lysosome fusion with the plasma membrane likewise released dextran while retaining ASO, showing that retention did not depend on LLOMe or on the particular process that opened the compartment.

This retention extended beyond the discrete ASO foci. Terminal lysosomes, whose larger size allowed us to resolve their contents without apilimod, contained both foci and a broadly distributed ASO signal. Apilimod-expanded compartments showed similar distributions, whereas dextran filled the lumen uniformly (71). In one spontaneous event captured serendipitously during three-dimensional live imaging (Supplementary Video S4), an endolysosome lost its dextran signal and contracted while retaining both an ASO focus and the surrounding distributed signal, with no detectable change in total ASO fluorescence. Thus, a uniform appearance did not identify a freely releasable ASO pool. The constrained motion of foci labeled with either Alexa Fluor 647 or Janelia Fluor 646 provided complementary evidence for association with endolysosomal constituents, although these trajectories describe only the resolved foci. Together, the release and tracking measurements support retention through association with luminal components, the limiting membrane, or both.

The molecular basis of this association remains unresolved. Individual compartments contained thousands of ASO molecules, suggesting that retention may involve multivalent interactions with an extended luminal or membrane-associated assembly. One-to-one binding to a few low-abundance proteins would be insufficient to accommodate such loads. Although quantitative cryo-ET has reported low copy numbers of several lysosomal membrane-associated proteins (88), those measurements do not establish the total binding capacity of the compartment. Identifying the constituents that retain ASOs will therefore require direct analysis of these associations.

Our observations differ from a prevalent view in the ASO literature, which proposes that productive release occurs from late endosomes or multivesicular bodies, in some cases through mechanisms linked to back-fusion of intraluminal vesicles, LBPA-containing membranes, or other endosomal factors (28, 89, 90). A simple ILV back-fusion model is particularly difficult to reconcile with endosomal topology. Although back-fusion could release ILV luminal contents into the cytosol, ASOs internalized into an endosome or multivesicular body reside in the organelle lumen, outside the ILVs. Back-fusion alone would therefore leave these ASOs within the organelle. Our results instead favor escape before ASOs encounter the strongly retentive environment of late endosomes and lysosomes. Early and recycling endosomes are plausible sites because they undergo frequent fusion and fission and extensive membrane remodeling (29, 31, 91). We propose that transient membrane discontinuities generated during this remodeling permit a small fraction of internalized ASOs to escape before extensive association with endosomal contents. Only those molecules that encounter an opening before it closes would gain cytosolic access; the remainder would continue toward sequestration in later compartments. Tantalosin could increase escape by prolonging these openings, as proposed for interference with IST1-associated membrane repair. Its effects establish that early escape can be enhanced, however, without yet defining the membrane lesion or its repair mechanism. This interpretation is consistent with reports that perturbations that prevent or delay progression to later endosomal compartments can enhance ASO activity (28, 71, 89, 90). Depletion of EEA1, which is required for early endosome biogenesis, or interference with Rab5C, which contributes to early-to-late endosomal homeostasis, impairs productive ASO delivery (89), whereas loss of the clathrin adaptor subunit AP1M1, which functions in traffic between endosomes and the trans-Golgi network, enhances it (28). Productive delivery also increases in response to SH-BC-893, which pharmacologically inactivates ARF6 and inhibits both early endosomal recycling and trafficking to later compartments (90). These perturbations do not identify the site of membrane passage, but they support a model in which delayed progression can prolong access to an early compartment permissive for escape. A small fraction of lysosome-retained ASO might eventually be released into the extracellular space during lysosome exocytosis and taken up again by endocytosis. Reinternalization would provide another opportunity for escape during passage through early or recycling endosomes. Such repeated uptake could contribute to sustained ASO activity in vivo, although this possibility remains to be tested.

Our cell-culture findings provide a framework for considering how ASOs might reach their targets in vivo. An initial wave of productive escape to cytosol might accompany ASO arrival in the interstitial fluid and uptake by target cells. Later, occasional fusion of ASO-containing lysosomes with the plasma membrane might release a fraction of their retained cargo. After local dispersal, these ASOs could be internalized by neighboring cells and gain another opportunity to escape during early or recycling endosomal trafficking. This latter possibility remains speculative. Our exocytosis experiments showed substantial ASO retention, and neither extracellular redistribution nor subsequent productive uptake has yet been measured in vivo.

Histological studies have mapped ASO accumulation in individual cells in brain, liver, and kidney (92–94), and single-nucleus transcriptomics has documented cell-type-specific target engagement in the brain (95). In these studies, however, punctate, lysosome-associated ASO and functional target modulation were not measured in the same individual cells. The proposed second route therefore makes a testable prediction: cells with large lysosomal ASO stores should lie near cells with target modulation but little detectable lysosomal ASO. Multiplex spatial measurements of total ASO, lysosomal ASO, and target RNA will be required to test this prediction.

We therefore propose two kinetically and mechanistically distinct fates for internalized ASO (Fig. 9). A small productive fraction escapes early, perhaps from early or recycling endosomes, before extensive association with endosomal contents. The much larger nonproductive fraction traffics onward to late endosomes, endolysosomes, and terminal lysosomes, where it remains retained even when these compartments neutralize or become permeable to soluble cargo. Future work should define the site of productive escape, identify the interactions that retain ASOs in late compartments, and determine whether either process can be manipulated to improve delivery. The approximately two-to threefold increase in nuclear ASO accumulation with tantalosin demonstrates that early escape can be enhanced pharmacologically. Understanding what limits this event, and what prevents release, thereafter, should guide further improvements in productive ASO delivery.

**Figure 9.**
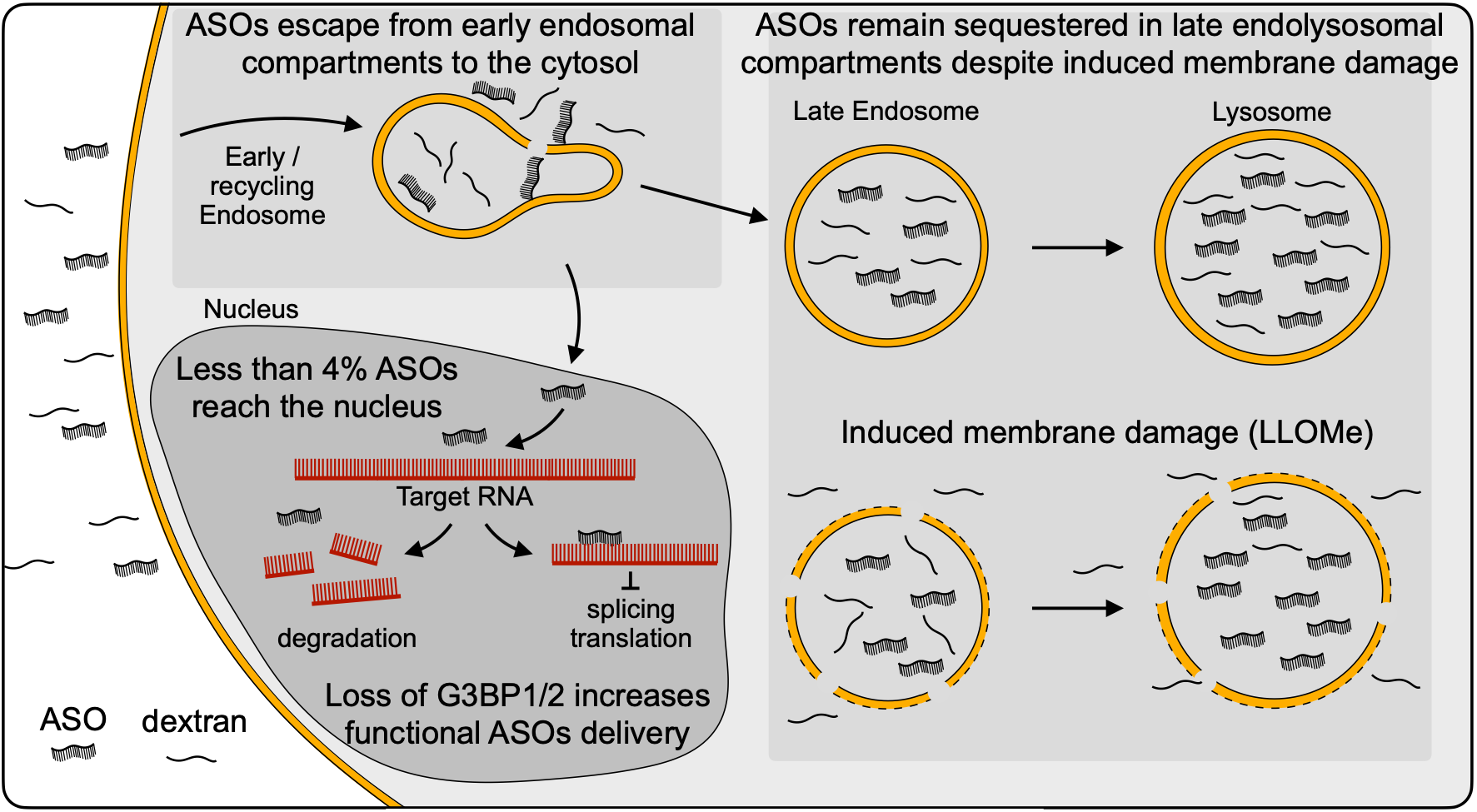
Proposed model. Productive escape of unmodified ASOs occurs early after endocytosis, most likely from early or recycling endosomes and before extensive association with endosomal contents. Less than 4% of the internalized ASO reaches the nucleus and modulates its RNA target. The much larger nonproductive fraction continues to late endosomes, endolysosomes, and lysosomes, where it remains sequestered and shows little release even after overt limiting membrane damage. Loss of G3BP1/2 does not substantially perturb the endolysosomal system but increases functional ASO delivery.

## ACKNOWLEDGMENTS

We thank Ricardo F Bango Da Cunha Correia for providing training in lattice light-sheet microscopy, Jose Inacio Costa-Filho for assistance denoising the FIB-SEM images, Beren Aylan and members of the Kirchhausen laboratory for support, encouragement and helpful discussions. We thank Anwesha Sanyal for assistance with the initial stages of gene editing and the Andersen laboratory for providing wild-type and G3BP1/2-knockout U2OS cells. We thank Yao-Wen Wu for providing an aliquot of tantalosin as a reference standard. We also acknowledge the L. Lavis lab and the Open the Chemistry team (Janelia Research Campus, HHMI) for the generous gift of Hoechst Janelia Fluor 549. C. Frank Bennett is the executive vice president and chief scientific officer at Ionis Pharmaceuticals. The authors declare no competing financial interests.

## AUTHOR CONTRIBUTIONS

A. Saminathan, E. Sitarska and T. Kirchhausen conceptualized and designed the experiments; A. Saminathan and E. Sitarska carried out the cell biological experiments and imaging; E. Sitarska and E. Somerville prepared the samples for FIB-SEM. E. Somerville collected and processed the FIB-SEM volumetric data. A. Saminathan, E. Sitarska and G. Scanavachi carried out the optical image analysis. Patrick Stock developed the analytical pipeline used to quantify nuclear SYTOX Green fluorescence. Pengxiang Shen in Daniel Kahne’s laboratory synthesized tantalosin. E. Sitarska, M. F. Courtney and D. A. Reid conducted and/or supervised the RT-PCR experiments and C. F. Bennett contributed to the conceptual design of these experiments. M. B. Danielsen, F. K. N. Davidsen generated Janelia Fluor 646-ASO with the supervision of K. J. Jensen. T. Kirchhausen edited the manuscript in close consultation with A. Saminathan and E. Sitarska, with final input from all co-authors.

## AI DISCLOSURE STATEMENT

During the preparation of this work the author(s) used ChatGPT to improve readability and language of the work. After using this tool/service, the author(s) reviewed and edited the content as needed and take(s) full responsibility for the content of the publication.

We also used ChatGPT and CODEX (OpenAI) to develop analysis scripts and an interactive HTML viewer for examining lysosomal cargo release following ionomycin treatment. The scripts linked endolysosomal objects across successive frames by position, generated fluorescence traces, and identified abrupt decreases in dextran fluorescence for inspection alongside the corresponding ASO signal. We reviewed the edited text and validated each candidate release event by visual inspection. The authors retain responsibility for the analyses, interpretation and final manuscript.

## DATA, SOFTWARE and CODE AVAILABILITY

Fiji macros, MATLAB and Python scripts used for colocalization analysis, pH-ratio measurements and ASO-uptake quantification were described previously (52) and are available in the Kirchhausen GitHub repository https://github.com/kirchhausenlab/pikfyve_endosome_fusion_mechanics, https://github.com/kirchhausenlab/aso_early_endocytic_trafficking. The qPCR data and a movie with a 3D FIB-SEM rendition illustrating a representative example of collapsed ER in a U2OS lacking G3BP1/2 of are available in ZENODO https://doi.org/10.5281/zenodo.23219586. The complete 3D FIB-SEM images are available at the AWS Bucket: https://kirchhausenlab.github.io/tklab_data_explorer/. An online publicly available tool based on Neuroglancer has been created to view the 3D volumes relevant to this study and others. Because of the large number and combined size (∼500 GB) of the fluorescence image files underlying this study, deposition in a public repository is impractical. The files are available from the corresponding author upon reasonable request.

