## Supplementary Information for "Naked antisense oligonucleotides access the cytosol during early endocytic trafficking"

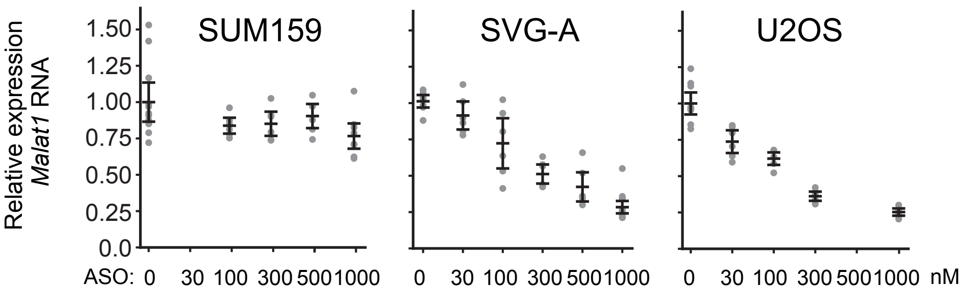


### Supplementary Figure S1

**Supplementary Figure S1. Cell-type-dependent activity of naked ASO against *Malat1***

SUM159, SVGA, and U2OS cells were incubated for 24 h with increasing concentrations of *Malat1*-targeting ASO. RNA abundance was measured by quantitative PCR, normalized to cyclophilin, and plotted as fold change relative to the corresponding untreated control. Each dot represents a replicate; black lines indicate the mean and error bars indicate SEM. Data are from at least six biological replicates across two independent experiments.

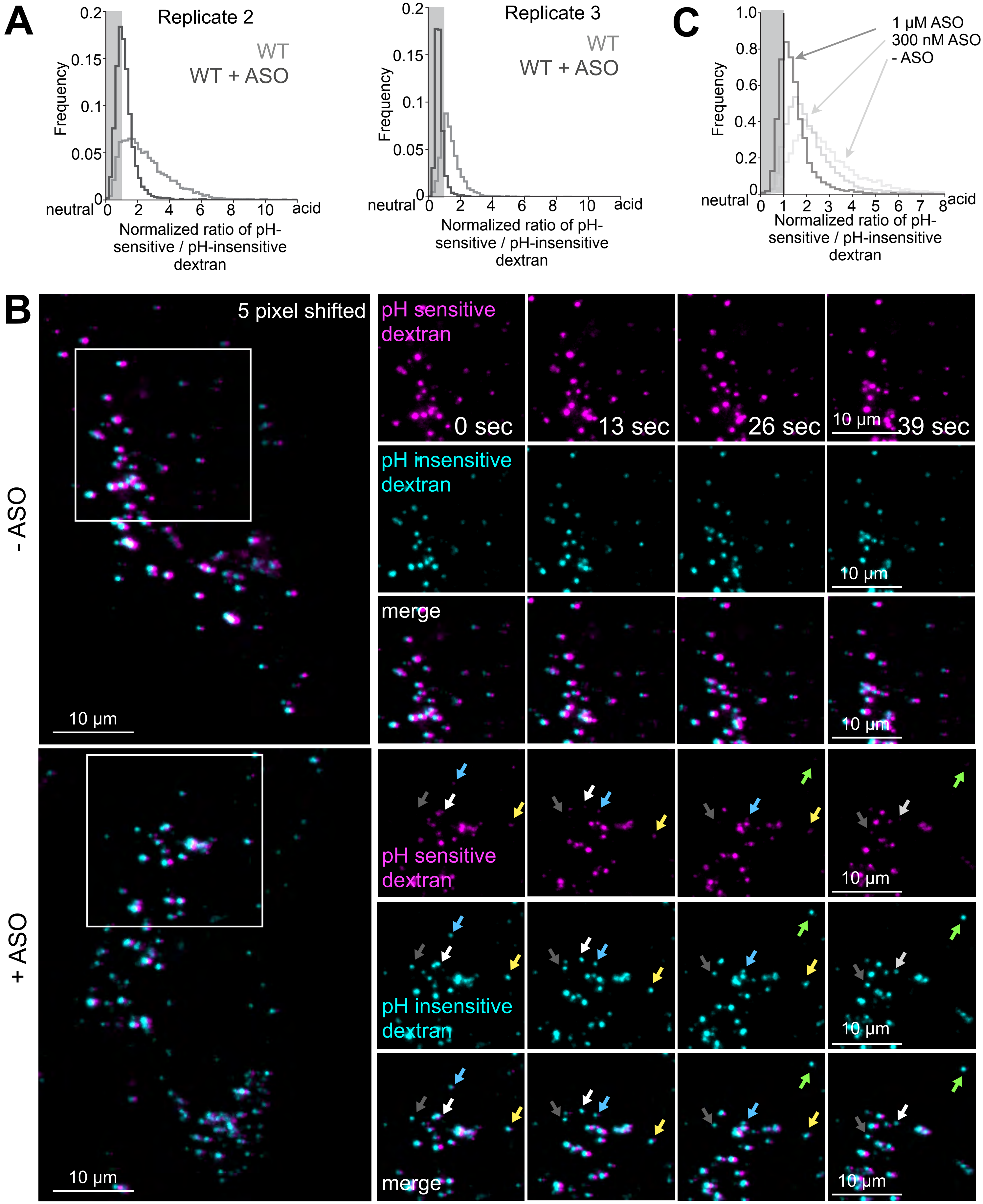


### Supplementary Figure S2

**Supplementary Figure S2. Related to Figure 2.**

**(A)** Quantification of endolysosomal pH in wild-type U2OS cells in the absence or presence of 1 μM non-fluorescent ASO (replicates of Fig. 2 B and C), calculated from the ratio of pH-sensitive pHrodo Green-dextran to pH-insensitive Alexa Fluor 647-dextran fluorescence and normalized to the mean + 1 SD of the pH 7.4-calibrated fluorescence. Frequency plots show the distribution of the pH-sensitive: pH-insensitive fluorescence ratio from one of three biological replicates, with 10 fields of view per replicate, for wild-type U2OS cells (total # endosomes = 15,476 (- ASO) and 16,093 (+ASO)). The shaded region denotes the mean + 1 SD (84th percentile) of the pH 7.4-clamped ratio.

**(B)** Representative 3D volumetric lattice light-sheet microscopy (MOSAIC) images of Alexa Fluor 647-dextran (cyan) and pHrodo Green-dextran (magenta) in the absence (upper) or presence (lower) of ASO. Images were offset by 5 pixels to aid visualization of colocalization. Colored arrows indicate the same endolysosomes over time. The volumetric 3D time series, acquired every 2.6 sec, is displayed as maximum-intensity projections. Scale bar, 10 μm.

**(C)** Quantification of endolysosomal pH in SVG-A wild-type cells, calculated using the ratio of pH-sensitive pHrodo Green-dextran to pH-insensitive dextran fluorescence normalized to mean + 1 standard deviation of pH 7.4 calibrated fluorescence. Frequency plots show the distribution of pH-sensitive/insensitive ratio from one biological replicate, 10 fields of view per replicate, (total # endosomes = 9,145 (- ASO), 9,589 (+ 300 nM ASO), and 8,597 (+ 1 μM ASO)). The shaded region indicates mean +1 standard deviation of the pH 7.4 clamped ratio of pH sensitive to insensitive fluorescence.

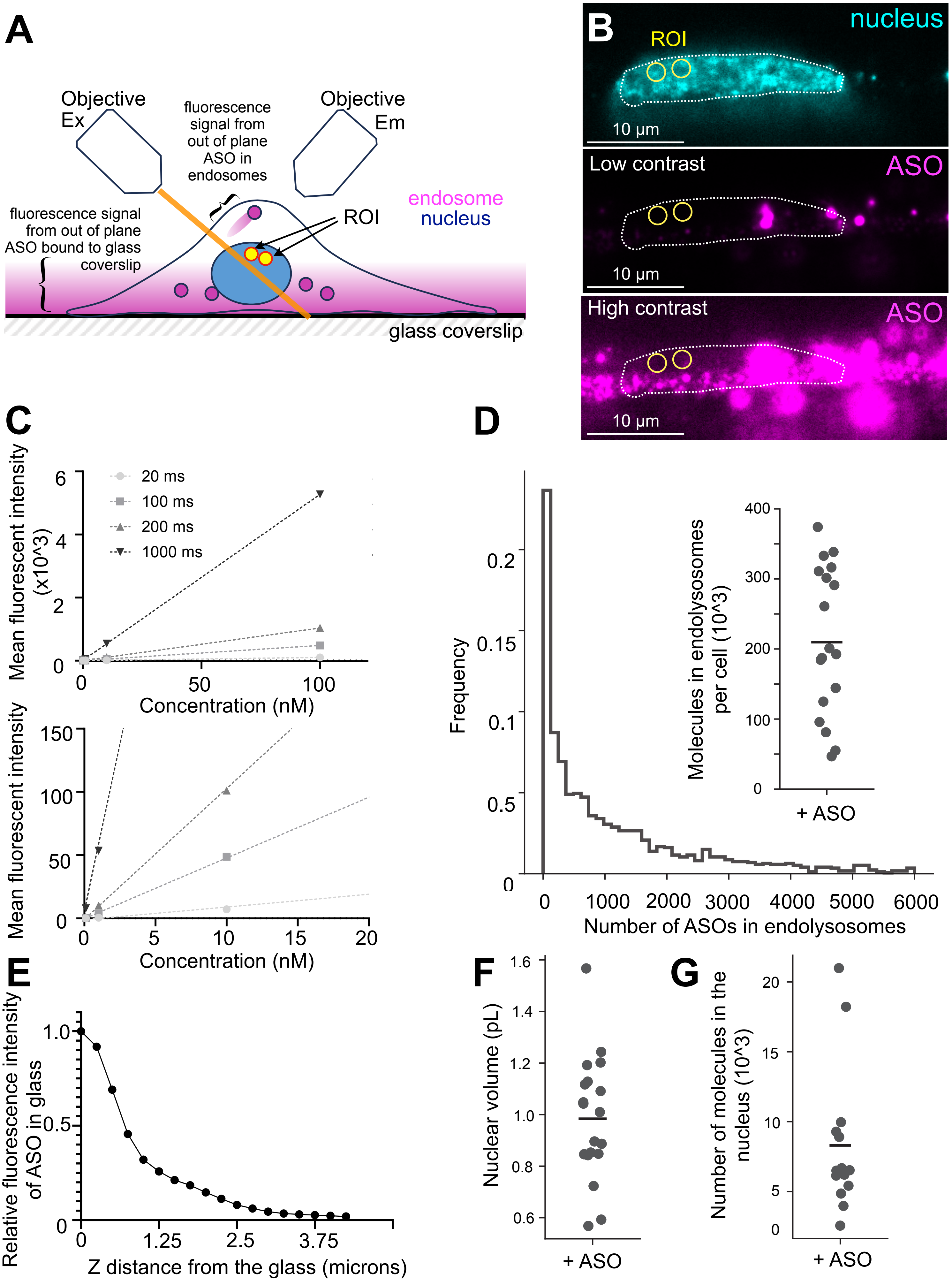


### Supplementary Figure S3

**Supplementary Figure S3. Related to Figure 2.**

**(A)** Schematic illustrating the selection of regions of interest (ROIs) within nuclei of wild-type U2OS cells incubated with 1 µM Alexa Fluor 647-ASO for 24 h, selected to minimize contributions from out-of-plane ASO signal arising from endolysosomes and from ASO bound to the glass coverslip.

**(B)** Representative single plane lattice light-sheet microscopy (MOSAIC) images of wild-type U2OS cells labeled with Hoechst JF 549 (cyan), incubated with Alexa Fluor 647-ASO (magenta) for 24 h. Alexa Fluor 647-ASO is shown at higher contrast on the left and lower contrast on the right. Scale bar, 10 µm.

**(C)** Representative quantification of mean Alexa Fluor 647 fluorescence intensity at the indicated concentrations in the MOSAIC lattice light-sheet imaging chamber. The lower panel shows the lower Alexa Fluor 647 concentrations from the upper panel. Linear regression was performed for the indicated exposure times. Data are representative of one of two independent experiments.

**(D)** Frequency distribution of the number of Alexa Fluor 647-ASO molecules in endolysosomes of wild-type U2OS cells after 24-hours incubation. Inset: number of endolysosomal ASO molecules per cell. Each dot represents one cell, and the line indicates the median (# cells = 19). Data are representative of two independent experiments.

**(E)** Line profile of relative ASO fluorescence intensity on the glass coverslip along the z-axis of the emission objective, as shown in **(A)**.

**(F)** Quantification of nuclear volume in wild-type U2OS cells (# nuclei = 19). Each dot represents one nucleus, and the line indicates the median. Data are representative of two independent experiments.

**(G)** Number of ASO molecules per nucleus. Each dot represents one nucleus, and the line indicates the median (# nuclei = 14). Data are representative of two independent experiments.

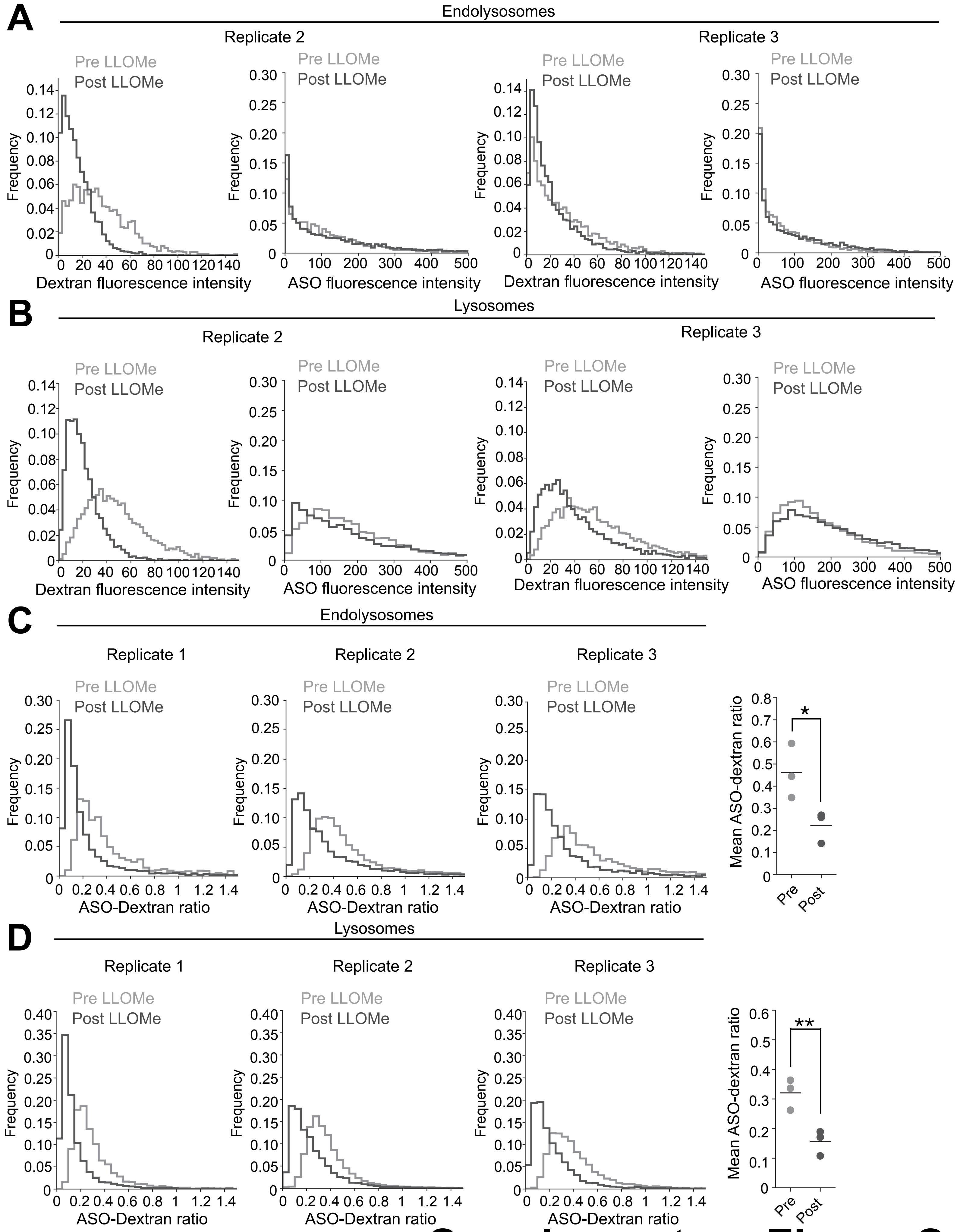


Supplementary Figure S4

**Supplementary Figure S4. Related to Figure 3.**

**(A)** Quantification of replicates from Fig. 3B, C, shown as frequency plots of Alexa Fluor 568-dextran fluorescence (left) and Alexa Fluor 647-ASO fluorescence (right) within endolysosomal compartments positive for Alexa Fluor 647-ASO after a 2.5-hours pulse and negative for Alexa Fluor 488-ASO after an 18-hours chase in wild-type cells, before and after treatment with 0.5 mM LLOMe for 30 min (# endosomes pre-LLOMe = 10,197; post-LLOMe = 15,539).

**(B)** Quantification of replicates from Fig. 3 B, D, shown as frequency plots of Alexa Fluor 568-dextran fluorescence (left) and Alexa Fluor 647-ASO fluorescence (right) within lysosomal compartments positive for both Alexa Fluor 647-ASO after a 2.5-hours pulse and Alexa Fluor 488-ASO after an 18-hours chase in wild-type cells, before and after treatment with 0.5 mM LLOMe for 30 min (# endosomes pre-LLOMe = 16,092; post-LLOMe = 14,404).

**(C)** Frequency plots of the Alexa Fluor 647-ASO fluorescence to Alexa Fluor 568-dextran ratio within endolysosomal compartments positive for Alexa Fluor 647-ASO after a 2.5-hours pulse and negative for Alexa Fluor 488-ASO after an 18-hours chase in wild-type cells, before and after treatment with 0.5 mM LLOMe for 30 min (# endosomes pre-LLOMe = 10,197; post-LLOMe = 15,539). Median values from the distributions, plotted as bar graph. Each dot represents an independent biological replicate. Data are from three biological replicates. Statistical significance: $p<0.05$ (*).

**(D)** Frequency plots of the Alexa Fluor 647-ASO fluorescence to Alexa Fluor 568-dextran ratio within lysosomal compartments positive for both Alexa Fluor 647-ASO after 2.5-hours pulse and Alexa Fluor 488-ASO after an 18-hours chase in wild-type cells, before and after treatment with 0.5 mM LLOMe for 30 min (# endosomes pre-LLOMe = 16,092; post-LLOMe = 14,404). Median values from the distributions, plotted as bar graph. Each dot represents an independent biological replicate. Data are from three biological replicates. Statistical significance: $p<0.01$ (**).

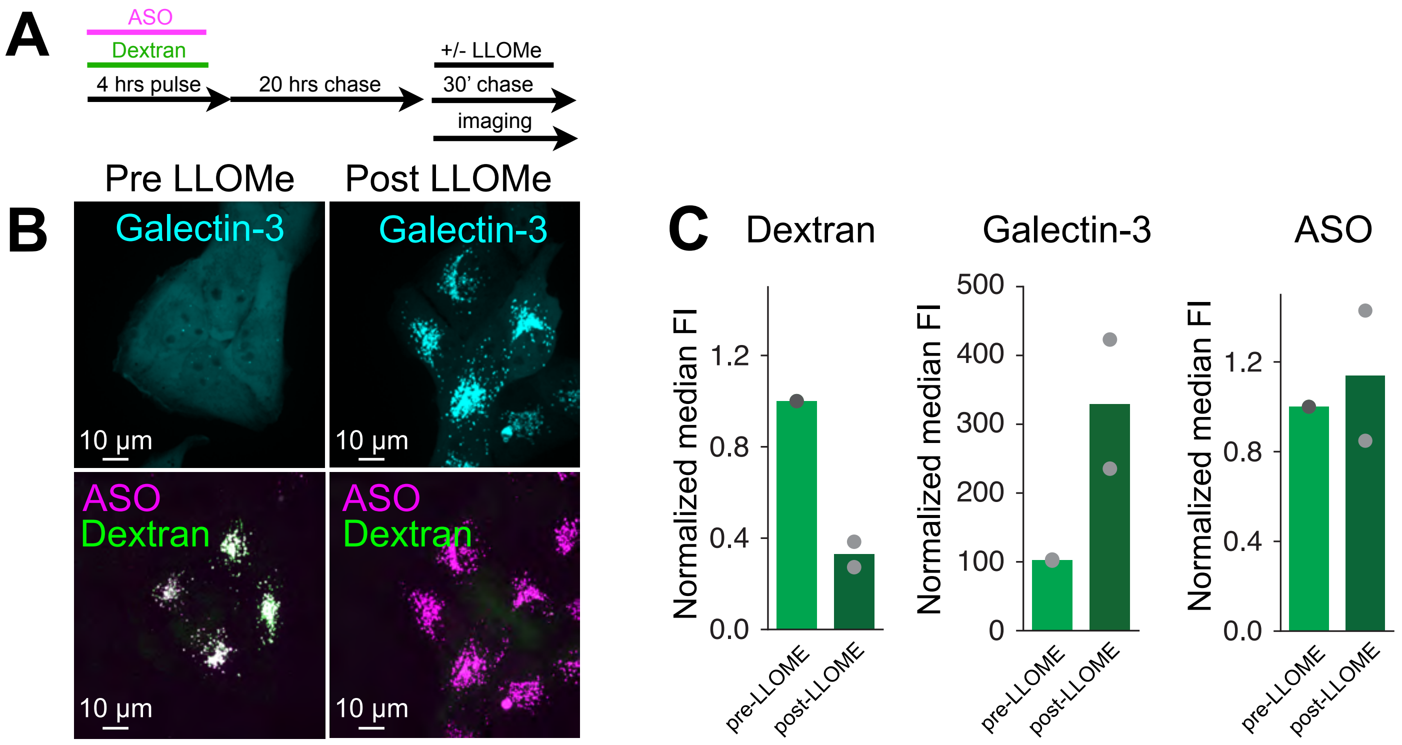


### Supplementary Figure S5

**Supplementary Figure S5. Related to Figure 3.**

**(A)** Experimental protocol. U2OS cells stably expressing galectin-3 eGFP (cyan) were preloaded with 1 μM Alexa Fluor 488-ASO (magenta) and 8 μM Alexa Fluor 568-dextran (green) for 4-hours and chased for 20-hours to terminal lysosomes with or without 0.5 mM LLOMe for 30 min before imaging.

**(B)** Representative maximum-projection spinning-disc confocal images of wild-type U2OS cells stably expressing galectin-3-eGFP (cyan), with terminal lysosomes labelled by pulse-chase with Alexa Fluor 568-dextran (green) and Alexa Fluor 647-ASO (magenta), untreated or treated with 0.5 mM LLOMe for 30 min. Scale bar, 10 μm.

**(C)** Quantification of **(B)**, shown as normalized median dextran, galectin-3 and ASO fluorescence intensities in ASO-positive compartments in untreated cells (light green) and cells treated with 0.5 mM LLOMe for 30 min (dark green). Fluorescence intensity in the LLOMe condition was normalized to the corresponding control in each independent experiment. Data are from two independent experiments.

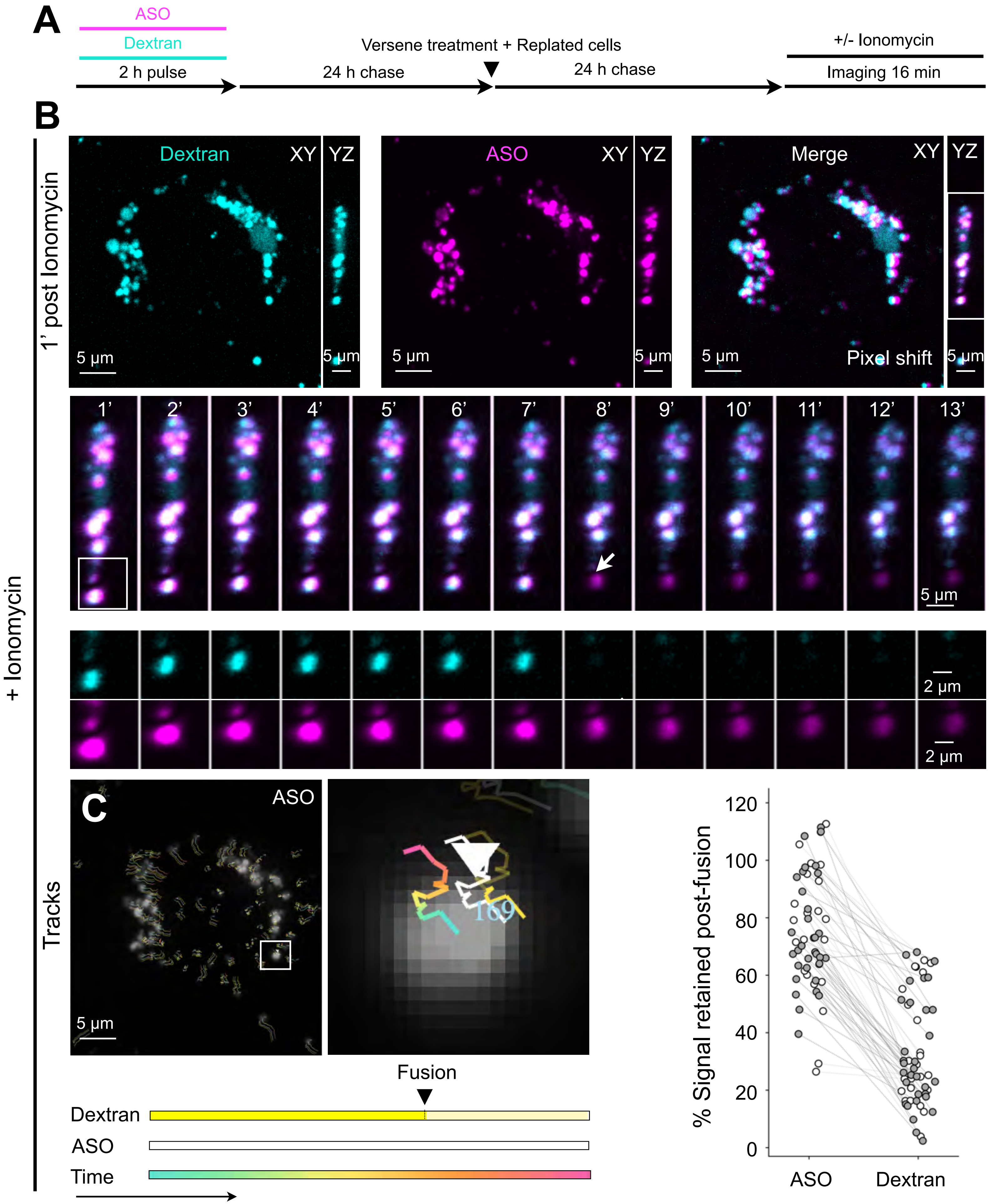


### Supplementary Figure S6

**Supplementary Figure S6. Related to Figure 3.**

**(A)** Experimental protocol. U2OS cells were pulsed for 2 h with Alexa Fluor 647–ASO and Alexa Fluor 568–labeled 10-kDa dextran, chased for 24 h, detached with Versene and replated on fresh coverslips. After a further 24 h, cells were treated with ionomycin and imaged by time-lapse spinning-disk confocal microscopy.

**(B)** Representative images of wild-type U2OS cells containing dextran (cyan) and ASO (magenta) in terminal lysosomes, acquired 1–13 min after ionomycin addition. In the merged image, the channels are offset by 5 pixels to aid visualization of colocalization. The white box marks a lysosome that underwent fusion in the time series of the YZ projection; the white arrow indicates the first frame showing dextran loss. Enlarged views are shown below.

**(C)** Trajectories from the cell in **(B)**, projected across z and time. Each trajectory is displayed in three adjacent representations: color-coded by time; in white, marking the ASO-mask centroid obtained using 3D-CME; and in yellow, with change in brightness to indicate the abrupt loss of dextran fluorescence intensity associated with the fusion event.

**(D)** The fusion event is shown at higher magnification. Scale bars, 2 or 5 μm, as indicated.

**(E)** Comparison of lysosomal dextran fluorescence immediately before and after fusion. Each point represents one lysosome; gray (n = 24) and white (n = 32) points show measurements from different cells in two independent experiments. The percentage of fluorescence retained after fusion was calculated as 100 × (post-fusion/pre-fusion signal), using the frames immediately before and after fusion and the same mask for ASO and dextran.

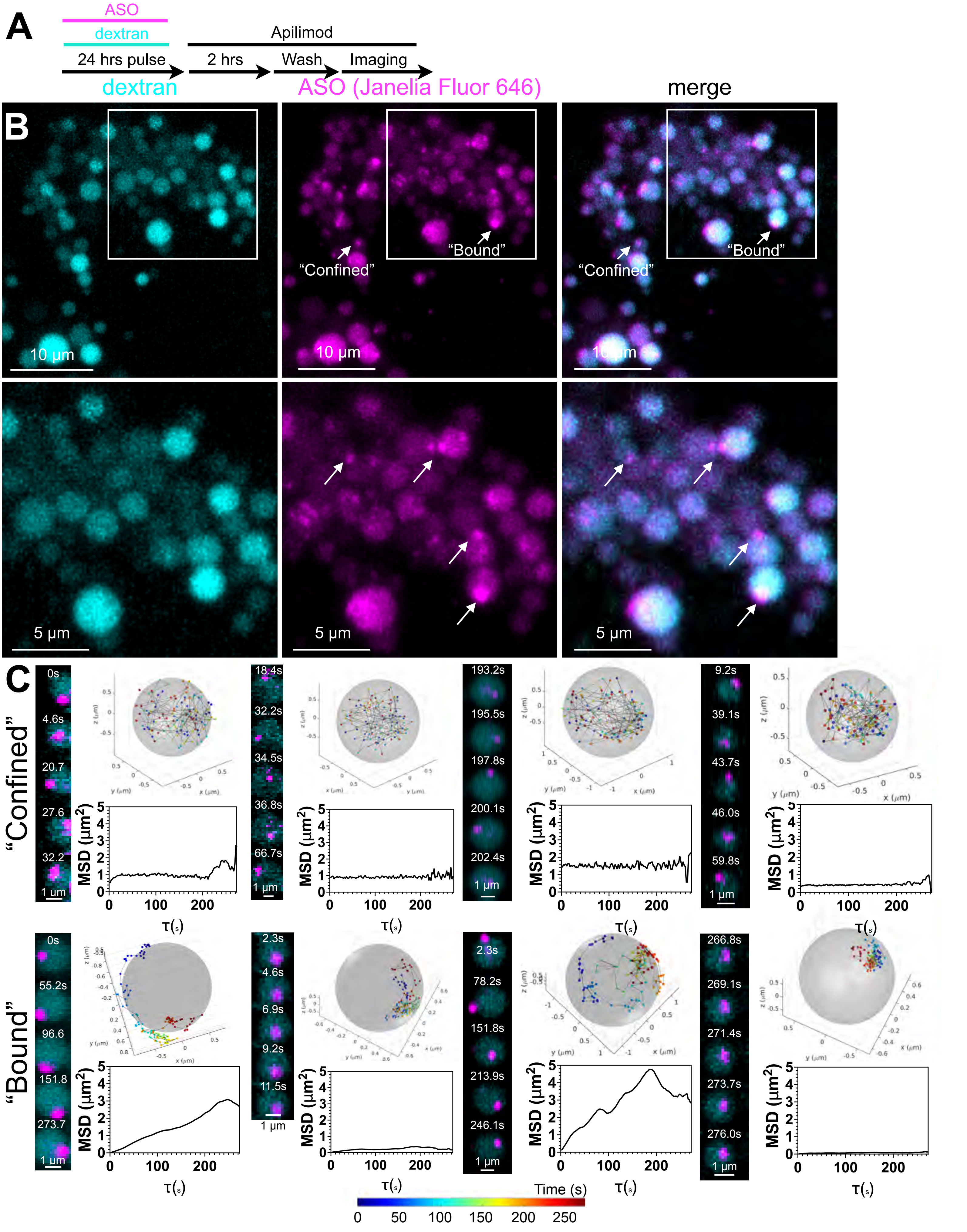


### Supplementary Figure S7

**Supplementary Figure S7. Related to Figure 5.**

**(A)** Experimental protocol. Wild-type U2OS cells were preloaded for 24-hours with 8 μM Alexa Fluor 568-dextran and 1 μM Janelia Fluor 646-ASO, chased for 2-hours in the presence of 5 μM apilimod, washed, and imaged.

**(B)** Representative lattice light-sheet (Zeiss) images of wild-type U2OS cells labeled as in **(A)**. A single middle plane is shown. Alexa Fluor 568-dextran is shown in cyan and Janelia Fluor 646-ASO in magenta. Arrows indicate ASO puncta undergoing confined and bound motion. Scale bar, 10 μm. Lower panel: Arrows indicate discrete punctate of intraluminal ASO in endolysosomes, whereas the rest of the intraluminal ASO and dextran fills the same compartment uniformly. Scale bar, 10 μm.

**(C)** Representative three-dimensional trajectories of individual ASO foci showing confined (top panel) or bound (bottom) motion in additional apilimod-expanded endolysosomes. Endolysosomes are represented as semitransparent spheres, with ASO positions color-coded from blue to red to indicate progression through time. Insets show maximum-intensity projections at successive time points, with ASO in magenta and the endolysosomal signal in cyan. Scale bar, 1 μm. Corresponding mean-squared-displacement plots are shown for each trajectory.

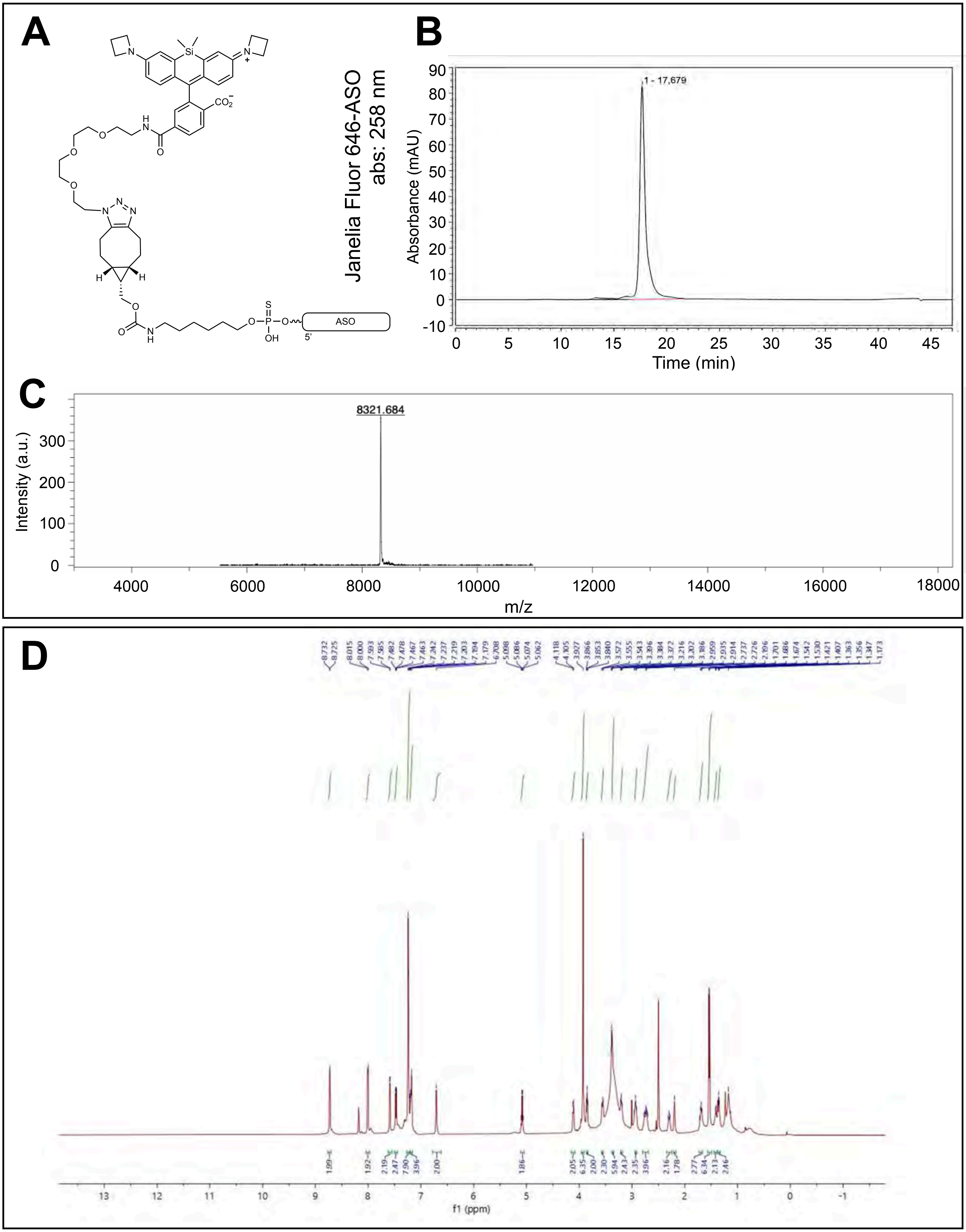


### Supplementary Figure S8

**Supplementary Figure S8: Related to Figure 4 and 5**

**(A)** Structure of Janelia Fluor 646 – ASO. *Oligonucleotide synthesis:* ASO synthesis was performed on an Applied Biosystems (ABI) 394 DNA/RNA Synthesizer at a 3 μmol scale utilizing the standard phosphoramidite approach. The solid support used was the Glenn UnySupport™ CPG 1000 45 μmol/g. The manufacturer's standard protocol was carried out with the following alterations: the coupling time was extended to 159 s, the sulfurization time was increased to 119 s, and additional washing steps were integrated throughout the synthesis cycle. The efficiencies of each coupling step were determined to be an average of 97.5% via monitoring the conductivity of the released dimethoxytrityl cation. The primary amine function was introduced to the 5'-end during the synthesis as a phosphoramidite by using 12 equivalents of 5'-MMT-Amino Modifier C6 phosphoramidite dissolved in 0.4 mL anhydrous acetonitrile and 0.6 mL of the activator (0.25 M 5-ethylthio-1H-tetrazole (ETT) in anhydrous acetonitrile) via manual coupling for 15 min. Following synthesis, cleavage from solid support and global deprotection were done using 28% aqueous ammonia (8 h, 65 °C). The resulting mixture was filtered, and the beads were washed with acetonitrile-water 1:1 and evaporated under nitrogen flow. The resulting ASO was purified with MMT on. Purification was conducted on a Vanquish UHPLC system equipped with a C18 column (250 x 10 mm from Phenomenex) maintained at 60°C. Elution system consisted of solvent A: 0.05 M triethylammonium acetate buffer in Milli-Q water, pH 7.4 and solvent B: acetonitrile. The flow rate of the system was 4 mL/min. The elution gradient started with a linear gradient from 5-60% of solvent B over 27.5 min. Subsequently, detritylation was performed using 150 μL 90% aqueous solution of acetic acid (30 min, 30 °C, 700 rpm shaking). The product was desalted by addition of sodium acetate (3 M, 15 μL) and sodium perchlorate (5 M, 15 μL), followed by precipitation with absolute ethanol at -20 °C for 1 h. The solution was centrifuged at (13200 rpm, 5 min, 4 °C), the supernatant was discarded, and the pellet was further washed with ice-cold absolute ethanol (3 x 1 mL) and dried under nitrogen flow to provide the 5'-amino-C6 modified ASO, and concentration was determined by UV-Vis measurements at 260 nm. *BCN-functionalization of ASO***.** *Endo*-BCN N-hydroxy succinimide carbonate (30 equiv., BroadPharm) was dissolved in acetonitrile (300 μL). 5'-Amino-C6 modified ASO (0.3 μmol) was dissolved in Milli-Q water (300 μL) and 0.5 M bicarbonate buffer (pH 8.5, 300 μL) and allowed to react (2 h, rt, 700 rpm shaking). The 5'-BCN-ASO was desalted using NAP-10 column (Cytiva) to remove the inorganic salts and purified by UHPLC as described above. *Janelia Fluor 646 azide coupling with 5'-BCN-ASO:* The 5'-BCN-ASO (50 nmol) was dissolved in Mili-Q water (500 μL) and Janelia 646 (1.2 equiv.) functionalized with an azide was dissolved in *N,N*-dimethylformamide (50 μL) and reacted in a Biotage microwave instrument (2 h, 65°C). The Janelia Fluor 646-ASO was purified by UHPLC as described above and dried under nitrogen flow. Desalting was done with NAP column following the manufactures procedure. The Janelia Fluor 646-ASO purity was confirmed on HPLC, and its composition confirmed on MALDI-TOF mass spectrum calcd. 8318.8; m/z found 8321.684.

**(B)** HPLC chromatogram (258 nm) of Janelia Fluor 646-ASO, obtained on Vanquish UHPLC equipped with a C18 column (250 x 10 mm from Phenomenex) at 60°C.

**(C)** MALDI-TOF mass spectrometry obtained on a Bruker Mircroflex®; calculated m/z, 8318.8; experimentally determined, 8321.684.

**(D)** NMR spectrum of tantalosin synthesized and purified for use in this study. Tantalosin was synthesized following the published route (52). Starting materials were obtained from commercial sources. Solvents were obtained from Sigma-Aldrich, Oakwood, and Ambeed and used directly without further purification. Analytical thin layer chromatography was performed on 0.25 mm silica gel 60-F254. Visualization was carried out with UV light and Bromocresol Green Stain. $^1$H NMR spectra were recorded on Bruker DRX-600 instrument (600 MHz). Chemical shifts (δ) are reported in parts per million (ppm) and referenced to the literature values of the center line of a multiplet at 2.50 ppm of DMSO-$d^6$ or a singlet at 3.33 ppm of water. The following abbreviations (or combinations thereof) were used to describe multiplicities: s = singlet, d = doublet, t = triplet, q = quartet, p = pentet, m = multiplet, br = broad. Coupling constants, *J*, were reported in Hertz unit (Hz). [1] High-resolution mass spectra (HRMS) were recorded on a Bruker Mass spectrometer using ESI-TOF (electrospray ionization-time of flight). The identity of product was confirmed by LC-MS comparison with a standard obtained from Yao-Wen Wu lab.

$^1$H NMR (600 MHz, DMSO-$d^6$) δ 8.73 (d, *J* = 4.2 Hz, 2H), 8.01 (d, *J* = 9.0 Hz, 2H), 7.58 (d, *J* = 4.8 Hz, 2H), 7.47 (dd, *J* = 9.0, 2.4 Hz, 2H), 7.22-7.24 (m, 8H), 7.18-7.20 (m, 4H), 6.71 (s, 2H), 5.08 (q, *J* = 7.2 Hz, 2H), 4.11 (d, *J* = 7.8 Hz, 2H), 3.93 (s, 6H), 3.83 (t, *J* = 7.8 Hz, 2H), 3.54-3.57 (m, 2H), 3.37-3.40 (m, 6H), 3.19-3.22 (m, 2H), 2.91-3.00 (m, 2H), 2.70-2.77 (m, 4H), 2.30 (t, *J* = 10.8 Hz, 2H), 2.20 (s, 2H), 1.67-1.72 (m, 2H), 1.54 (d, *J* = 7.2 Hz, 6H), 1.41-1.42 (m, 2H), 1.35-1.36 (m, 2H). HRMS (ESI-TOF) *m/z* Calcd for $C_{66}H_{70}N_8O_{10}^{2+}$ $[M+2H]^{2+}$ 567.2602, found 567.2604.

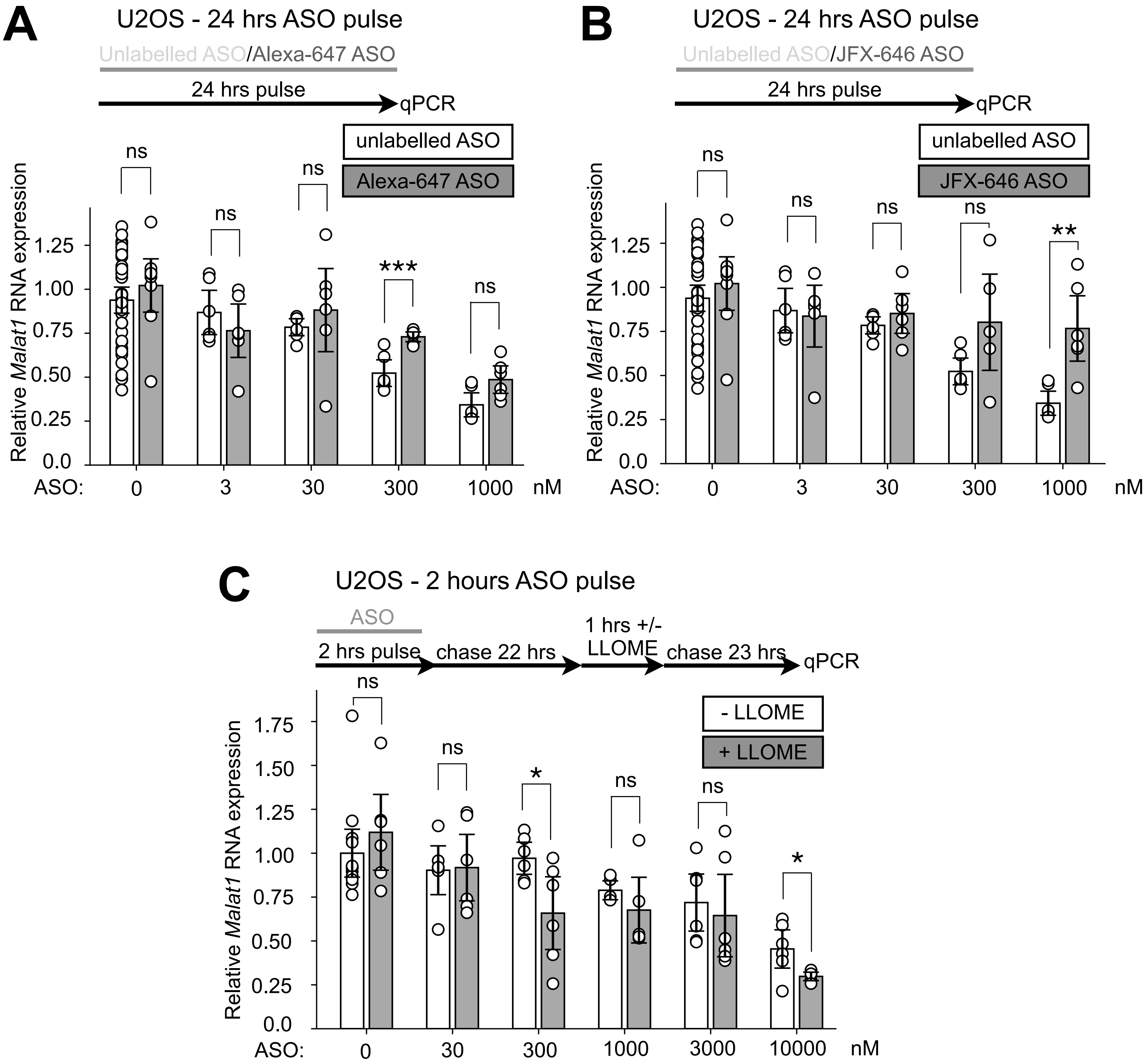


**Supplementary Figure S9**

**Supplementary Figure 9. Related to Figure 4.**

**(A)** Effect of 24-hour treatment with increasing amounts of Malat1 ASO on *Malat1* RNA levels in U2OS cells with unconjugated ASO or Alexa-647-conjugated ASO. RNA abundance was measured by quantitative PCR, normalized to cyclophilin, and plotted as fold change relative to the corresponding untreated control. Each dot represents a replicate; black lines indicate the mean and error bars indicate SEM. Data are from at least six biological replicates from two independent experiments.

**(B)** Effect of 24-hour treatment with increasing amounts of Malat1 ASO on *Malat1* RNA levels in U2OS cells with unconjugated ASO or Janelia-646-conjugated ASO. RNA abundance was measured by quantitative PCR, normalized to cyclophilin, and plotted as fold change relative to the corresponding untreated control. Each dot represents a replicate; black lines indicate the mean and error bars indicate SEM. Data are from at least six biological replicates from two independent experiments.

**(C)** Effect of 2-hour treatment and 22 hours chase with increasing amounts of Malat1 ASO on *Malat1* RNA levels in U2OS cells with or without 0.5 mM LLOME incubation (when ASO reached terminal lysosomes) for 1 hours followed by additional 23 hours chase. RNA abundance was measured by quantitative PCR, normalized to cyclophilin, and plotted as fold change relative to the corresponding untreated control. Each dot represents a replicate; black lines indicate the mean and error bars indicate SEM. Data are from at least six biological replicates from two independent experiments.

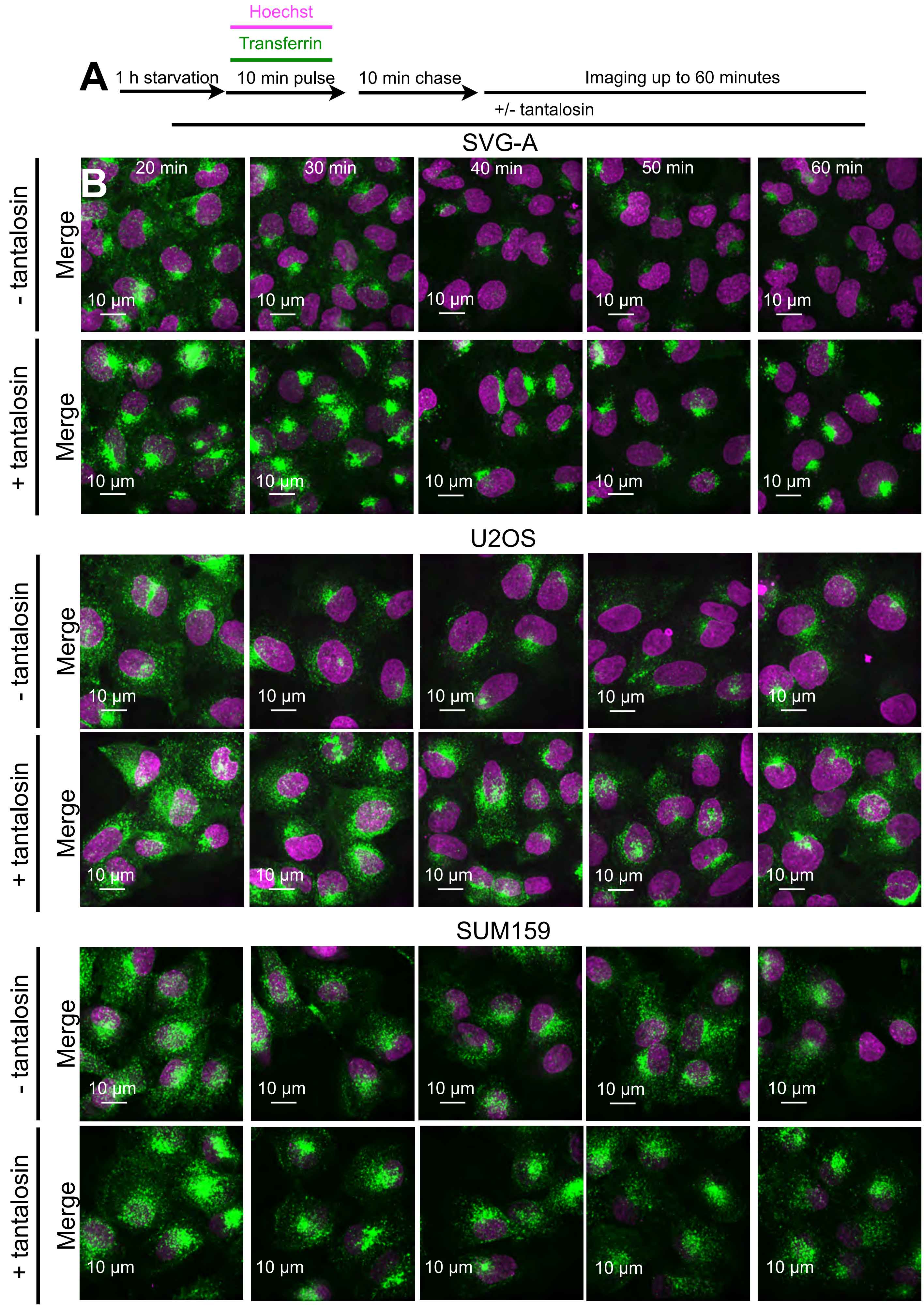


### Supplementary Figure S10

**Supplementary Figure S10. Related to Figure 5.**

**(A)** Experimental protocol. After 1 hour starvation in medium without serum cells were pulsed for 10 min with Alexa Fluor 647–transferrin, washed, chased for 10 min, and then imaged live for up to 60 min in the absence or presence of 5 µM tantalosin.

**(B)** Representative maximum-intensity projections from spinning-disk confocal image stacks of wild-type SVG-A cells (top), U2OS (middle) and SUM159 (bottom) in the absence or presence of 5 µM tantalosin. Alexa Fluor 647–transferrin is shown in green and Hoechst-labeled nuclei in magenta. Scale bar, 10 µm.

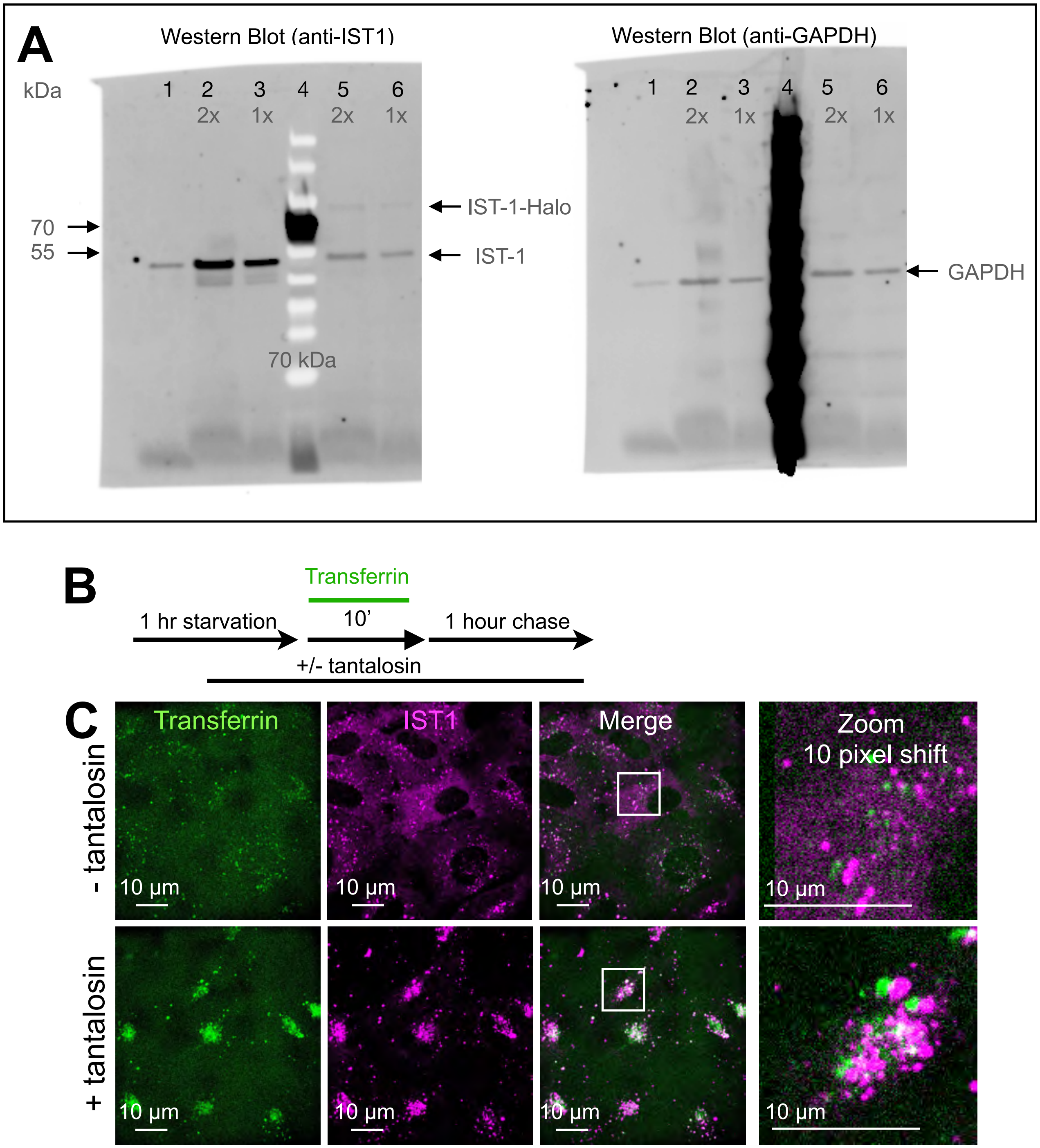


### Supplementary Figure S11

**Supplementary Figure S11. Related to Figure 5.**

**(A)** Immunoblots for IST1 and GAPDH in parental SVG-A cells (lane 1), a negative clone (lanes 2 and 3, 1× and 2× loading, respectively), and an SVG-A clone with one IST1 allele edited to express IST1–Halo (lanes 5 and 6, 1× and 2× loading, respectively). Arrows indicate bands at the expected molecular mass of IST1–Halo.

**(B)** Experimental protocol. After 1 h of serum starvation, IST1–Halo SVG-A cells were pulsed for 10 min with Alexa Fluor 647–transferrin, washed, chased for 10 min and imaged live for up to 60 min in the absence or presence of 5 μM tantalosin.

**(C)** Representative maximum-intensity projections of spinning-disk confocal image stacks from IST1–Halo SVG-A cells imaged with or without 5 μM tantalosin. Alexa Fluor 647–transferrin is shown in green and IST1–Halo labeled with Janelia Fluor 549 HaloTag ligand in magenta. Zoom-in image is shifted by 10 pixels for visualization purposes. Images show tantalosin-induced accumulation of IST1–Halo on transferrin-positive early and recycling endosomes during early endocytic trafficking. Scale bar, 10 μm.

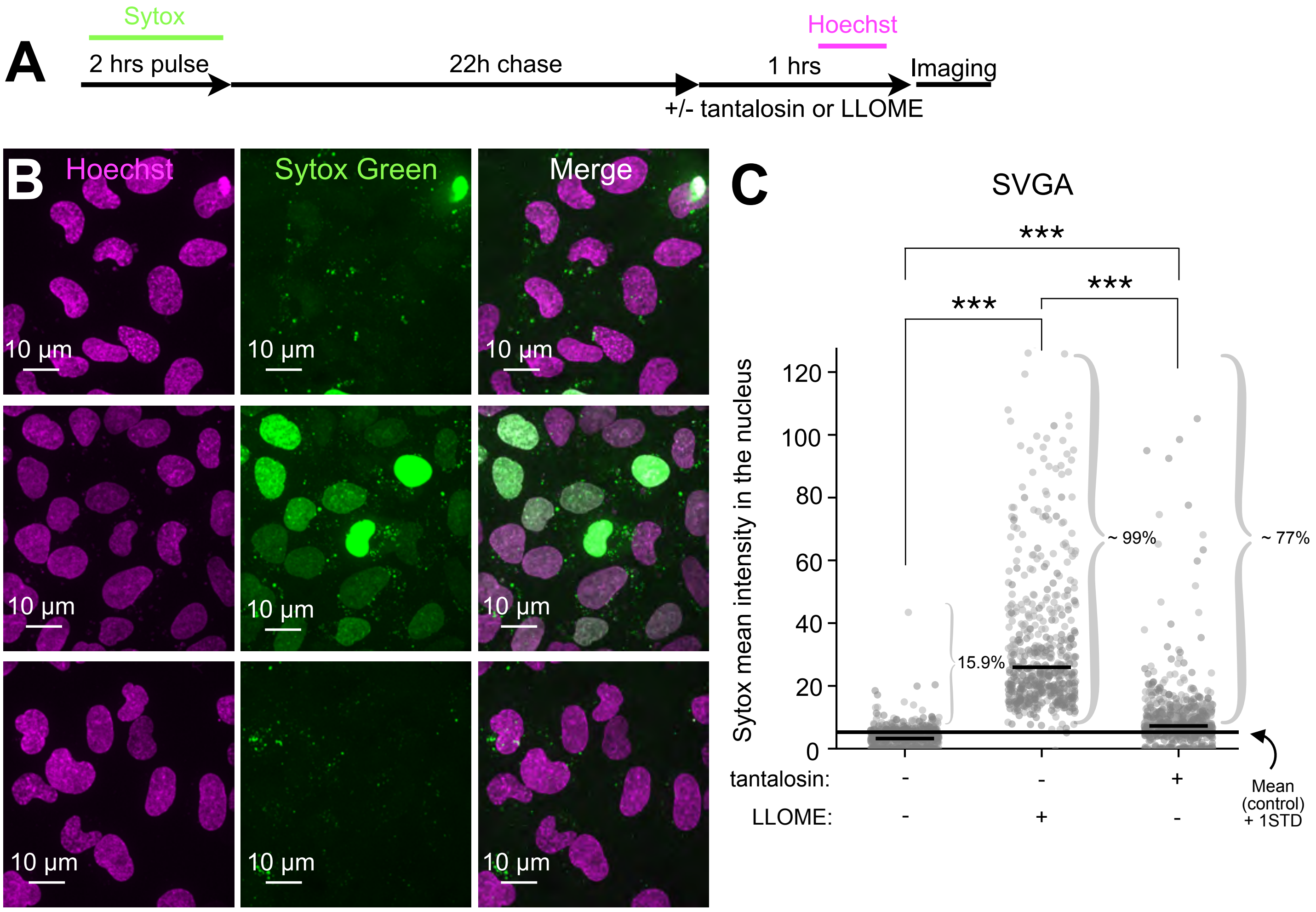


### Supplementary Figure S12

**Supplementary Figure S12. Related to Figure 5.**

**(A)** Experimental protocol. Cells were pulsed for 2 h with 5 µM SYTOX Green, washed and chased for 22 h to allow the dye to accumulate in terminal lysosomes. Cells were then left untreated or treated for 1 h with 5 µM tantalosin or 0.5 mM LLOMe and imaged live by spinning-disk confocal microscopy.

**(B)** Representative maximum-intensity projections of wild-type SVG-A cells treated as in **(A)**. SYTOX Green is shown in green and Hoechst-labeled nuclei in magenta. Increased nuclear fluorescence reports escape of internalized SYTOX Green from lysosomes into the cytosol.

**(C)** Mean SYTOX Green fluorescence within three-dimensionally segmented nuclei. Numbers of nuclei analyzed were 894 untreated, 873 tantalosin-treated and 547 LLOMe-treated, from four, four and two independent experiments, respectively. Each point represents one nucleus; black horizontal lines indicate medians. The percentage of SYTOX-positive cells and its SD are shown below each graph. Cells were scored as positive when their mean nuclear fluorescence exceeded the matched control mean by more than one SD. Statistical significance: $p < 0.001$ (***).

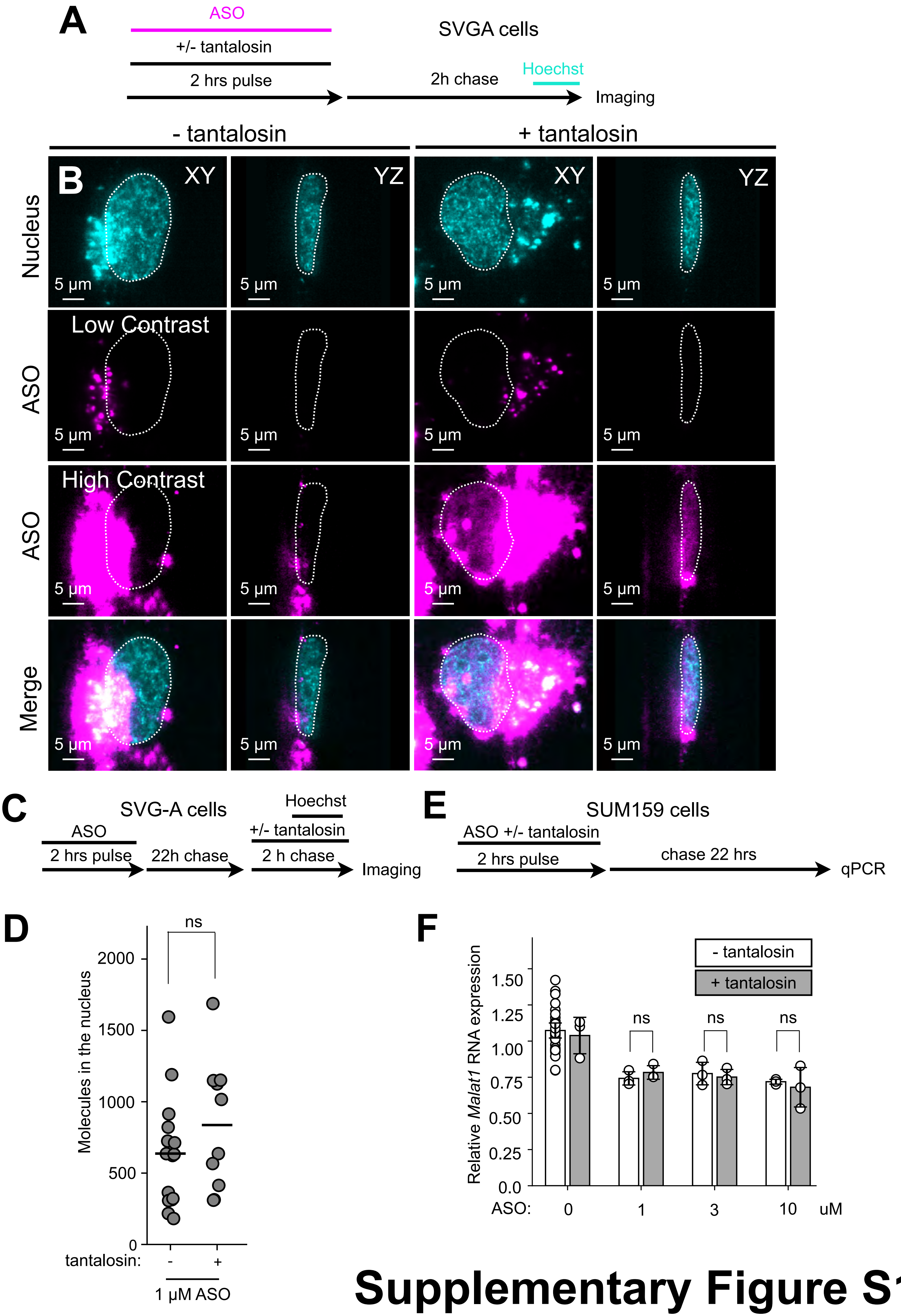


### Supplementary Figure S13

**Supplementary Figure S13. Related to Figure 6.**

**(A)** Experimental protocol. SVGA cells were incubated for 2 h with 1 μM ASO in the absence or presence of 5 μM tantalosin and then chased for 2 h. JF549–Hoechst was added during the final 30 min of the chase, after which the cells were imaged live.

**(B)** Representative single optical sections acquired by MOSAIC lattice light-sheet microscopy from wild-type SVG-A cells treated as in (**A**). Alexa Fluor 647–ASO is shown in green and JF549–Hoechst-labeled nuclei in magenta. ASO fluorescence is displayed at increased contrast to reveal the nuclear signal; fluorescence from endosomal ASO is therefore saturated. White dotted lines outline the nuclei. Corresponding XY and YZ views are shown. Scale bar, 5 μm.

**(C)** Experimental protocol. SVGA cells were incubated for 2 h with 1 μM ASO, chased for 22 h (to target them to lysosomes) and then incubated for 2 hrs in the absence or presence of 5 μM tantalosin. JF549–Hoechst was added during the final 30 min of the chase, after which the cells were imaged live.

**(D)** Number of ASO molecules per nucleus of SVGA cells after a 2-hour pulse with 1 μM ASO followed by a 22 hour chase, treated or not treated with 5 μM tantalosin when ASO is in the terminal lysosome. Each point represents one nucleus; the horizontal line indicates mean. Numbers of nuclei analyzed were 16 control and 10 tantalosin-treated after 22 hours chase. Data shown are representative of two independent experiments.

**(E)** Experimental protocol. SUM159 cells were incubated for 2 h with different concentrations of ASO in the absence or presence of 5 μM tantalosin and then chased for 22 h followed by qPCR quantification.

**(F)** Effect of tantalosin on the functional activity of ASOs in SUM159 cells. Cells were incubated for 2 h with increasing concentrations of *Malat1* ASO, in the absence or presence of 5 μM tantalosin, and then chased for 22 h. *Malat1* RNA abundance was measured by quantitative PCR, normalized to cyclophilin, and expressed relative to the corresponding untreated control. Each point represents one biological replicate; black lines indicate means and error bars indicate SEM. Data are from at least six biological replicates collected in two independent experiments.

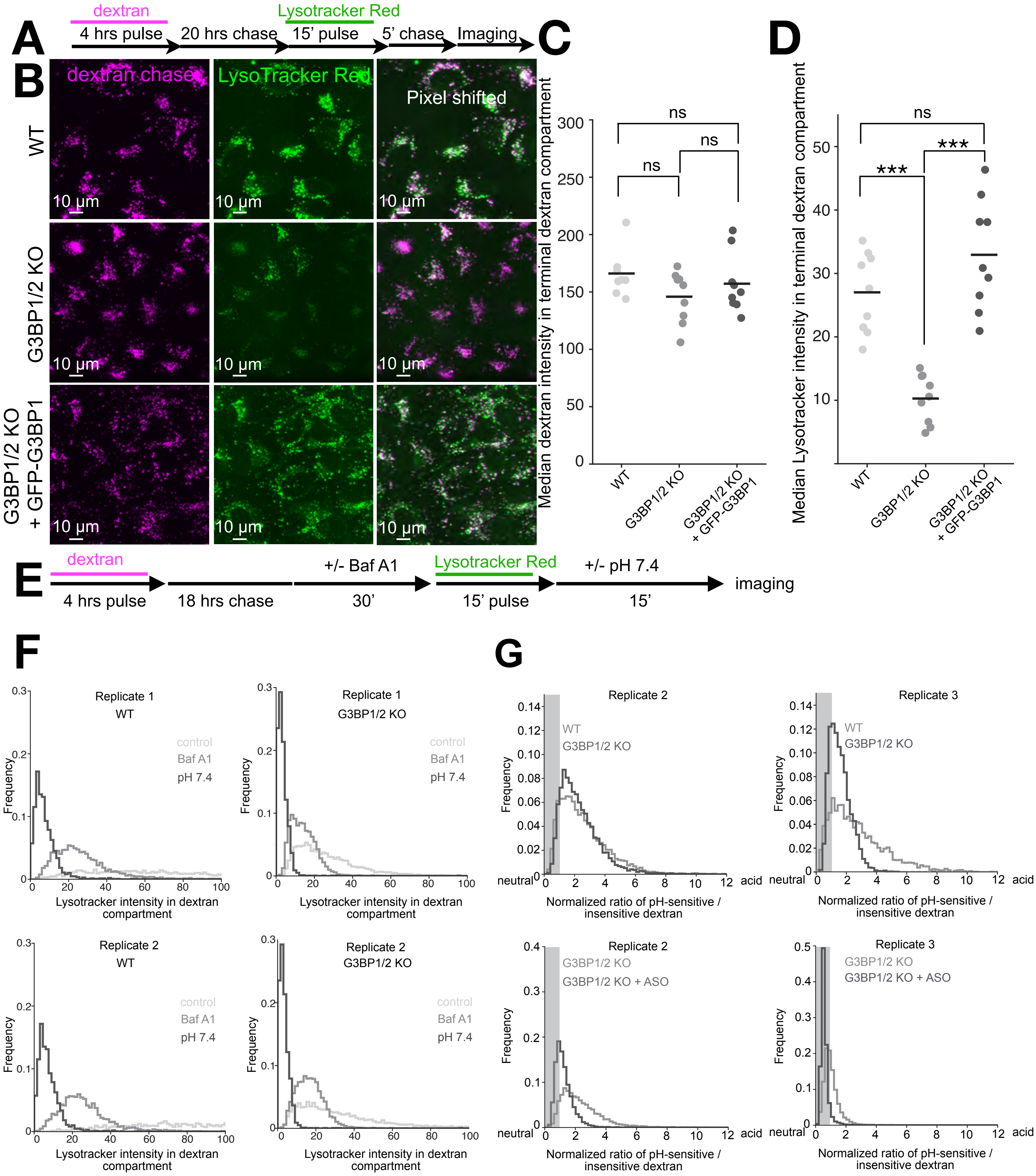


### Supplementary Figure S14

**Supplementary Figure S14. Related to Figure 7.**

**(A)** Experimental Protocol. U2OS wild-type, U2OS G3BP1/2 knockout and U2OS G3BP1/2 knockout cells stably expressing eGFP-G3BP1 cells were treated with 4 µM dextran for 4 hours, then chased for 20 hours to the terminal lysosome. Next, Lysotracker Red was added for 15 minutes, chased for 5 minutes and then cells subjected to live cell imaging.

**(B)** Representative maximum-projection lattice light-sheet (Zeiss) images of wild-type U2OS cells, G3BP1/2 knockout cells and G3BP1/2 knockout U2OS cells stably expressing eGFP-G3BP1 cells pulsed with 8 µM Alexa Fluor 647-dextran (magenta) for 4-hours and stained with LysoTracker™ Red (green). Scale bar, 10 µm.

**(C)** Quantification of Alexa Fluor 647-dextran fluorescence from **(A)** in wild-type U2OS cells (gray), G3BP1/2 knockout cells (dark gray) and G3BP1/2 knockout U2OS cells stably expressing eGFP-G3BP1 cells (black). Plots show median dextran fluorescence intensity from three biological replicates, with three fields of view per replicate, for wild-type cells (# endosomes = 25,183), G3BP1/2 knockout cells (# endosomes = 33,803) and G3BP1/2 knockout cells stably expressing eGFP-G3BP1 (# endosomes = 32,465). Statistical significance: $p>0.05$ (ns); $p<0.05$ (*); $p<0.01$ (**); $p<0.001$ (***).

**(D)** Quantification of LysoTracker™ Red fluorescence within dextran-positive compartments from **(B)** in wild-type cells (gray), G3BP1/2 knockout cells (dark gray) and G3BP1/2 knockout U2OS cells stably expressing eGFP-G3BP1 cells (black). Plots show median LysoTracker™ Red fluorescence intensity from three biological replicates, with three fields of view per replicate, for wild-type cells (# endosomes = 25,183), G3BP1/2 knockout cells (# endosomes = 33,803) and G3BP1/2 knockout cells stably expressing eGFP-G3BP1 (# endosomes = 32,465). Statistical significance: $p>0.05$ (ns); $p<0.001$ (***).

**(E)** Experimental protocol. Wild-type U2OS cells were treated with 4 µM dextran for 4 hours and chased for 18 hours to the terminal lysosome. Next, cells were treated for 30 minutes with either 100 nM bafilomycin A1 or with pH 7.4 universal buffer containing 10 µM nigericin plus 10 µM monensin.

**(F)** Frequency distributions of LysoTracker™ Red fluorescence intensity within dextran-positive compartments of wild-type U2OS cells and G3BP1/2 knockout cells treated with 100 nM bafilomycin for 30 min or clamped at pH 7.4, at the indicated time points in the experimental schematic shown.

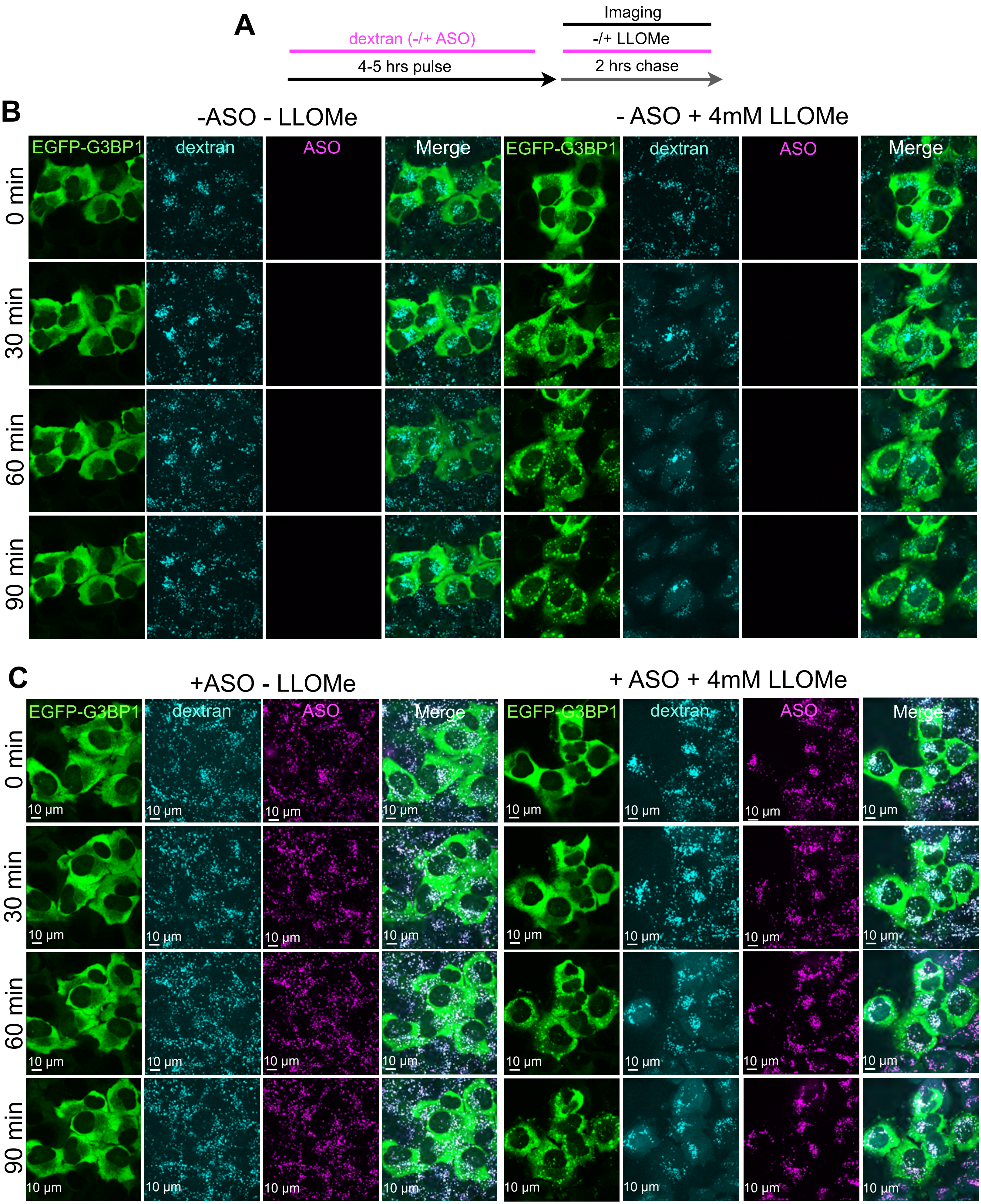


### Supplementary Figure S15

**Supplementary Figure S15. Related to Figure 7.**

**(A)** Experimental protocol. G3BP1/2-knockout U2OS cells stably expressing eGFP-G3BP1 were pulsed for 4 h with 8 µM Alexa Fluor 568–dextran to label late endosomes, endolysosomes and lysosomes, with or without 1 µM Alexa Fluor 647–ASO and imaged during a 2-h chase in the absence or presence of 4 mM LLOMe.

**(B)** Representative maximum-intensity projections from time-lapse spinning-disk confocal image stacks of G3BP1/2-knockout U2OS cells stably expressing eGFP-G3BP1 (green). Cells were pulsed for 4 h with 8 µM Alexa Fluor 647–dextran (cyan) and imaged during a 2-h chase with or without 4 mM LLOMe. Scale bar, 10 µm.

**(C)** Representative maximum-intensity projections from time-lapse spinning-disk confocal image stacks of the same cell line, showing eGFP–G3BP1 (green), Cy3–ASO (magenta) and Alexa Fluor 647–dextran (cyan). Cells were pulsed for 2 h with 1 µM Cy3–ASO and 8 µM Alexa Fluor 647–dextran, chased for 2 h, and then left untreated or treated with 4 mM LLOMe. Scale bar, 10 µm.

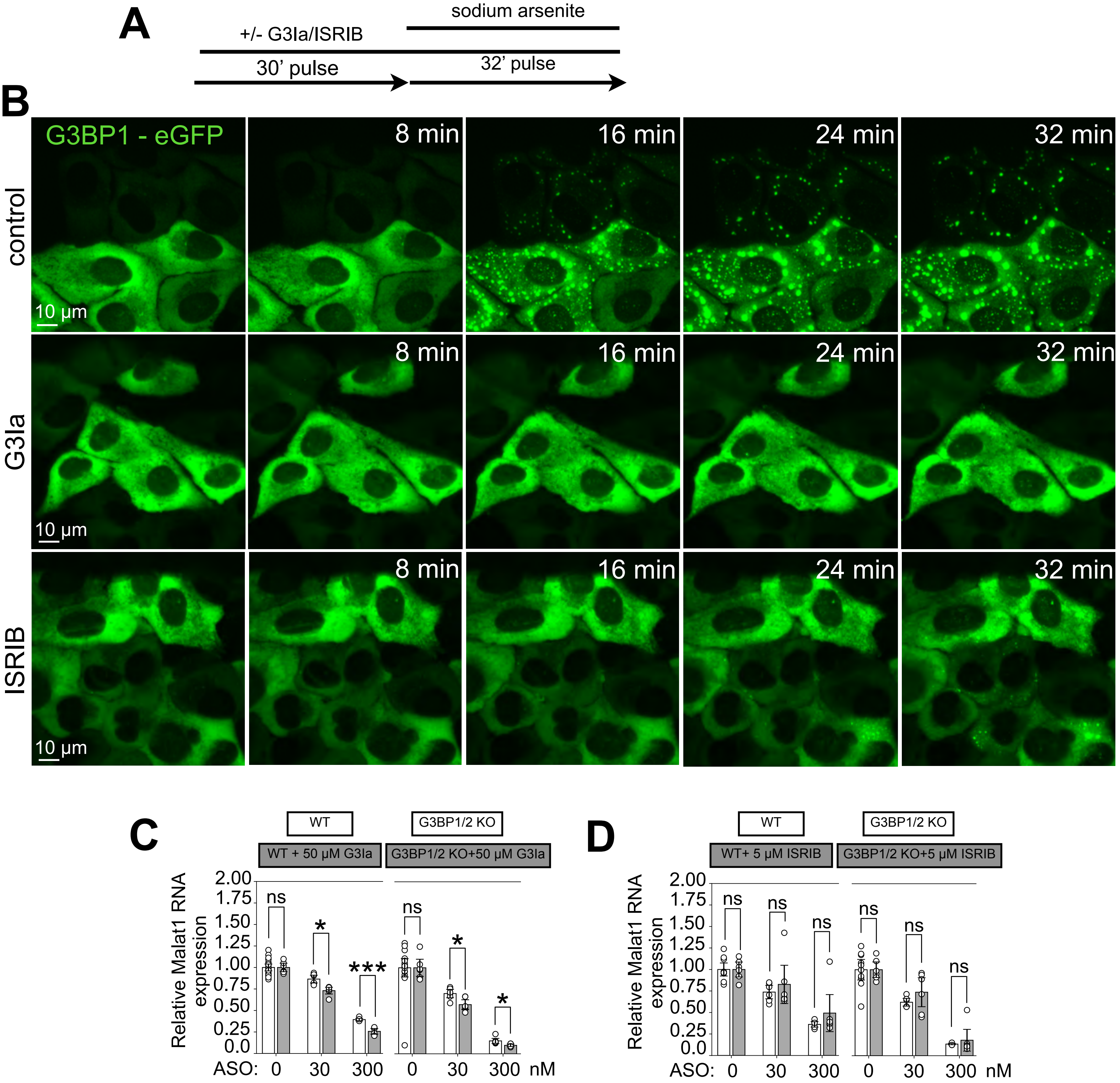


### Supplementary Figure S16

**Supplementary Figure S16. Related to Figure 7.**

**(A)** Experimental protocol. G3BP1/2-knockout U2OS cells stably expressing eGFP–G3BP1 were preincubated for 30 min with or without 1 μM ISRIB or 50 μM G3Ia, then treated with 0.25 mM sodium arsenite.

**(B)** Representative maximum-intensity projections from lattice light-sheet microscopy (Zeiss) time series of cells treated with sodium arsenite alone (control) or in the presence of ISRIB or G3Ia, as in **(A)**. Scale bar, 10 μm.

**(C)** Relative *Malat1* RNA levels following treatment with the indicated ASO concentrations alone in wild-type or G3BP1/2-knockout U2OS cells (gray), or together with 50 μM G3Ia (dark gray).

**(D)** Relative *Malat1* RNA levels following treatment with the indicated ASO concentrations alone in wild-type or G3BP1/2-knockout U2OS cells (gray), or together with 5 μM ISRIB (gray).

For **(C,D)**, RNA abundance was measured by quantitative PCR, normalized to cyclophilin and plotted as fold change relative to the corresponding untreated control. Black lines indicate means; error bars indicate SEM. Data are from three independent experiments. Statistical significance: $p > 0.05$ (ns); $p < 0.05$ (*); $p < 0.001$(***).

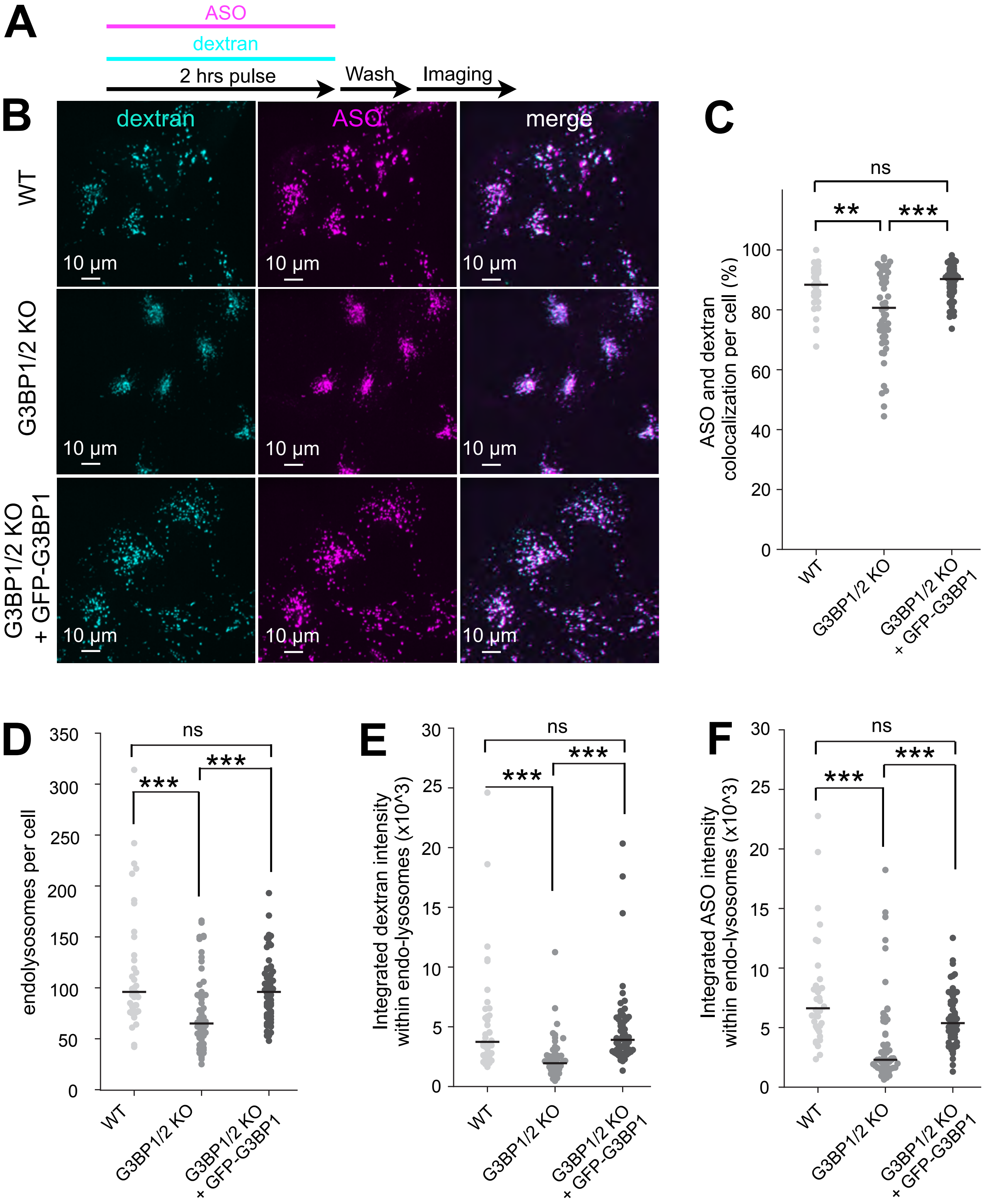


### Supplementary Figure S17

**Supplementary Figure S17. Related to Figure 8.**

**(A)** Experimental protocol. Cells were pulsed for 2 h with 8 µM Alexa Fluor 568–labeled 10-kDa dextran and 1 µM Alexa Fluor 647–ASO, washed and imaged.

**(B)** Representative maximum-intensity projections from spinning-disk confocal image stacks showing dextran (cyan) and ASO (magenta) uptake in wild-type U2OS cells, G3BP1/2-knockout cells and knockout cells stably expressing eGFP–G3BP1. The ASO channel is offset by 5 pixels in merged images to aid visualization of colocalization. Scale bar, 10 µm.

**(C)** Percentage of ASO colocalizing with dextran-labeled compartments per cell. Numbers of cells analyzed were 38 wild-type, 59 G3BP1/2-knockout and 64 knockout cells expressing eGFP–G3BP1. Each point represents one cell; black horizontal lines indicate medians. Statistical significance: $p>0.05$ (ns); $p<0.01$ (**); $p<0.001$ (***).

**(D–F)** Number of endolysosomes per cell in the experiments highlighted in **(D)**, dextran uptake **(E)** and ASO uptake **(F)** after the 2-h pulse described in **(A)**. Wild-type cells are shown in gray (38 cells), G3BP1/2-knockout cells in dark gray (59 cells) and knockout cells expressing eGFP–G3BP1 in black (64 cells). Each point represents one cell; black horizontal lines indicate medians. Data are from two biological replicates. Statistical significance: $p>0.05$ (ns); $p<0.05$ (*); $p<0.01$ (**); $p<0.001$ (***).

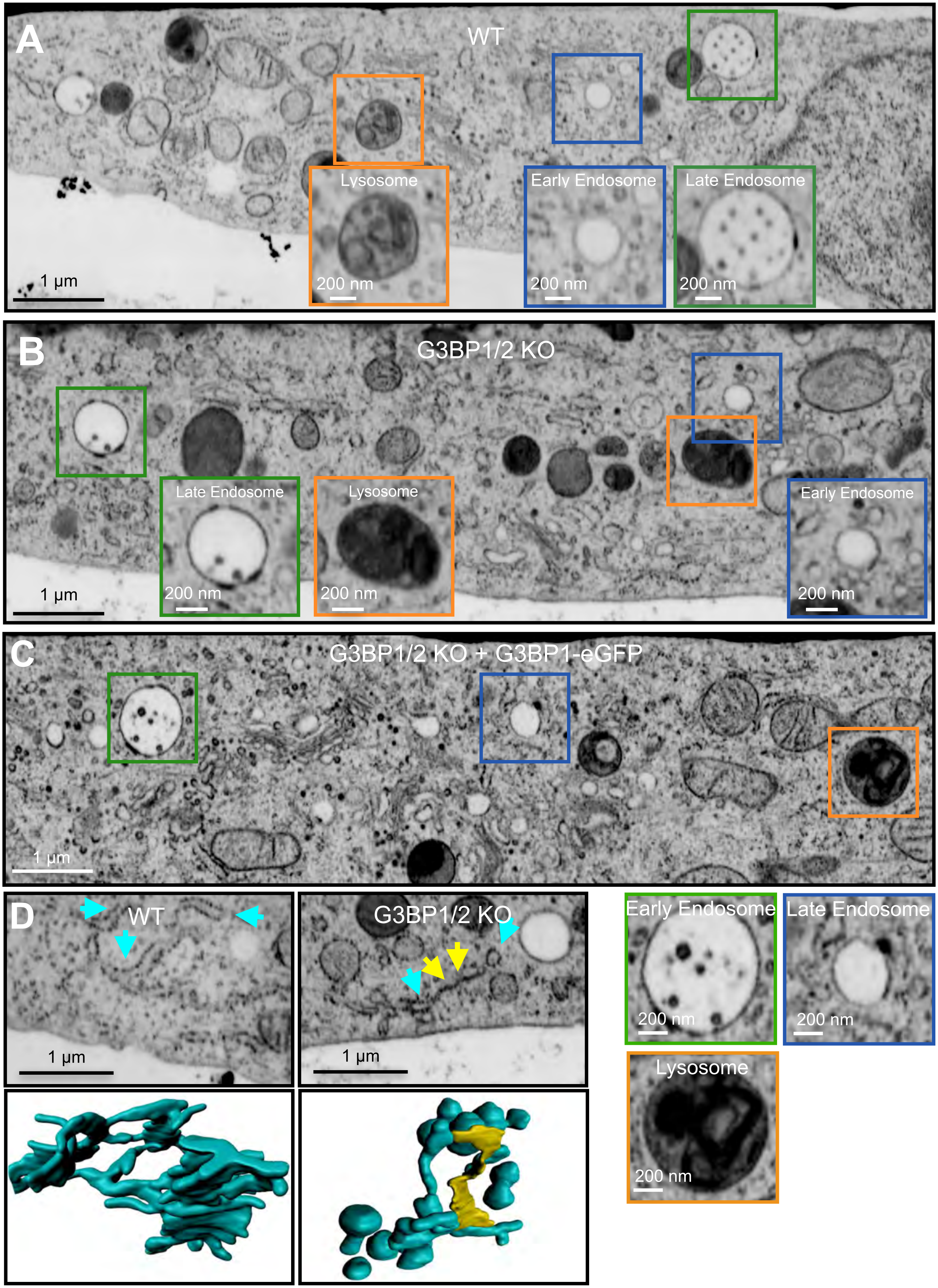


**Supplementary Figure S18**

**Supplementary Figure S18. Related to Figure 8.**

**(A–C)** Representative FIB-SEM images of wild-type U2OS cells (A), G3BP1/2-knockout cells (B) and knockout cells stably expressing eGFP–G3BP1 (C), acquired with 10 × 10 × 10 nm voxels. Insets highlight early endosomes (blue), late endosomes (green) and lysosomes (orange).

**(D)** Representative enlarged views of the ER and corresponding three-dimensional segmentations from wild-type U2OS cells (left) and G3BP1/2-knockout cells (right), sampled from additional regions of the volumes shown in (A) and (B). Blue arrowheads indicate ER sheets bearing ribosomes; yellow arrowheads indicate unusually tight ER sheets lacking detectable bound ribosomes. Scale bars, 1 μm; insets, 200 nm.

#### VIDEOS

**Supplementary Video S1. Related to Figure 1**

Live-cell 3D time-lapse imaging of Alexa Fluor 647-ASO (magenta) trafficking in wild-type U2OS cells with terminal lysosomes labeled by pulse-chase loading of Alexa Fluor 568 dextran (cyan; 4-hours pulse, 20-hours chase). Cells were incubated with fluorescent ASO for 15 min and then chased for 110 min. Live-cell imaging by LLSM was performed during the chase period. The volumetric 3D time-lapse series, acquired every 10 min, is displayed as maximum-intensity projections. Scale bar, 10 μm.

**Supplementary Video S2: Related to Figure 1**

Inset of Supplementary Video 1. Live-cell 3D time-lapse imaging of Alexa Fluor 647-ASO (magenta) trafficking in wild-type U2OS cells with terminal lysosomes labeled by pulse-chase loading of Alexa Fluor 568 dextran (cyan). Cells were incubated with fluorescent ASO for 15 min and then chased for 110 min. The volumetric 3D time-lapse series, acquired every 10 min, is displayed as maximum-intensity projections. Images were shifted by 5 pixels along the x axis to facilitate visualization of colocalization. Scale bar, 10 μm.

**Supplementary Video S3: Related to Figure 3**

Live-cell 3D time-lapse imaging of wild-type U2OS cells stably expressing galectin-3-eGFP, pulsed for 24-hours with Alexa Fluor 568-dextran and Alexa Fluor 647-ASO. Cells were left untreated or treated with 0.5 mM LLOMe for 30 min. Live-cell imaging by LLSM was performed during treatment, and the 3D time-lapse series, acquired every 5 min for 30 min, is displayed as maximum-intensity projections. Arrows indicate cells positive or negative for endolysosomal membrane damage, as assessed by galectin-3 recruitment. Scale bar, 10 μm.

**Supplementary Video S4. Related to Figure 3**

Live cell 3D timelapse imaging of apilimod-treated Alexa Fluor 568-dextran labeled lysosomes (cyan) containing Alexa Fluor 647-ASO (magenta) in wild-type U2OS cells. The white arrow indicates a spontaneous dextran release event accompanied by contraction of the lysosome, while a substantial fraction of the ASO signal remains associated with the lysosome. The time series was acquired by LLSM every 2.3 s and is displayed as maximum-intensity projections. Scale bar, 5 μm.

**Supplementary Video S5: Related to Figure 4**

Live-cell 3D time-lapse imaging of Alexa Fluor 568 dextran-labeled endolysosomes (cyan; outlined by dotted circles) containing ASOs (magenta; arrows) undergoing confined motion (left) or membrane-bound motion (right) in wild-type U2OS cells. The time series was acquired by LLSM every 2.3 s and is displayed as maximum-intensity projections. Scale bar, 1 μm.

**Supplementary Video S6: Related to Figure 8**

Live-cell 3D time-lapse imaging of U2OS cells expressing eGFP-G3BP1 (green), treated with 0.25 mM sodium arsenite alone (Untreated) or in the presence 1 μM ISRIB or 50 μM G3ia. Sodium arsenite was added at time 0. Live-cell imaging by LLSM was performed for 32 min, and the 3D time-lapse series, acquired every 2 min, is displayed as maximum-intensity projections. Scale bar, 10 μm.

**Supplementary Video S7. Related to Figure 8**

Live-cell 3D time-lapse imaging of Alexa Fluor 647-ASO (magenta) trafficking in U2OS G3BP1/2 knockout cells with terminal lysosomes labeled by pulse-chase loading of Alexa Fluor 568 dextran (cyan; 4-hours pulse, 20-hours chase). Cells were incubated with fluorescent ASO for 15 min and then chased for 110 min. The volumetric 3D time-lapse series, acquired every 10 min, is displayed as maximum-intensity projections. Scale bar, 10 μm.

**Supplementary Video S8: Related to Figure 8**

Inset of Supplementary Video 7. Live-cell 3D time-lapse imaging of Alexa Fluor 647-ASO (magenta) trafficking in U2OS G3BP1/2 knockout cells with terminal lysosomes labeled by pulse-chase loading of Alexa Fluor 568 dextran (cyan). Cells were incubated with fluorescent ASO for 15 min and then chased for 110 min. The volumetric 3D time-lapse series, acquired every 10 min, is displayed as maximum-intensity projections. Images were shifted by 5 pixels along the x axis to facilitate visualization of colocalization. Scale bar, 10 μm.
